# A modular toolkit for environmental *Rhodococcus, Gordonia*, and *Nocardia* enables complex metabolic manipulation

**DOI:** 10.1101/2024.02.21.581484

**Authors:** Zachary Jansen, Abdulaziz Alameri, Qiyao Wei, Devon L. Kulhanek, Andrew R. Gilmour, Sean Halper, Nathan D. Schwalm, Ross Thyer

## Abstract

Soil-dwelling Actinomycetes are a diverse and ubiquitous component of the global microbiome, but largely lack genetic tools comparable to those available in model species such as *E. coli* or *Pseudomonas putida*, posing a fundamental barrier to their characterization and utilization as hosts for biotechnology. To address this, we have developed a modular plasmid assembly framework along with a series of genetic control elements for the previously genetically intractable Gram-positive environmental isolate *Rhodococcus ruber* C208 and demonstrate conserved functionality in diverse environmental isolates of *Rhodococcus, Nocardia* and *Gordonia*. This toolkit encompasses Mycobacteriale origins of replication, broad-host range antibiotic resistance markers, transcriptional and translational control elements, fluorescent reporters, a tetracycline-inducible system, and a counter-selectable marker. We use this toolkit to interrogate the carotenoid biosynthesis pathway in *Rhodococcus erythropolis* N9T-4, a weakly carotenogenic environmental isolate and engineer higher pathway flux towards the keto-carotenoid canthaxanthin. This work establishes several new genetic tools for environmental Mycobacteriales and provides a synthetic biology framework to support the design of complex genetic circuits in these species.

**IMPORTANCE:** Soil-dwelling Actinomycetes, particularly the Mycobacteriales, include both diverse new hosts for sustainable biomanufacturing and emerging opportunistic pathogens. *Rhodococcus, Gordonia* and *Nocardia* are three abundant genera with particularly flexible metabolisms and untapped potential for natural product discovery. Among these, *Rhodococcus ruber* C208 was shown to degrade polyethylene, *Gordonia paraffinivorans* can assimilate carbon from solid hydrocarbons, and *Nocardia neocaledoniensis* (and many other *Nocardia*) possesses dual isoprenoid biosynthesis pathways. Many species accumulate high levels of carotenoid pigments, indicative of highly active isoprenoid biosynthesis pathways which may be harnessed for fermentation of terpenes and other commodity isoprenoids. Modular genetic toolkits have proven valuable for both fundamental and applied research in model organisms, but such tools are lacking for most Actinomycetes. Our suite of genetic tools and DNA assembly framework were developed for broad functionality and to facilitate rapid prototyping of genetic constructs in these organisms.

## INTRODUCTION

Actinomycetota are a large phylum of Gram-positive bacteria characterized by a high GC content(1). While some genera are relatively well-studied, such as *Streptomyces* for antibiotic and natural product discovery, *Corynebacterium* as an established fermentation host, and *Mycobacterium* as a human and livestock pathogen(2–5), most are poorly characterized (6–8). Among these, *Rhodococcus, Nocardia*, and *Gordonia* are three genera that contain numerous species with industrially relevant traits including natural product production, degradation of anthropogenic pollutants, and diverse carbon and nitrogen assimilation pathways that enable oligotrophic growth (8–15). Members of the genus *Rhodococcus*, the best characterized of the three, display an especially broad range of phenotypes. *R. ruber* C208 and *R. erythropolis* DCL14 can degrade solid polyolefins and paraffin wax, respectively(16–18). *R. opacus* MR11 encodes a functional Calvin-Benson-Bassham cycle and a hydrogenase cluster that enables fully autotrophic growth(19), and several plant-associated *R. qingshengii* isolates either produce plant growth regulators or fix inorganic nitrogen(20, 21). Importantly, these genera are easily maintained in the laboratory under aerobic and mesophilic conditions, and they do not require specialized growth media nor handling precautions.

A significant barrier to investigating the fundamental biology of these species and their application as potential industrial hosts is a lack of dedicated genetic tools. Molecular biology in model prokaryotes employs standardized genetic tools, which recently have been adapted for efficient, modular assembly to enable rapid construction of plasmids and gene circuits (22, 23). Commonly, this is accomplished by using Golden Gate assembly, which relies on Type IIs restriction enzymes to generate a series of user-specified cloning overhangs. Individual genetic elements are flanked by different overhangs that set the assembly order and form the basis of a Modular Cloning (MoClo) system. In MoClo systems, different parts with identical cloning overhangs are seamlessly interchangeable. The development of these toolkits for non-model microorganisms can significantly ‘close the gap ‘ in terms of genetic tractability and accessibility for researchers(24–28). While sets of genetic parts have been characterized in some *Rhodococcus* species(29, 30), the development of genetic tools for *Nocardia* and *Gordonia* is lacking. Furthermore, within the genus *Rhodococcus*, work largely has been restricted to a small number of genetically tractable species, most often *Rhodococcus erythropolis* and *R. opacus*. Less genetically tractable species, commonly *R. ruber* or *R. rhodochrous*, are sometimes compared as ‘outgroup ‘ *Rhodococcus* and generally exhibit poor transformation efficiency (31– 33). Furthermore, efforts to standardize plasmid and circuit construction in these species are in their infancy, and they typically rely on *ad hoc* DNA assembly methods(34–36).

Several bacterial and eukaryotic toolkits are now available to researchers with varying assembly architecture but offering a shared set of advantages. Chief among these are the abilities to rapidly construct plasmids and to prototype new genetic designs in parallel. Plasmids are constructed with reusable, modular DNA fragments that do not rely on bespoke primers for each unique connection, phenotypic dropout markers that facilitate identification of correct plasmid constructs, and a simplified quality assurance workflow that reduces the need for DNA sequencing. Transcriptional units with variable expression strengths easily can be prototyped in parallel using libraries of predictable and modular transcriptional and translational control elements. Additional advantages include the use of standardized assembly protocols, genetic elements to reduce variability between different experiments or users, and a large body of documentation and introductory resources for new users(37).

MoClo systems are typically built around a framework of origins of replication and selectable markers previously demonstrated to function in the target species. These are used to assemble the plasmid backbones which can accommodate new transcriptional units or other DNA parts for evaluation. For Gram-negative bacteria, there are several broad host range (BHR) plasmid origins of replication that function in a wide range of species. In contrast, the host range of most origins of replication for Gram positive bacteria remains poorly defined. This presents a significant challenge when designing MoClo systems for poorly studied Gram-positive bacteria, and it necessitates additional investment to characterize and reformat potentially large numbers of genetic elements. To maintain the advantages inherent to modular cloning for evaluation of core elements of plasmid backbones, such as origins of replication and selectable markers, we propose that a simplified plasmid assembly hierarchy has utility.

In this work, we describe the development of RhoClo, a modular DNA assembly framework consisting of two complementary and interconnected assembly hierarchies that enable the construction of simplified plasmid backbones and genetic circuits built from transcriptional units, respectively, in non-model Mycobacteriales. Plasmids can be constructed using combinations of five different Mycobacteriale origins of replication and five broad-host-range antibiotic resistance markers, which we have evaluated in 17 species of *Rhodococcus, Nocardia*, and *Gordonia*. Plasmids support the construction of multi-component genetic circuits, and a range of new, cross-species transcriptional and translational control elements afford fine control of gene expression. We anticipate this design framework will facilitate the characterization and development of new genetic components for non-model Mycobacteriales and show proof-of-principle experiments including the construction of novel counter-selectable markers and genetic circuits to perturb carotenoid pigment production in the environmental isolate *R. erythropolis* N9T-4.

## RESULTS

### Shuttle plasmid construction

The RhoClo genetic toolkit features two complementary and interconnected assembly hierarchies: plasmid backbone assembly and genetic circuit assembly (**Figure 1**). In both hierarchies, individual genetic elements are deconstructed into minimal units termed ‘parts ‘, which are flanked by type IIs restriction sites and archived on a simple plasmid for storage or propagation in *E. coli*. In the construction of a shuttle plasmid backbone, five different parts flanked by AarI restriction sites are used (**Figure 1a-b**). Those parts are a cargo site containing a fluorescent dropout reporter (i-ii), a Mycobacteriale origin of replication (denoted AC, ii-iii), a broad host range selectable marker (iii-iv), an accessory transcriptional unit or vacant spacer sequence (iv-v), and an *E. coli* origin of replication (denoted EC, v-vi), part overhangs are indicated in parentheses. This initial plasmid construction step provides a streamlined system for rapid assembly of shuttle plasmids using the minimum number of parts to aid in characterizing new origins of replication or selection markers. The restriction enzyme AarI was selected for this system due to its extended recognition sequence, which reduces the need to domesticate large DNA sequences upfront. Only once functional replicons or markers have been identified do researchers need to invest in removal of the more abundant BsaI and Esp3I restriction sites, which are used in the second assembly hierarchy. A fluorescent cloning marker was selected to facilitate plasmid assembly and subsequent cargo insertion in non-typical *E. coli* hosts, such as methylation deficient or λpir+ strains(37).

**Figure 1:**
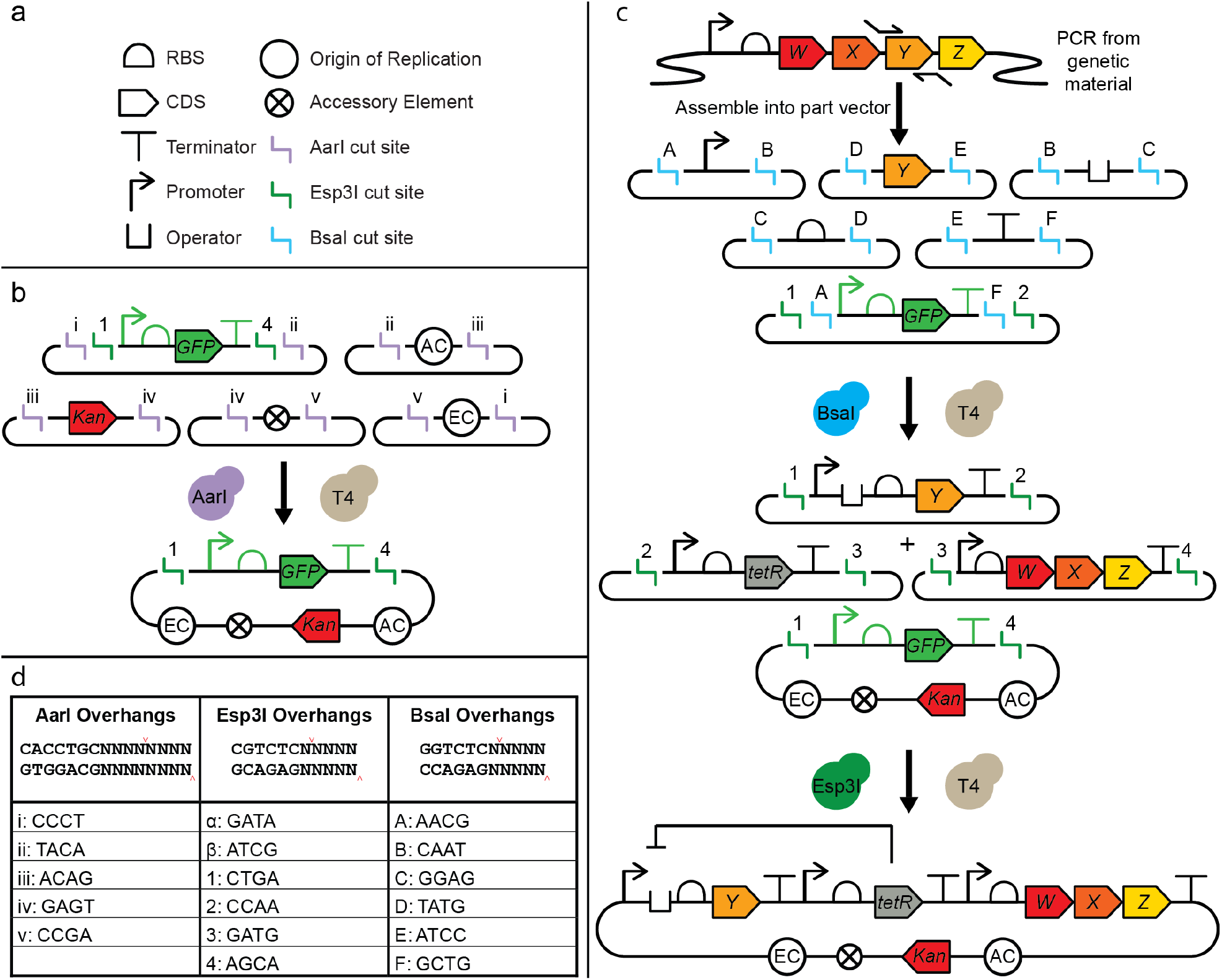
Overview and cartoon schematic of the RhoClo modular cloning architecture. (a) SBOL symbols used to represent each of the genetic elements. (b) Genetic elements of plasmids are broken down into single parts and separated onto part vectors in hierarchy one for shuttle vector assembly. These part plasmids can be assembled into modular backbone plasmids in a standard Golden Gate reaction using the type IIs restriction enzyme AarI. Assembled plasmids containing a fluorescent dropout reporter are then used as backbones for assembly of transcriptional units. (c) Assembly of transcriptional unit and multiple transcriptional unit plasmids starting from individual Part plasmids. Part plasmids can be combined with other Part plasmids spanning all necessary elements of a transcriptional unit and assembled into a backbone plasmid in a single Golden Gate reaction using the type IIs restriction enzyme BsaI. Multiple transcriptional unit plasmids can be concatenated in a second reaction using the type IIs restriction enzyme Esp3I. (d) The recognition and cut sites of each restriction enzyme used and four bp cloning overhangs used set the assembly order of the individual DNA parts.

### Characterization of Mycobacteriale origins of replication

We hypothesized that using a *Rhodococcus* strain with low genetic tractability (*R. ruber* C208) as the host for toolkit development would increase the chance of compatibility in additional *Rhodococcus* species with varying degrees of genetic tractability and other closely related genera such as *Nocardia* and *Gordonia*. Twenty Actinomycete origins of replication were evaluated, nineteen of which were originally isolated from Mycobacteriales and one from *Arthrobacter rhombi* (Micrococcales) (**Table A1**). Template DNA was obtained from either public plasmid repositories or via direct synthesis. Synthesis of most DNA sequences was challenging due to the high GC content and presence of complex secondary structure, and minor sequence optimization was required in some cases. DNA sequence changes were restricted to synonymous codons within open reading frames to minimize the disruption of cryptic regulatory elements. All DNA sequences are provided in **Table A2**. DNA fragments encoding the origins of replication were assembled with a GFP cassette under the control of a constitutive *E. coli* promoter, an *E. coli* pUC origin of replication, and the APH(3 ‘)-IIa kanamycin marker previously reported to work in several *Nocardia* and *Rhodococcus* species(38, 39).

For each plasmid, *R. ruber* cells were transformed by electroporation and kanamycin resistant transformants were recovered. Three transformants were inoculated in TSB medium supplemented with kanamycin and cultured three times to confluence. Plasmid DNA was then isolated from *R. ruber* and transformed back into *E. coli* to confirm recovery of GFP+ transformants. Six origins met this threshold: pNC903, pRET1100, pNG2, pSOX, pNC500, and pYS1, with a seventh (pXT107) yielding a mix of GFP+ and GFP negative *E. coli* transformants. Analysis of GFP-pXT107 transformants revealed a truncation of the plasmid which largely preserved the pXT107 origin of replication, kanamycin marker, and an *E. coli* pUC origin of replication, but eliminated the entirety of the GFP dropout. For all origins of replication, remaining type IIs restriction sites were removed from the origin fragments, and in all but one case (pSOX), the sites were located within open reading frames, which allowed for synonymous codon replacements. The pSOX origin contained a BsaI site within a conserved structural region and a small 6 nt library spanning the site was screened to identify clones capable of replication. Despite several attempts we were unable to fully domesticate the pYS1 origin. pXT107 and pYS1 were not evaluated further, but they are promising candidates for further engineering efforts in Mycobacteriales.

### Mycobacteriale selectable markers

While many Mycobacteriales are sensitive to kanamycin, *Nocardia* species are commonly resistant to this antibiotic(40). To broaden the utility of our cloning framework, the pNC903 origin, which has been reported to work in diverse Mycobacteriales, was reassembled with a *Rhodococcus*-compatible E2-Crimson expression cassette, and either the kanamycin resistance marker or one of four additional selectable markers; apramycin (Apr, *aac(3)-IVa*), spectinomycin (Spec, *aadA1*), zeocin (Zeo, *Sh ble*), or nourseothricin (Ntc, *nat1*). The four additional markers were expressed using the strong constitutive P-45 promoter from the Actinomycete *Corynebacterium glutamicum*, and the *Sh ble* and *nat1* genes were codon optimized to increase their GC content. To evaluate performance of the new markers, plasmids were transformed into *R. ruber, G. paraffinivorans*, and *N. jiangsuensis* (**Figure 2a, Table A3**). All markers conferred resistance in the three strains tested.

**Figure 2:**
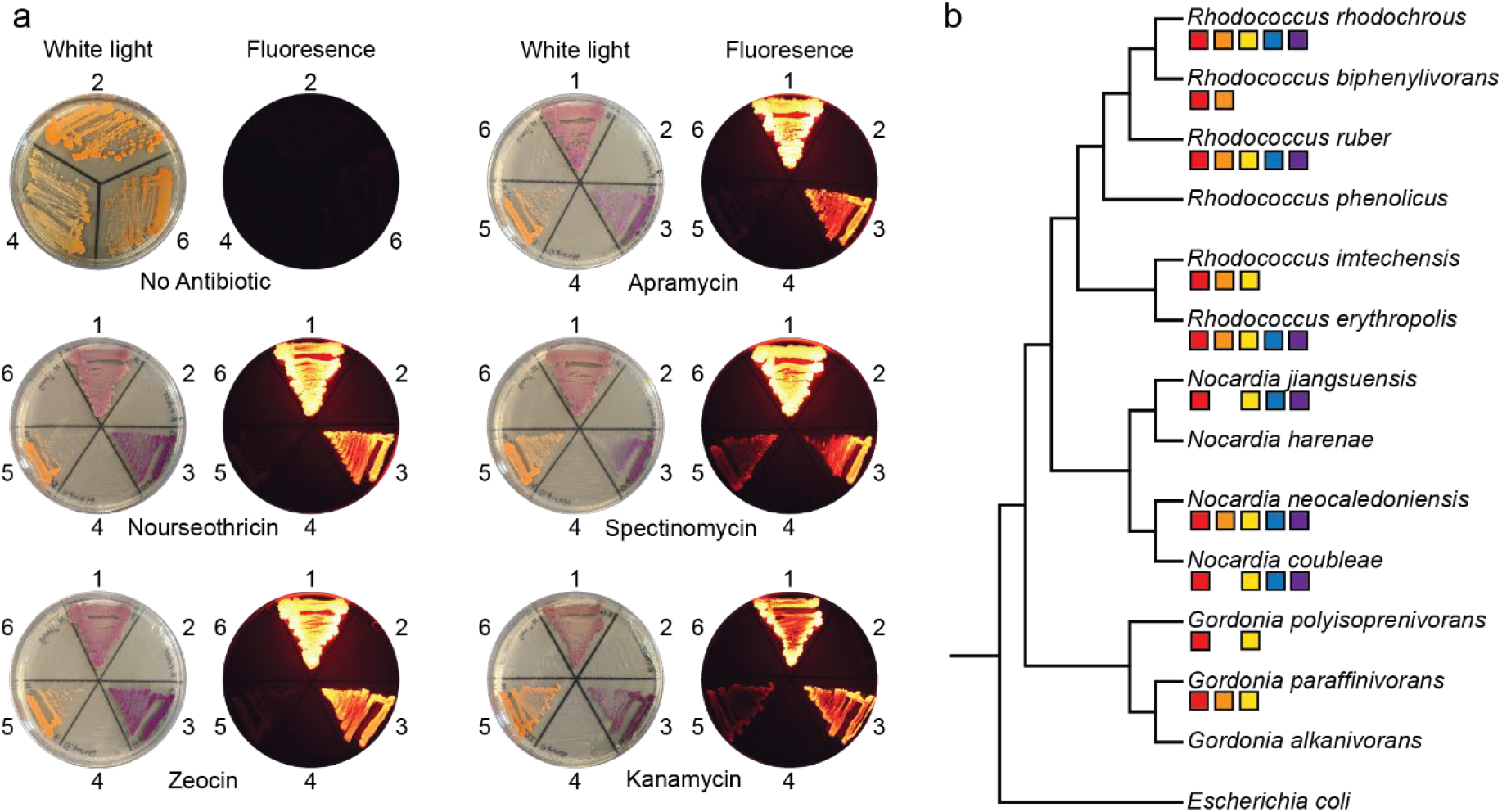
Screening selectable markers and plasmid origins of replication (a) The five different resistance markers confer resistance to at least 100 μg mL^-1^ of each antibiotic in the three strains tested: *R. ruber, G. paraffinivorans*, and *N. jiangsuensis*. Three strains, each with and without a plasmid encoding the different selectable markers and an expression cassette for E2-Crimson were plated on each of the five antibiotics. (1) *R. ruber* + plasmid, (2) *R. ruber* cells only, (3) *G. paraffinivorans* + plasmid, (4) *G. paraffinivorans* cells only, (5) *N. jiangsuensis* + plasmid, and (6) *N. jiangsuensis* cells only. (b) A phylogenetic tree of the species tested showing compatibility with the different plasmid origins of replication. Red squares indicate pNC903, orange squares indicate pRET1100, yellow squares indicate pNG2, blue squares indicate pNC500, and purple squares indicate pSOX.

To further establish the broad compatibility of the shuttle plasmids, all five domesticated origins were assembled individually with the selection markers for the two most broadly effective antibiotics (Apr and Ntc) and the E2-Crimson expression cassette and transformed into an additional 16 species spanning the genera *Rhodococcus, Nocardia, Gordonia*, and *Dietzia* (**Table A4-5**). The pNC903 origin had the broadest observed host range, yielding E2-Crimson positive transformants in ten different species (**Figure 2b**). As expected, all five origins supported replication in *R. erythropolis*, the most genetically tractable *Rhodococcus* species. No transformants were recovered for any of the *Dietzia* species except for *D. dagingensis*, which yielded a small number (<5) of confirmed transformants with pNC903. These results are largely consistent with previous efforts to evaluate the host range of pNC903 and pRET1100, although it should be noted that a different transformation protocol was used than in previous efforts, and no efforts were made to optimize the transformation protocol for different species(41, 42).

### Transcriptional unit assembly

A second assembly hierarchy is used for construction and concatenation of individual transcriptional units. Transcriptional units are assembled from five basic genetic elements: promoters, operators, translational control elements, coding sequences, and transcriptional terminators. Each genetic element is first constructed as a discrete genetic part of a high copy number plasmid with flanking BsaI restriction sites that generate the corresponding assembly overhangs (**Figure 1c-d**). A schematic for the construction of Part plasmids is provided in the Appendix (**Figure A1**). Additional subparts with unique assembly overhangs can be added as required. To assemble a single transcriptional unit, these five parts are combined with a Transcriptional Unit (TU) plasmid (TU-X-DO) containing a fluorescent cloning marker using a Golden Gate reaction mediated by BsaI. Individual TU plasmids set the position of the transcriptional unit in any further assemblies denoted by X. To preserve compatibility with the widely used Yeast Toolkit (YTK), the three genetic elements which comprise the 5′-UTR are collectively flanked by AACG and TATG assembly overhangs which correspond to a Type 2 part(22). Similarly, the coding sequence and transcriptional terminator parts use the same assembly overhangs as YTK Type 3 and Type 4 parts respectively. Colonies with the correct phenotype can easily be identified by loss of the fluorescent cloning marker and following recovery of plasmid DNA may be digested with restriction enzymes to confirm the correct product size. All TU plasmids contain two EcoRI sites outside of the fluorescent cloning marker which can be used to quickly validate assemblies.

Single transcriptional units can be concatenated in a similar process where a series of TU plasmids now take the place of individual genetic parts, and a Multi-Transcriptional Unit (MTU) plasmid serves as the assembly scaffold. This reaction is mediated by a second Golden Gate reaction using the enzyme Esp3I. These plasmids are suited to the construction of larger genetic circuits, for example, allowing for more complex regulation using transcription factors or assembling multi-enzyme biosynthetic pathways. The unique assembly overhangs and a schematic of this process are shown in **Figure 1c-d**.

### Modular genetic control elements

Synthetic transcriptional and translational control elements that result in predictable levels of protein expression are essential genetic elements of a modular toolkit and enable the construction of finely tuned genetic circuits(43). To build a series of transcriptional control elements, we selected the P*amiM* promoter as our starting point, a strong constitutive promoter previously engineered from the *Rhodococcus ruber* P*ami* promoter(44). This promoter was initially adapted to our modular cloning framework by converting the last four bases of the -10 region from AAAT to CAAT (**Figure 3a**). This mutation was well-tolerated and the newly designated P*amiM* T2A promoter maintained strong transcriptional activity. This design allows similarly modular operator sites for transcription factors to be appended directly adjacent to the - 10 region of the promoter and enables any constitutive promoter to function as an inducible promoter if desired. To generate a genetic reporter suitable for testing a library of promoter variants, the P*amiM* T2A promoter was assembled with a modular TetO operator and a pre-existing bicistronic translational control element upstream of the coding sequence for E2-Crimson (**Figure 3b**). Inclusion of a TetO operator paired with assembly of plasmids in the *E*.

**Figure 3:**
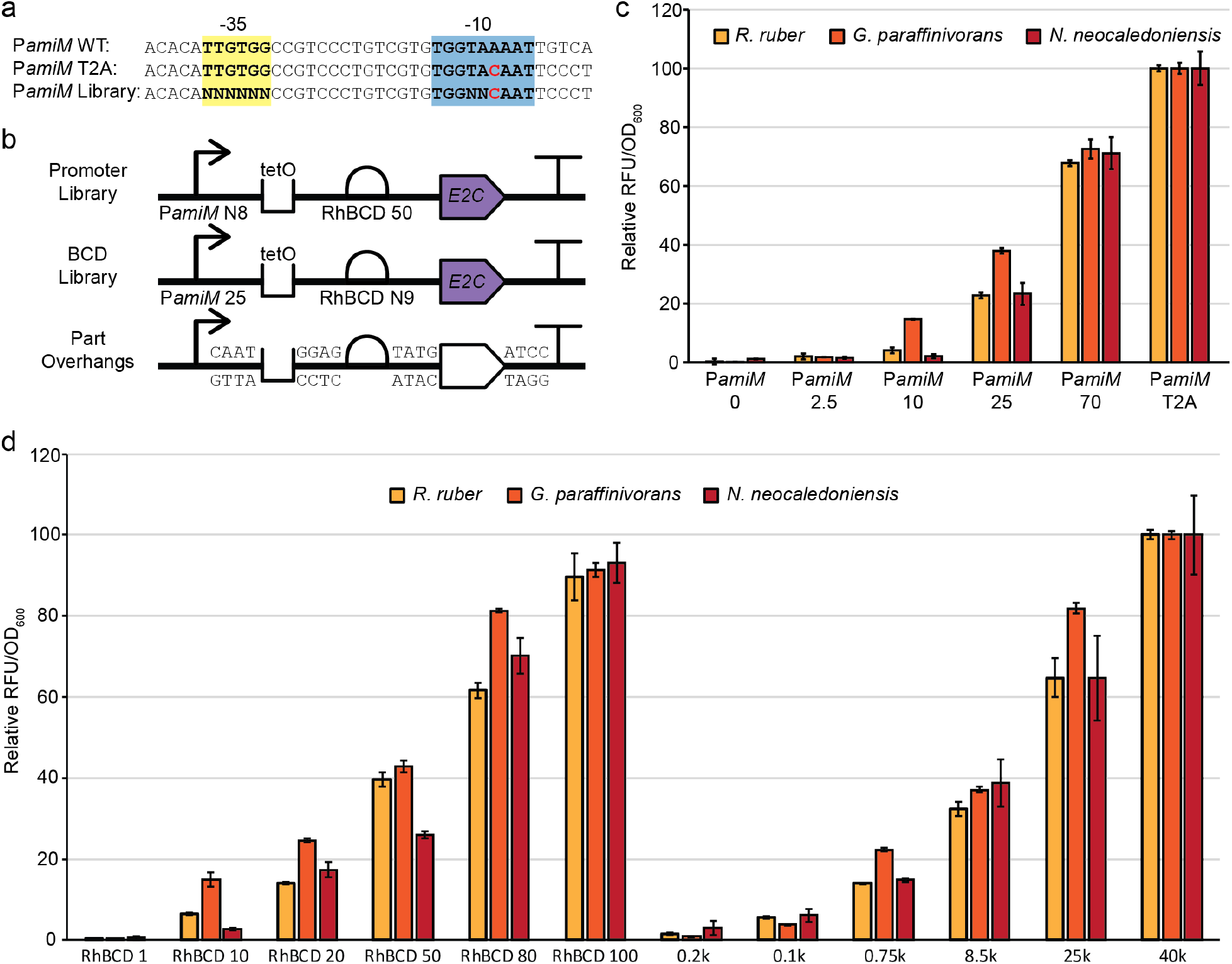
Modular series of promoters and BCD translational control elements. (a) The sequence of the -35 and -10 sites in the P*amiM* promoter and the immediately surrounding context as it was adapted for the modular cloning toolkit. (b) The transcriptional units used to select the libraries for promoters and BCDs respectively. The third schematic shows the four base pair overhangs used in between each genetic part. (c) The relative strength of the six promoters normalized across the three species such that the strongest promoter has a relative activity of 100. (d) The relative strength of 11 BCD translational control elements. The relative strength of the two BCD series is normalized across three species such that the strongest BCD has a relative activity of 100.

*coli* strain Marionette-Clo DH10B, which encodes the TetR repressor, silences the reporter to prevent any toxicity during cloning(45).

A library of the P*amiM* T2A promoter was generated by randomizing eight nucleotides spanning the entire six bp -35 region and two bp in the -10 region using degenerate oligonucleotide primers (**Figure 3a**). The library was transformed into *R. ruber*, and individual colonies spanning a range of E2-Crimson expression levels were selected. Plasmids encoding promoter variants were isolated and sequenced, followed by reconstruction of chosen promoter variants and re-phenotyping in *R. ruber*. Variants spanning nearly two orders of magnitude of transcriptional activity were identified, then validated in *G. paraffinivorans* and *Nocardia neocaledoniensis*, and found to maintain their relative rank order across the three genera (**Figure 3c, Table A6**).

We previously published two series of Bi-Cistronic Design translational control elements (BCDs), one designed for *E. coli* and one designed for *Rhodococcus*(46). We also observed that the series of 12 BCDs designed for *E. coli* are functional in *R. ruber* C208, although they do not maintain the same rank order. The series of BCDs developed for *Rhodococcus* was evaluated in *G. paraffinivorans* and *N. neocaledoniensis* and found to maintain the same relative expression strength across all three genera. Absolute fluorescence varied greatly between the strains with much lower fluorescence observed in *N. neocaledoniensis*. A curated set of five BCDs from the series previously designed for *E. coli* evenly spaced across the range of translational activity in *R. ruber* was also evaluated in *G. paraffinivorans*, and *N. neocaledoniensis* and found to maintain their relative strengths. To extend the range of the BCD series, an additional BCD was designed by transferring the strongest 9 nt Shine Dalgarno (SD) site from the BCD context developed for *Rhodococcus* (AGAAGGAGA) into the BCD context developed for *E. coli* in the corresponding SD 2 position. This BCD was designated 40k and was observed to have the highest strength of any of the translational control elements tested among the three species. Conversely, the sequence of the weakest SD site from the series developed in *E. coli* (BCD 0.2k, TCACGTCCC) was transferred into the BCD context developed for *Rhodococcus* to expand the series and was designated RhBCD 1 (**Figure 3d, Tables A7-8**). Data that is not normalized across strains is provided in **Figure A2**.

Inducing transcription via protein regulators in response to small molecules or other stimuli are key biological strategies that facilitate dynamic control over the amount of protein in a cell(47). Ideal inducible systems should have minimal interplay with the native function of the cell, activate transcription over a wide range of inducer concentrations (dynamic range), and lead to a significant change in gene expression (fold-induction). To achieve these latter functions, inducible systems should be optimized to result in low transcription when fully repressed (or otherwise inactive) and achieve maximal expression when fully induced(48). To construct a model inducible system, we assembled a plasmid containing two transcriptional units that place E2-Crimson under the control of a modular promoter and TetO operator, and constitutively express TetR, the standard tetracycline-responsive transcriptional repressor (**Figure 4a**). The performance of this circuit in *R. ruber* was evaluated by measuring E2-Crimson induction in response to a gradient of anhydrous tetracycline (aTc) concentrations, a potent inducer of TetR (**Figure 4b**). The results show near zero signal when the plasmid is fully repressed, and a maximum induction of 55-fold at 2000 ng mL^-1^ aTc. The induction curve for this simple circuit spans a large range of aTc concentrations, providing the opportunity for precise control over the level of induction. Several iterative designs were used to optimize the performance of this circuit, making use of the different translational and transcriptional control elements, and a detailed discussion is provided in the Appendix (**Supplementary Results, Figure A3**).

**Figure 4:**
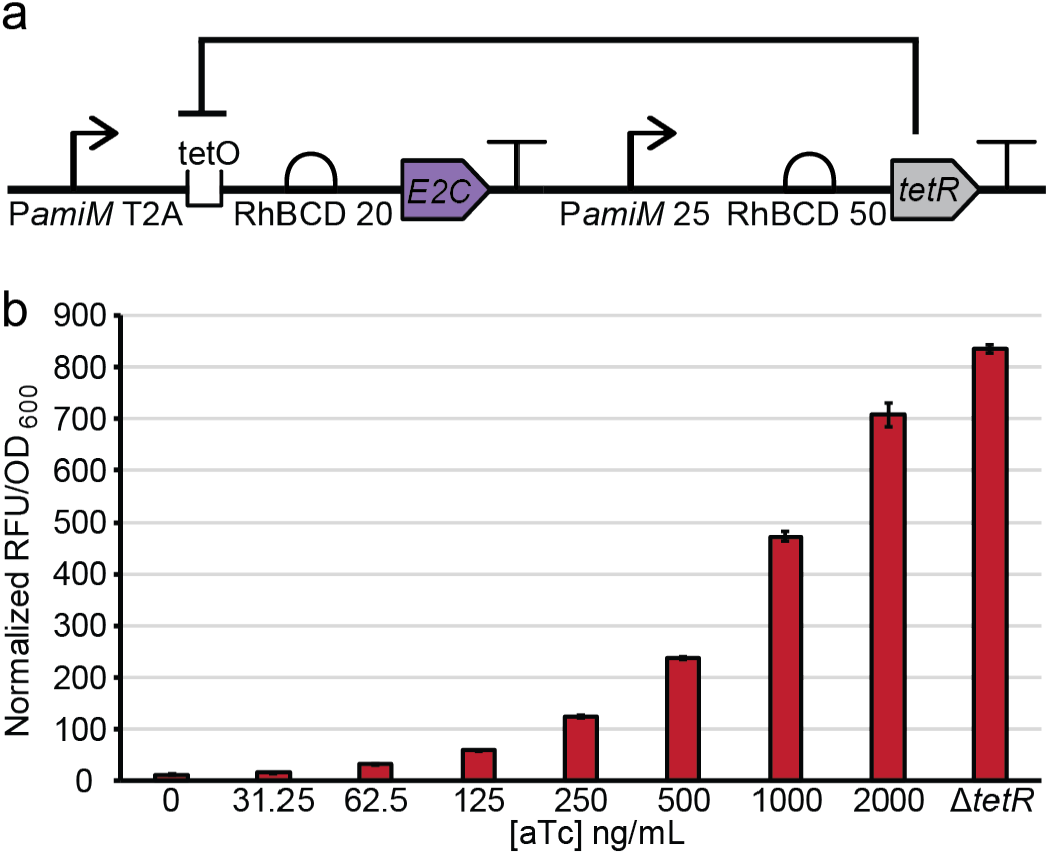
A TetR inducible system in *R. ruber* C208. (a) The selected plasmid design for the tetracycline inducible system with the chosen promoters and BCDs labeled. (b) The fluorescent signal from *R. ruber* cultures induced with a gradient of different concentrations of anhydrous tetracycline (aTc).

### Fluorescent and counter-selectable markers

Fluorescent proteins are essential tools for modern molecular biology that enable a variety of quantitative measurements in living cells. We evaluated several different fluorescent proteins in *R. ruber* and identified four that express well using strong constitutive promoters, resulting in a visible color change (**Figure A4**). Three of these fluorescent proteins, E2-Crimson, mScarlet-I, and mCherry have emission maxima in the red region between 600 nm and 650 nm(49–51). This is a desirable phenotype because the background fluorescence of most bacterial cells in this region is close to zero resulting in a high signal to noise ratio. Additionally, E2-Crimson is a native tetramer reported to have low cellular toxicity, while mScarlet-I and mCherry are monomeric and useful for applications such as tethering or protein fragment complementation assays. A fourth fluorescent protein, BFP-Bluebonnet, with an emission maximum of 464 nm also proved effective in *R. ruber* and possesses excitation and emission spectra orthogonal to the three red fluorescent proteins enabling multi-channel fluorescence imaging(52). Interestingly, derivatives of *Av*GFP (GFP+, sfGFP, mClover3, and sfTq2ox) and two genetically distinct green fluorescent proteins, ZsGreen and mNeonGreen, performed poorly and yielded few fluorescent transformants, possibly indicating toxicity during folding or maturation in *R. ruber* (**Table A9**).

Counter-selectable markers are powerful genetic tools that are commonly used to cure plasmids from living cells and perform scarless genomic integrations. Previously, a counter-selectable marker utilizing the *codA::upp* gene fusion was developed for use in *Rhodococcus equi*. Cytosine deaminase (CD), encoded by *codA*, converts the cytosine analog 5-fluorocytosine (5-FC) into the toxic pyrimidine nucleoside analog 5-fluorouracil (5-FU), which is further converted by uracil phosphoribosyltransferase (UPRT, encoded by *upp*) into 5-fluoro-dUMP, a potent inhibitor of DNA and RNA synthesis (**Figure 5a**). Previous work suggests that *Rhodococcus* and many other genera of Mycobacteriales, with the notable exception of *Nocardia*, are broadly resistant to 5-FC and highly sensitive to 5-FU, and that most Actinomycetes encode a *upp* gene(53).

**Figure 5:**
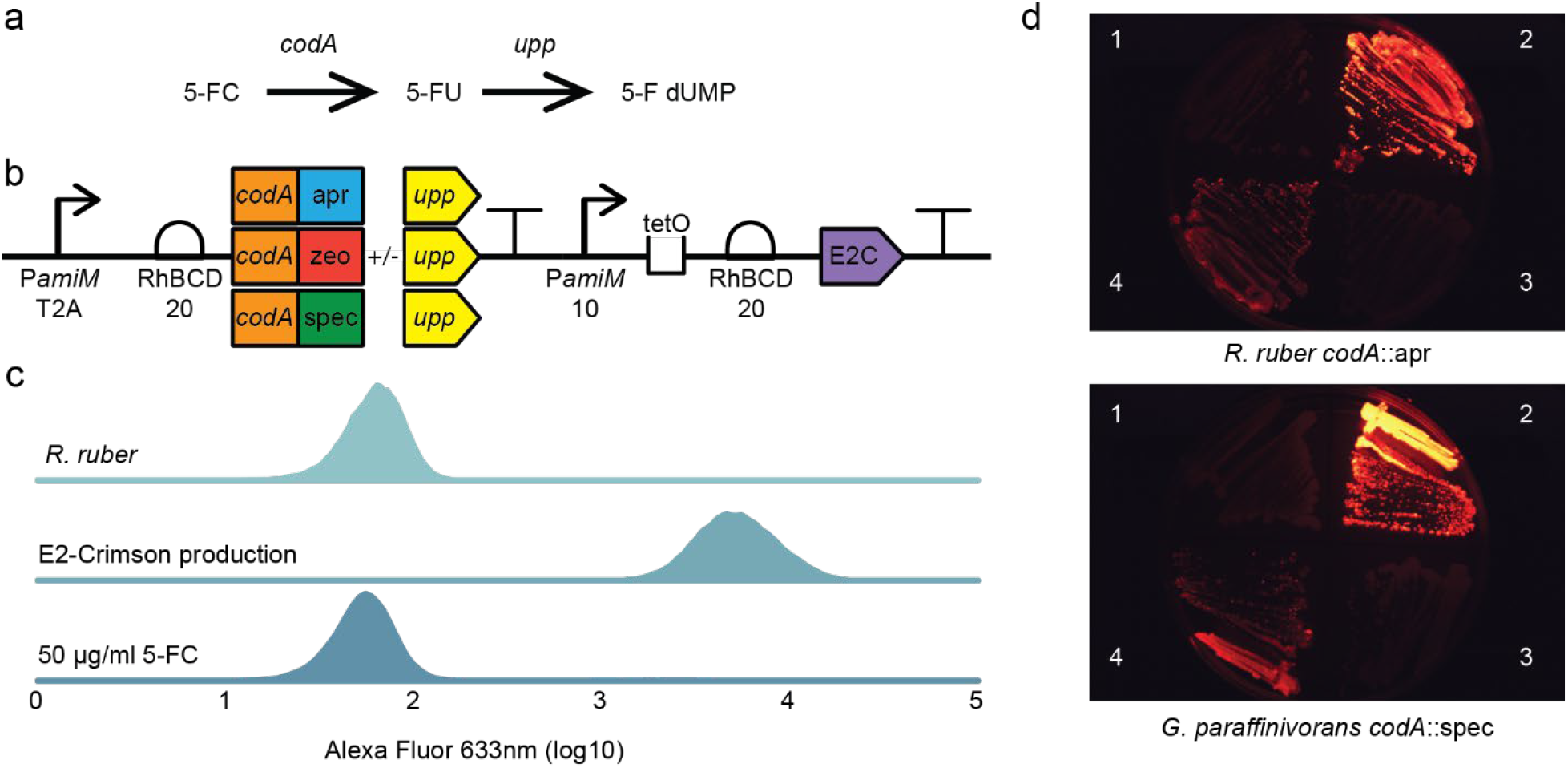
A modular counter-selectable marker can be used to rapidly cure plasmids in Mycobacteriales. (a) The pathway by which 5-FC is converted into the toxic compound 5-fluoro-dUMP. (b) The layout of plasmid variants used for counter-selection with the promoters and BCDs labeled. (c) Cytometry data showing the red fluorescence detected in three different cell populations. Cells are either non-transformed *R. ruber* cells (top), cells transformed with the *codA::zeo::upp* reporter but not treated with 5-FC (middle), and the transformed cells treated with 50 μg mL^-1^ 5-FC. (d) Agar plates showing successful *R. ruber* and *G. paraffinivorans* plasmid curing using the *codA::*apr or *codA::*spec reporters. (1) no plasmid, (2) transformants under selection with antibiotic, (3) transformant following counter-selection with 50 μg mL^-1^ 5-FC, and (4) transformant passaged in TSB without antibiotic selection.

To improve the utility of this reporter, we designed a new series of translational fusions which link the *codA* gene to one of the *Sh ble, aac(3)-IVa*, or *aadA1* genes, which encode resistance to zeocin, apramycin, and spectinomycin respectively, via a short G4SA linker. The *upp* gene can optionally be linked to the 3 ‘ end of this fusion when needed without impairing function of the resistance markers. Plasmids were assembled containing the pNC903 origin, an E2-Crimson expression cassette, and all three of these gene fusions in place of a standalone resistance marker (**Figure 5b**) and transformed into *R. ruber* and *G. paraffinivorans*. Each gene fusion conferred the expected antibiotic resistance and sensitivity to 5-FC in both species. Flow cytometry of *R. ruber* cell populations initially containing the *codA::Sh ble:upp* gene fusion indicated complete loss of the plasmid following growth for two days in the presence of 50 μg mL^-1^ 5-FC and absence of antibiotic selection (**Figure 5c**). Selection and counter-selection with each of the three markers were evaluated in both strains, and all showed complete plasmid loss after one 5-FC treatment (**Figure 5d, Figure A5**).

### Probing the isoprenoid biosynthesis pathway in *Rhodococcus*

Many *Rhodococcus, Gordonia*, and *Nocardia* species natively biosynthesize high levels of carotenoid pigments, even under mild, mesophilic conditions(54). While carotenoids are valuable specialty chemicals, their biosynthetic precursors are common to all isoprenoids, a diverse family of secondary metabolites with many commercial applications. Interestingly, members of these genera include some of the few bacteria that encode both known pathways for biosynthesizing the isoprenoid precursors, the methylerythritol phosphate (MEP) pathway and the mevalonate (MVA) pathway, making them promising new hosts for isoprenoid biosynthesis at scale. As a proof-of-principle experiment to show control over isoprenoid biosynthesis, we selected three key steps to control the flux through the carotenoid biosynthesis pathway: *dxs*, which catalyzes the first step of the MEP pathway and has been shown to have a strong impact on overall isoprenoid biosynthesis, the *crtEYIB* operon, which converts farnesyl pyrophosphate (FPP), the last universal isoprenoid precursor, into the orange colored carotenoid β-carotene in four steps, and *crtO*, which encodes β-carotene ketolase and converts β-carotene stepwise into echinenone and finally canthaxanthin, a red ketocarotenoid (**Figure 6a**).

**Figure 6:**
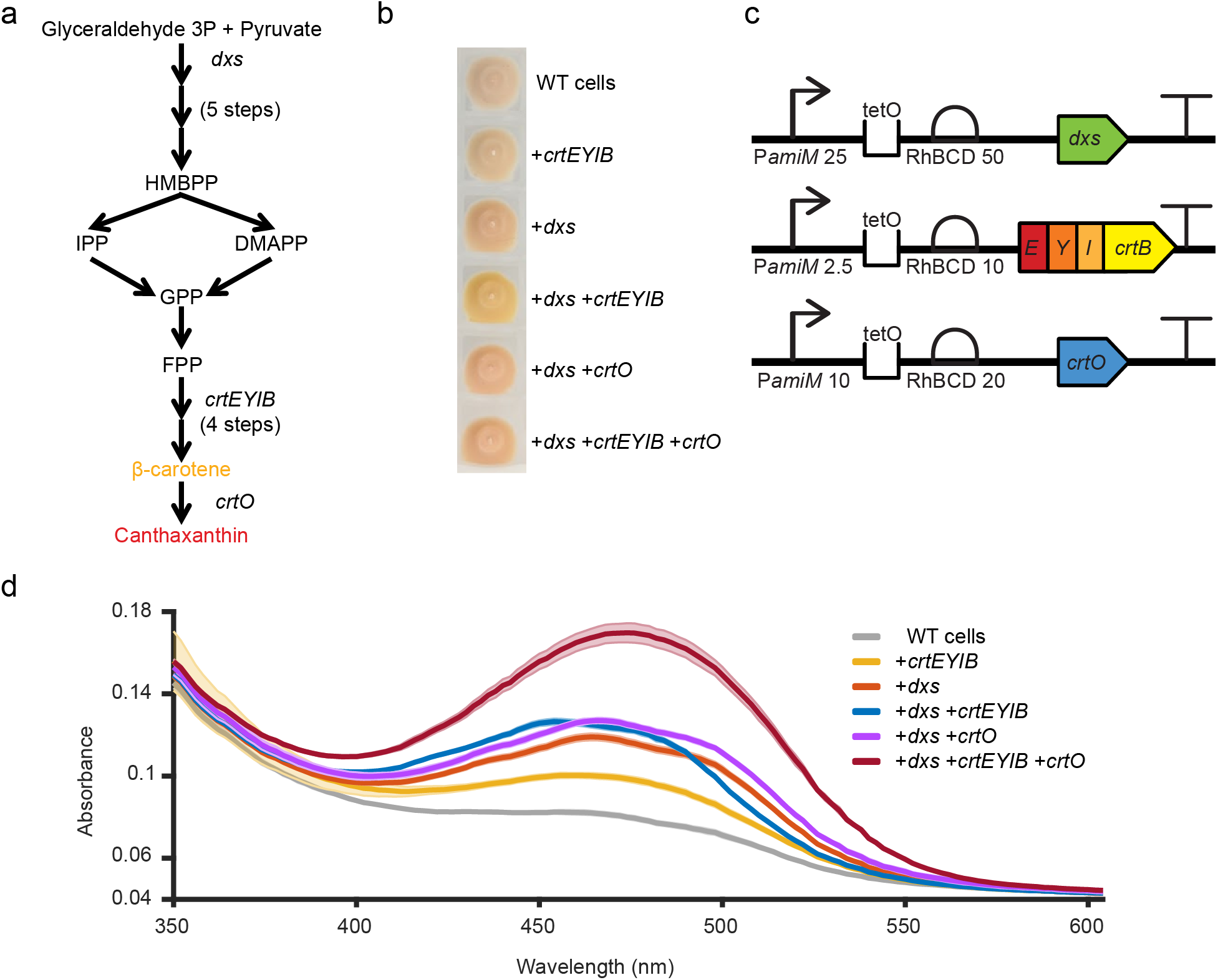
Carotenoid production and quantification in *R. erythropolis*. (a) The carotenoid production pathway for canthaxanthin starting via the MEP pathway. (b) Cell pellets of *R. erythropolis* carrying plasmids expressing different combinations of *dxs, crtEYIB*, and *crtO*. (c) The design of the final transcriptional units used to express each of the genes. (d) The absorbance spectrum of carotenoid pigments extracted from cell pellets of *R. erythropolis* carrying plasmids expressing the different combinations of genes. Shading around the lines represents ± one standard deviation.

We constructed plasmids with different combinations of these genes and transformed them into *R. erythropolis* N9T-4. This strain was selected for these experiments for two reasons. First, under laboratory conditions it accumulates very low levels of colored carotenoids enabling us to easily evaluate the effect of single genes, and second, it is an industrially relevant strain with the ability to replicate under extremely nutrient limiting conditions(55). Initial designs using strong transcriptional and translational control elements resulted in high cellular burdens and genetic instability as determined by colonies with varying color formation and size when transformed with the same construct. Further iterations with progressively weaker transcriptional and translational control elements were constructed resulting in colonies which maintained a consistent color and size, indicating a tolerable metabolic burden (**Figure 6c**). Total carotenoids were extracted using methanol from *R. erythropolis* cell pellets and visualized using UV-Vis spectroscopy (**Figure 6b, 6d**). We observed *dxs* expression alone had a stronger effect on carotenoid pigment production than expressing the *crtEYIB* cassette. The combination of *dxs* and *crtEYIB* expression shifted the absorbance spectrum of the carotenoid pigments towards a peak of 460 nm, the absorbance maxima of β-carotene. When *dxs* is combined with *crtO*, no absorbance shift is observed, suggesting no change in the carotenoid profile. This is consistent with the fact that *Rhodococcus erythropolis* N9T-4 natively encodes a *crtO* gene for synthesis of 4-keto-γ-carotene. When *crtEYIB* is expressed in combination with *dxs* and *crtO*, the absorbance shifts towards a maximum at 475 nm due to expected production of the non-native ketocarotenoid canthaxanthin.

## DISCUSSION

Due to their high concentration of biosynthetic gene clusters for natural products and extreme metabolic flexibility, Mycobacteriales are emerging hosts for bioproduction (1, 10, 14, 56). Tight and tunable control of multiple genes and gene clusters will be essential to fully developing these capabilities and making use of synthetic biology tools in these organisms. In recent years, numerous modular genetic toolkits have been developed that provide fine control of gene expression but have largely been optimized around individual model microorganisms(24–27). These can supplant *ad hoc* DNA assembly methods and collectively represent valuable initiatives that serve to reduce variation between experiments and different research environments. Here, we introduce a modular DNA assembly toolkit that enables the rapid assembly of complex plasmids for use in several different Mycobacteriales. This toolkit contains several new genetic elements that were thoroughly validated in three different genera, *Rhodococcus, Nocardia*, and *Gordonia* and that enable the implementation of synthetic biology methods in a range of environmental isolates and non-model species.

The separation of plasmid backbone assembly and genetic circuit assembly into two hierarchies in this work significantly simplified evaluation of many of the poorly characterized genetic elements. Large DNA fragments such as origins of replication, particularly from high GC-content bacteria, often contain multiple restriction sites for common type IIs restriction enzymes, including BsaI and Esp3I. Removal of individual restriction sites or direct synthesis of new sequences, both of which involve amplification of structured, repetitive, and high GC-content DNA can be time consuming. Additionally, restriction sites that occur within conserved non-coding regions may require complete sequence randomization to isolate new functional sequences. Here, only when functional origins of replication or markers have been identified do researchers need to invest in these processes to bring genetic elements into compliance with the more complex cloning scheme used in the second assembly hierarchy. For future toolkit expansion, additional origins of replication or selectable markers can be evaluated within this framework and interchanged with the parts reported as necessary to enable the rapid development of new shuttle plasmids.

Transcriptional and translational control elements that give predictable expression across different genes and genera are highly enabling for synthetic biology in non-model bacteria. The series of promoters and BCDs described here allow for rapid and reproducible tuning of gene expression across three genera of Mycobacteriales. Furthermore, the architecture of the BCD translational control elements results in predictable translation rates agnostic to the gene in question, which is a major challenge for translational control elements in general. Although we used the highly active P*amiM* promoter from *R. ruber* as the template for our series of promoters, gene expression in *Nocardia* species was much lower than in *Rhodococcus* or *Gordonia*. This result was surprising given the close evolutionary relationship between *Rhodococcus* and *Nocardia*. The reasons for this are unclear and suggest a need to develop *Nocardia* specific promoters in the future.

We identified and fully domesticated five origins of replication that functioned in *R. ruber*, a *Rhodococcus* species with comparatively low genetic tractability, however, we were unable effectively transform seven out of the 17 species tested, including all four *Dietzia*. We attribute this to three possible factors: (1) the use of a generic transformation protocol with no optimization for the different genera, (2) the use of minimal origins which may have resulted in the removal of regions important for stable plasmid maintenance, and (3) the presence of strong restriction-modification barriers in some species. In support of this latter hypothesis, during the course of this work we identified three genes whose presence on a plasmid greatly reduced transformation efficiency; the gene encoding the CymR transcriptional regulator and the *codA* component of the *codA::X::upp* fusion in *R. ruber*, and the *aac(3)-IVa* gene encoding the apramycin resistance marker in *R. rhodochrous*. Using the counter-selectable marker developed in this work, the plasmids were cured from successful transformants and re-transformation of these isolates resulted in a significant increase in transformation efficiency. This observation may be the result of inactivation of a native restriction-modification system, and further characterization of these more genetically tractable strains is underway.

Collectively, these results highlight the need for both methylome analysis and improved annotation of Mycobacteriale origins of replication, which can guide future efforts to determine minimal origins of replication and to eliminate unwanted restriction sites. For this toolkit, most of the origins of replication were previously defined by deletion studies to identify the smallest contiguous fragment capable of autonomous replication. While effective and easily accomplished using traditional molecular biology methods, this approach does not preserve adjacent cryptic features that may promote plasmid maintenance or segregation but are not strictly necessary for independent replication. Future engineering efforts which can make use of new techniques for synthesizing native, high GC-content DNA to characterize these features in plasmids from Mycobacteriales would be broadly useful. Although we focused on evaluating genetic components in three genera, the complete host range of these origins remains to be determined. We anticipate further characterization and development of this system enabling complex yet reproducible molecular biology in several related Mycobacteriales, including *Mycobacterium, Corynebacterium*, and *Tsukamurella*.

## MATERIALS AND METHODS

### Bacterial strains and growth conditions

*Rhodococcus ruber* C208 (DSM 45332), *Rhodococcus erythropolis* N9T-4 (NBRC 110906) and *Rhodococcus imtechensis* (DSM 45091) (syn. *R. opacus)*(57) were cultured at 28 °C in Tryptic Soy Broth (TSB) supplemented with antibiotics at an appropriate concentration for each strain. Apramycin was added at a concentration of 100 μg mL^-1^ for *R. ruber*, 20 μg mL^-1^ for *N. neocaledoniensis*, and 50 μg mL^-1^ for all other Actinomycetota strains. Kanamycin was added at a concentration of 100 μg mL^-1^ for *R. ruber*, and 50 μg mL^-1^for all other Actinomycetota strains. Spectinomycin was used at a concentration of 100 μg mL^-1^ for all Actinomycetota strains. Nourseothricin and zeocin were used at a concentration of 50 μg mL^-1^ for all Actinomycetota strains unless otherwise specified. *E. coli* strain DH10B was used for plasmid construction and was cultured at 37 °C in Terrific Broth supplemented with antibiotics at a concentration of 33 μg mL^-1^ for chloramphenicol or zeocin, and 50 μg mL^-1^ for all other antibiotics used for plasmid maintenance.

#### Plasmid construction and molecular biology

All plasmids were constructed using the hierarchical Golden Gate assembly system reported here with the parts reported for each experiment. Golden Gate assembly reactions were performed with T4 DNA ligase, T4 DNA ligase buffer, and the appropriate restriction enzyme. Reactions were incubated using a thermal cycler at 37 °C for one minute and 16 °C for two minutes, repeated for 25 cycles, followed by incubation at 37 °C for 30 minutes and enzyme inactivation at 85 °C for 15 minutes. Assemblies were transformed into *E. coli* and a plasmid miniprep was recovered to provide supercoiled DNA for electroporation into Mycobacteriales. For plasmid recovery from *Rhodococcus* cells, pellets of *R. ruber* cells were resuspended in 2 mL of 15 mg mL^-1^ lysozyme and incubated at 37 °C for one hour at 200 rpm. Plasmids were then isolated using a standard alkaline lysis plasmid miniprep protocol.

### Promoter and BCD library construction

Construction of the libraries of promoters was performed by PCR using oligonucleotide primers containing degenerative bases. PCR amplicons were circularized using Gibson Assembly for 16 hr. Assembled libraries were desalted and transformed into *E. coli* to generate supercoiled plasmid DNA which was transformed into *R. ruber* as previously described.

### Multispecies Mycobacteriale electroporation protocol

Electrocompetent *Rhodococcus, Nocardia*, and *Gordonia* cells were prepared by culturing the bacteria in 200 mL TSB containing 1.5% w/v glycine and 1.5% w/v sucrose. Cells were cultured at 28 ºC for 16 h and then washed three times with 20 mL sterile 10% glycerol, split into 200 μL aliquots and flash frozen in liquid nitrogen. Cell aliquots were stored at -80 ºC until use.

100 μL aliquots of electrocompetent *Rhodococcus, Nocardia*, or *Gordonia* cells were mixed with 100 ng of plasmid DNA, transferred to 0.2 mm cuvettes, and electroporated at 2.5 kV with a capacitance of 25 μF. Cells were recovered in 2 mL TSB at 28 °C for three hours with 250 rpm agitation, harvested by centrifugation and resuspended in 150 μL TSB, and transferred to petri plates containing solid tryptic soy agar medium supplemented with an antibiotic. Plates were incubated at 28 °C for several days until colonies were observed.

### Fluorescence assays

For *Rhodococcus, Nocardia* and *Gordonia* species, transformants were selected and cultured in 24-well deep-well plates agitated at 500 rpm in TSB supplemented with apramycin at 28 °C. Cultures were incubated for two to three days with constant agitation until reaching stationary phase. Subcultures were started using 50 μL of confluent culture to inoculate 2 mL of fresh medium in 24-well deep-well plates. These cultures were incubated at 28 °C for two days. 500 μL aliquots from each well were centrifuged, and the cell pellets were resuspended in an equal volume of PBS pH 7.4. 100 μL of cell suspension was transferred to a 96-well microtiter plate and assayed by measuring absorbance at 600 nm and fluorescence using an λex 595 nm and λem 650 nm.

### Plasmid curing

*R. ruber* and *G. paraffinivorans* were electroporated with a plasmid encoding an E2-Crimson expression cassette and one of the three modular counter-selectable markers. Single E2-Crimson positive transformants were selected and cultured in TSB to confluence. 10 μL aliquots of bacterial culture were then passaged into one of three different media conditions: TSB with no additives, TSB with the 50 μg mL^-1^ of 5-FC, or TSB with the antibiotic matching the counter-selectable marker at concentrations of 100 μg mL^-1^ apramycin, 100 μg mL^-1^ spectinomycin, and 33 μg mL^-1^ zeocin for *R. ruber*, and 50 μg mL^-1^ apramycin, 100 μg mL^-1^ spectinomycin and 33 μg mL^-1^ zeocin for *G. paraffinivorans*. Cultures were grown to confluence after which 5 μL of medium was struck out onto TSB agar plates for imaging or the cell population was evaluated using cytometry.

### Isoprenoid measurements

*Rhodococcus erythropolis* transformants were cultured in 24-well deep-well plates in TSB supplemented with apramycin at 28 °C. Cultures were incubated for two to three days with constant agitation until reaching stationary phase. 1 mL of culture was transferred to a new deep well plate and centrifuged, after which the cell pellet was resuspended in 500 μL of 100% methanol. The pellet was allowed to settle for three hours in the fridge, then 100 μL of methanol extraction was transferred to a 96-well microtiter plate and assayed by measuring the absorbance spectrum from 350 nm to 600 nm.

### Statistical analysis and data visualization

Unless otherwise indicated, all data was collected from a minimum of three biological replicates and two technical replicates. Error bars represent the standard deviation of the biological replicates.

## Data availability

Plasmids which comprise the RhoClo toolkit are available as a complete set from the Genome Design and Engineering Center (GDEC) at Rice University. All DNA sequences for relevant plasmids and modular parts are provided in the Appendix.

## AUTHOR CONTRIBUTIONS

Z.J., N.D.S., S.H., and R.T. conceived the study, designed the experiments, and coordinated the experimental work. Z.J., A.A., and Q.W. performed the experiments. Z.J., A.A., Q.W., A.R.G., D.L.K and R.T. analyzed the data and contributed to the experimental troubleshooting. All authors contributed to writing and editing the manuscript.

## ACKNOWLEDGEMENTS

Note: The authors declare no competing financial interests.

This work was supported by funding from the National Institute of Standards and Technology to R.T. (70NANB21H102). This research was conducted utilizing the DEVCOM Army Research Laboratory Cooperative Research and Development Agreement CRADA-18-046-J004.

## Notes

### Competing Interest Statement

The authors have declared no competing interest.

