## Supporting Information for "A modular toolkit for environmental *Rhodococcus, Gordonia*, and *Nocardia* enables complex metabolic manipulation"

This file contains:
Supplementary Results
Figures A1-A5
Tables A1-A9

### SUPPLEMENTARY RESULTS

#### Optimization of a tetracycline-inducible genetic circuit containing a single *tetO* operator

One of the advantages of using a modular cloning framework is the ability to rapidly prototype new genetic circuits by tuning expression of key genes using a combination of transcriptional and translational control elements. This provides an unprecedented level of control and flexibility which is useful when adapting components to new genetic contexts, particularly legacy materials which have either never been modularized or evaluated in non-model species. The tetracycline-inducible system described in this work was adapted from well-characterized genetic elements (TetR and *tetO*) which are in common use in *E. coli* and have previously been shown to function in *Rhodococcus* and other Mycobacteriales(1, 2). In the most common configuration, this system uses two *tetO* operators, one located between the -35 and -10 regions of the promoter, and the second downstream of the -10 region. This arrangement achieves highly effective repression and can be constructed as a part encompassing both the promoter and operator functionalities. We sought to build a fully modular series of constitutive promoters that can be converted into inducible promoters with the addition of a single operator site directly adjacent to the -10 site of the promoters. This required an iterative series of experiments to balance the expression of the TetR transcriptional repressor and reporter gene which we have described in detail as it may prove helpful for other researchers.

This genetic circuit consists of two transcriptional units, the first expressing the far-red fluorescent protein E2-Crimson under the control of a synthetic promoter, the tet operator, and a BCD, and the second expressing the TetR transcriptional repressor (**Figure A5**). Our initial configuration used a moderately strong promoter (P*amiM* 25) and a very strong BCD (RhBCD 100) to drive expression of E2-Crimson, and a weak promoter and BCD upstream of TetR. When this circuit was evaluated for E2-Crimson expression using a gradient of aTc concentrations, the results showed that while red fluorescence increased with increasing aTc concentration, neither full repression nor de-repression were achieved (**Figure A5a**). These are both important parameters and largely determine the induction of the reporter in response to given concentration of aTc and the dynamic range for the inducer molecule. This initial iteration of the circuit achieved ~5-fold induction at the highest tested aTc concentration.

During our first attempt to improve circuit performance, we made two key changes. First, we assembled a new E2-Crimson transcriptional unit driven by the strong promoter (P*amiM* T2A) but paired it with a much weaker BCD (RhBCD 20). This change was designed to make E2-Crimson expression more responsive to transcriptional regulation. Secondly, we significantly increased the strength of the promoter and BCD driving expression of TetR to ensure tighter repression. While this second iteration of the circuit successfully lowered the background signal when no inducer was present (to near zero), excessive levels of TetR prevented induction in the presence of aTc, even at elevated concentrations (**Figure A5b**). To resolve this issue, we retained the new E2-Crimson transcriptional unit and adjusted the promoter and BCD to reduce expression of TetR. Evaluation of this circuit showed that it retained near zero background signal in the absence of aTc while achieving 55-fold induction of red-fluorescence at the maximal inducer concentration (**Figure A5c**) A fourth circuit design which further lowered TetR expression led to a 3-fold increase in the background signal and was not pursued further (not shown).

a

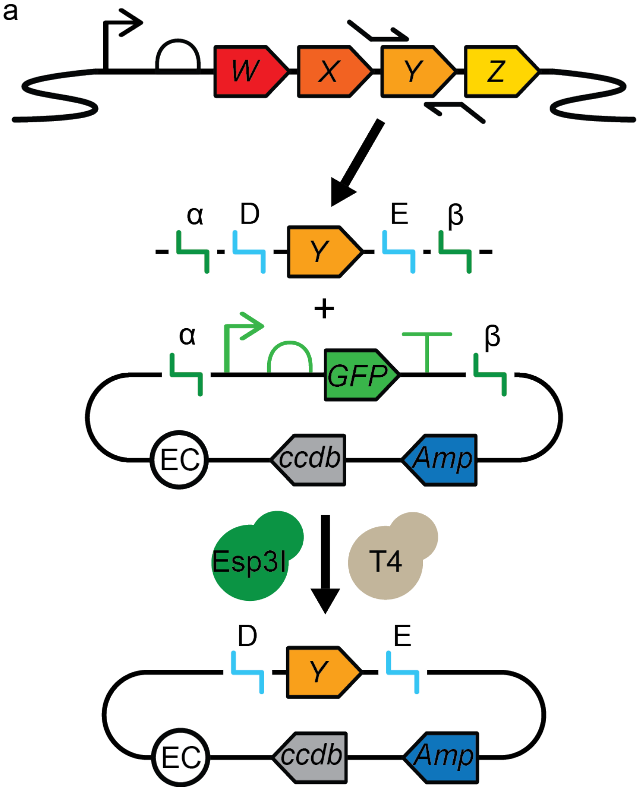


b

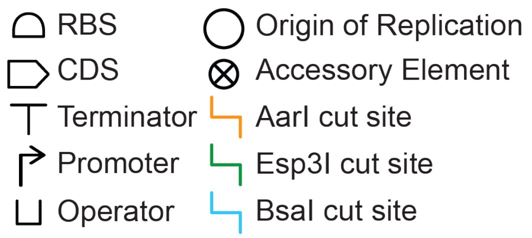


c

| Esp3I Overhangs<br>CGTCTCNNNNNN<br>GCAGAGNNNNNN | BsaI Overhangs<br>GGTCTCNNNNNN<br>CCAGAGNNNNNN |
|---|---|
| α: GATA | A: AACG |
| β: ATCG | B: CAAT |
| 1: CTGA | C: GGAG |
| 2: CCAA | D: TATG |
| 3: GATG | E: ATCC |
| 4: AGCA | F: GCTG |

**Figure A1:** Workflow for assembly of a single Part plasmid. Template DNA for individual parts may include legacy plasmids, genomic DNA (depicted), or synthetic DNA. Here, oligonucleotide primers are designed to flank the relevant genetic element and incorporate the necessary type IIs restriction sites and corresponding 4 bp overhangs. The resulting amplicon can be combined with a part dropout plasmid in an Esp3I-mediated Golden Gate assembly to generate a part plasmid where the relevant genetic element is flanked solely by BsaI sites which enable further assembly into transcriptional units. Part plasmids encode the CcdB toxin and must be maintained in a CcdB-resistant *E. coli* strain such as DB3.1. This toxin prevents Part plasmids from contaminating further Golden Gate assemblies which are performed in CcdB-sensitive *E. coli* strains. (b) A key to the SBOL symbol for each genetic element. (c) The restriction sites and overhangs used in the toolkit.

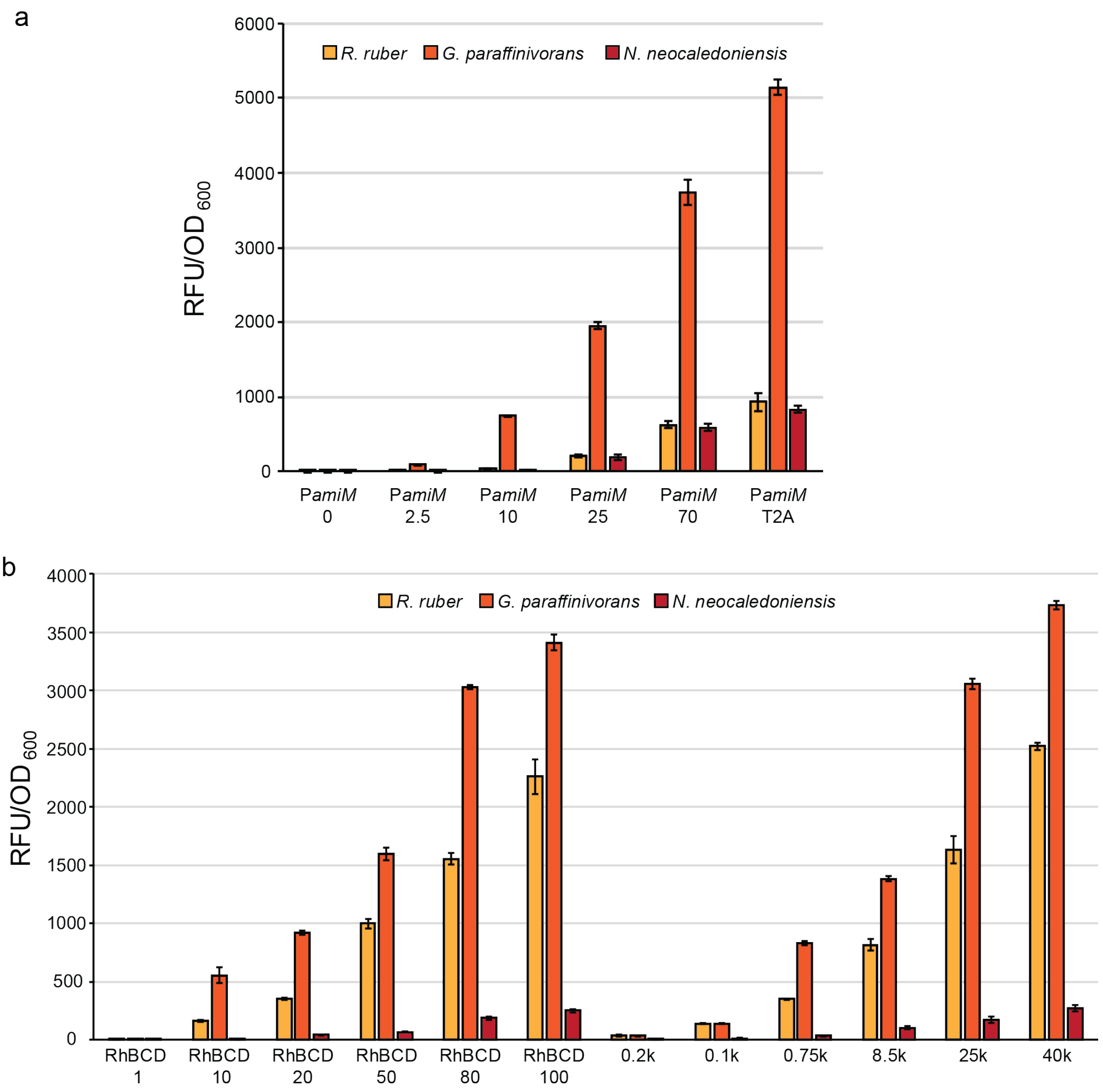


**Figure A2:** Non-normalized fluorescence assay data for the transcriptional and translational control elements collected in the three Mycobacteriale species. (a) The fluorescence values of the series of synthetic promoter variants normalized only to cell density in *R. ruber, G. paraffinivorans*, and *N. neocaledoniensis*. (b) The fluorescence of the two series of BCDs tested in *R. ruber, G. paraffinivorans*, *and N. neocaledoniensis* normalized only to cell density.

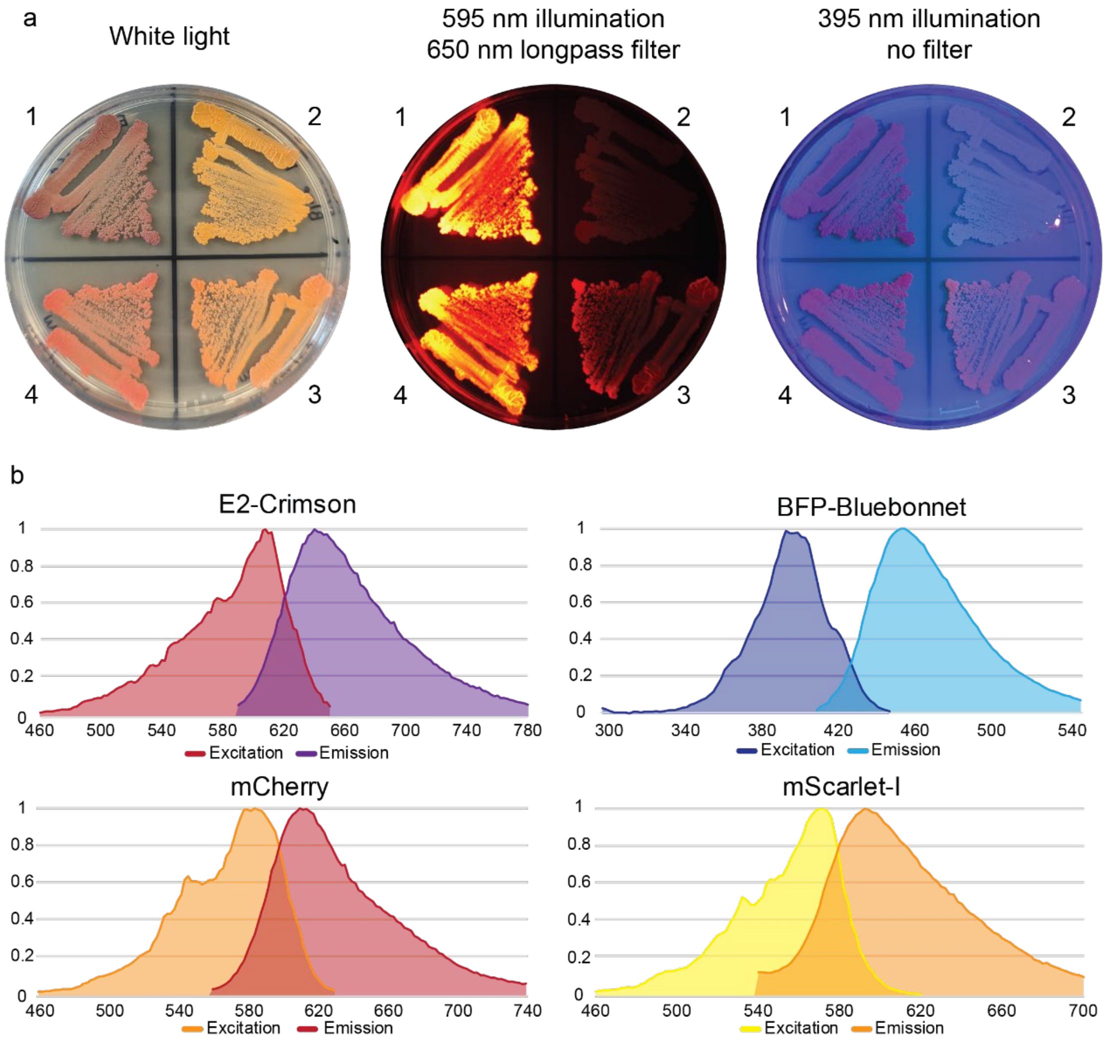


**Figure A3:** Red and blue fluorescent proteins are functional in *Rhodococcus ruber* C208. (a) *Rhodococcus ruber* C208 transformed with plasmids expressing four different fluorescent proteins. The first image is illuminated with white light, the second image is illuminated with a 595 nm amber LED light source and imaged through a 650 nm longpass filter. The third image is illuminated using a UV light at 395 nm under ambient lighting and imaged without any filter. The first quadrant contains cells expressing E2-Crimson, the second quadrant BFP-Bluebonnet, the third quadrant mScarlet-I, and the fourth quadrant mCherry. (b) Excitation and emission spectra for each of the four fluorescent proteins collected in *R. ruber* cells.

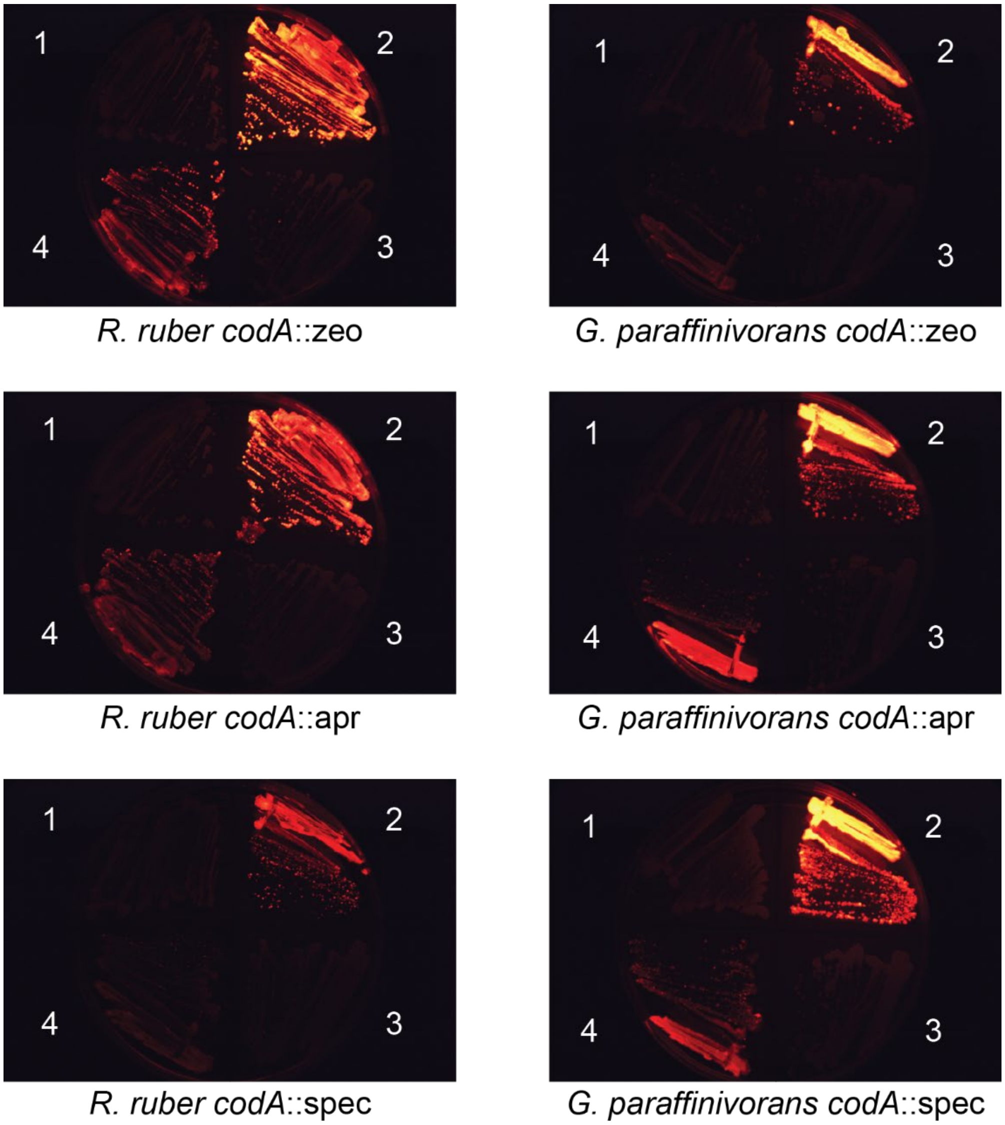


**Figure A4:** The *CodA* fusion counter-selectable markers tested across *R. ruber* and *G. paraffinivorans.* Quadrant one shows the indicated strain without a plasmid. Quadrant two shows the indicated strain containing the counter-selection plasmid with the corresponding antibiotic. Quadrant three shows the strain after undergoing counter-selection with 50 mg $ml^{-1}$ 5-FC. Quadrant four shows the same isolate passaged in TSB media without any antibiotic or 5-FC. Quadrants 2-4 represent a single clonal transformant.

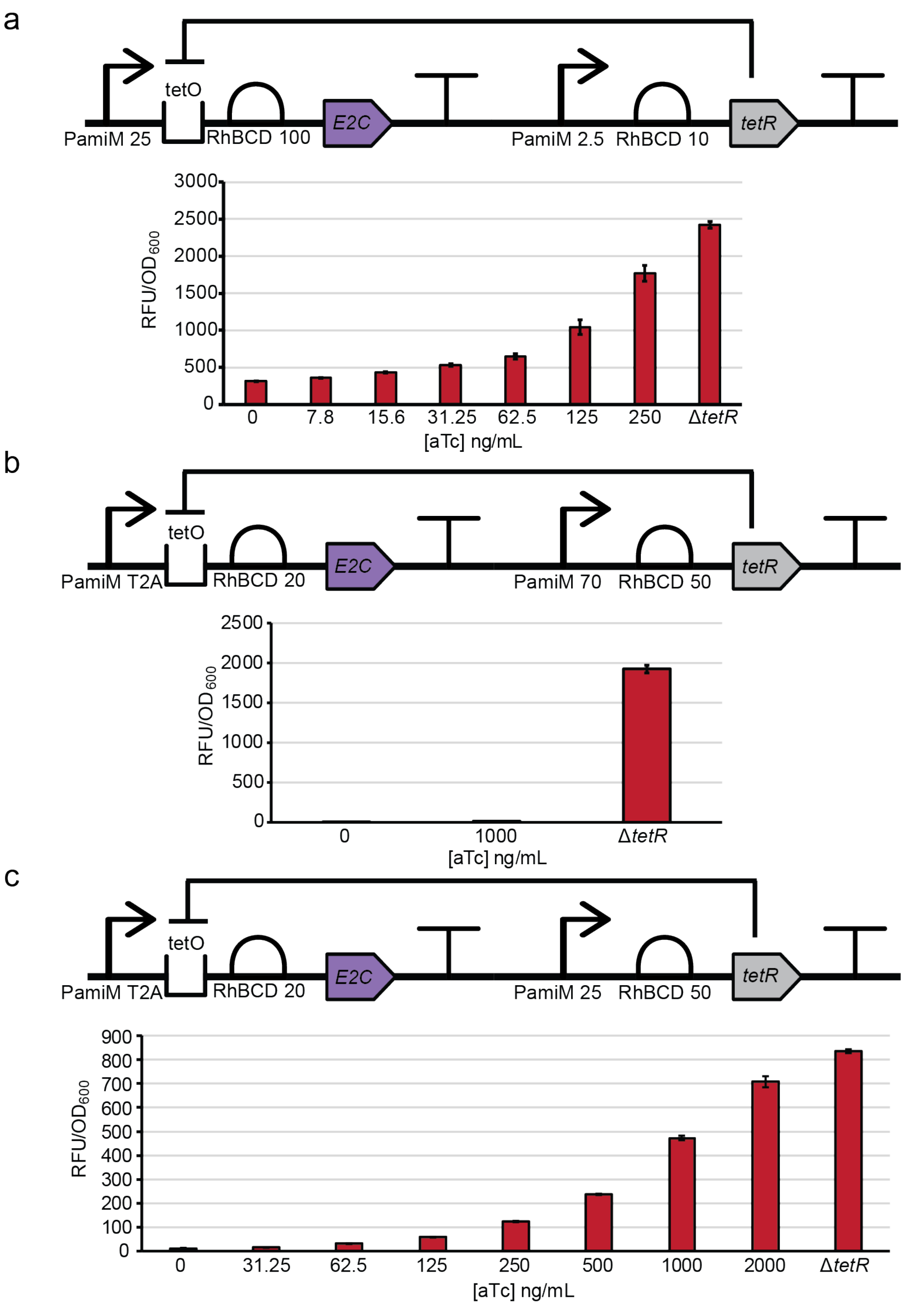


**Figure A5:** Optimization of a tetracycline-inducible genetic circuit. (a) Circuit schematic and fluorescence assay for the initial circuit design using a gradient of aTc concentrations. (b) Circuit schematic and fluorescence assay for a modified circuit which was designed to lower background signal in the absence of aTc. No de-repression was observed. (c) Circuit schematic and fluorescence assay for the final circuit design using a gradient of aTc concentrations. This circuit retains low background signal and achieves nearly maximal induction at high aTc concentrations.

**Table A1:** List of all Actinomycete origins of replication used in this work.

| Origin | Species | Source | Replication | *R. ruber* transformants |
|---|---|---|---|---|
| pAL5000S | *Mycobacterium fortuitum* (3) | Addgene: 119886-pDD131 | θ | - |
| pB264 | *Rhodococcus* sp. B264 (4) | Addgene: 119879-pDD120 | θ | - |
| pN30 | *Rhodococcus erythropolis* 30 (5) | Synthetic | θ | - |
| pNC500 | *Rhodococcus rhodochrous* B-267 (6) | Synthetic | θ | Yes |
| pNC903 | *Rhodococcus ruber* P-II-123-1 (7) | Synthetic | θ | Yes |
| pOTS | *Rhodococcus equi* 103 (8, 9) | Synthetic | θ | n/a - toxic in *E. coli* |
| pPRH | *Arthrobacter rhombi* PRH1 (10) | Synthetic | θ | - |
| pGA1 | *Corynebacterium glutamicum* LP-6 (11) | Synthetic | RCR | - |
| pNG2 | *Corynebacterium diptheriae* (12, 13) | Synthetic | RCR | Yes |
| pRET1100 | *Rhodococcus erythropolis* IAM1400 (14) | Synthetic | RCR | Yes |
| pSR1 | *Corynebacterium glutamicum* ATCC 19223 (15) | Addgene: 59449-pSRKBB-empty | RCR | - |
| pXT107 | *Nocardia* sp. 107 (16) | Synthetic | RCR | Yes - unstable |
| pYS1 | *Nocardia aobensis* IFM 10795 (17) | Synthetic | RCR | Yes – unable to domesticate |
| pJAZ38 | *Mycobacterium fortuitum* 138 (18) | Addgene: 32361-pMycVec2 | LR7-family | - |
| pKB1 | *Gordonia westfalica* Kb1 (19) | Synthetic | LR7-family | - |
| pMF1 | *Mycobacterium fortuitum* 110 (20) | Synthetic | LR7-family | - |
| pMyong2 | *Mycobacterium yongonense* DSM $45126^{T}$ (21) | Synthetic | LR7-family | - |
| pSOX | *Rhodococcus* sp. X309 (22) | Synthetic | LR7-family | Yes |
| pVT2 | *Mycobacterium avium* MD1 (23) | Synthetic | LR7-family | n/a - toxic in *E. coli* |
| pDA71 | *Rhodococcus erythropolis* (24) | ATCC: 77474 | Nocardiophage Q4 | - |

**Table A2:** DNA sequences of all Actinomycete origins of replication used in this work. AarI restriction enzyme recognition sequences are highlighted in green. The cut site is underlined and in bold.

| Origin |
| --- |
| pAL5000S |
| CACTAGCTCAGATTCAGTAGACCGCTGTTGCACCTGCATTA**TACA**CTCAGCGAAATATCTGACTTGGAG<br>CTCGTGTCGGACCATACACCGGTGATTAATCGTGGTCTACTACCAAGCGTGAGCCACGTCGCCGACGAA<br>TTTGAGCAGCTCTGGCTGCCGTACTGGCCGCTGGCAAGCGACGATCTGCTCGAGGGGATCTACCGCCAA<br>AGCCGCGCGTCGGCCCTAGGCCGCCGGTACATCGAGGCGAACCCAACAGCGCTGGCAAACCTGCTGGTC<br>GTGGACGTAGACCATCCAGACGCAGCGCTCCGAGCGCTCAGCGCCCGGGGGTCCCATCCGCTGCCCAAC<br>GCGATCGTGGGCAATCGCGCCAACGGCCACGCACACGCAGTGTGGGCACTCAACGCCCCTGTTCCACGC<br>ACCGAATACGCGCGGCGTAAGCCGCTCGCATACATGGCGGCGTGCGCCGAAGGCCTTCGGCGCGCCGTG<br>GACGGCGACCGCAGTTACTCAGGCCTCATGACCAAAAACCCCGGCCACATCGCCTGGGAAACGGAATGG<br>CTCCACTCAGATCTCTACACACTCAGCCACATCGAGGCCGAGCTCGGCGCGAACATGCCACCGCCGCGC<br>TGGCGTCAGCAGACCACGTACAAAGCGGCTCCGACGCCGCTAGGGCGGAATTGCGCACTGTTCGATTCC<br>GTCAGGTTGTGGGCCTATCGTCCCGCCCTCATGCGGATCTACCTGCCGACCCGGAACGTGGACGGACTC<br>GGCCGCGCGATCTATGCCGAGTGCCACGCGCGAAACGCCGAGTTCCCGTGCAACGACGTGTGTCCCGGA<br>CCGCTACCGGACAGCGAGGTCCGCGCCATCGCCAACAGCATTTGGCGTTGGATCACAACCAAGTCGCGC<br>ATTTGGGCGGACGGGATCGTGGTCTACGAGGCCACACTCAGTGCGCGCCAGTCGGCCATCTCGCGGAAG<br>GGCGCAGCAGCGCGCACGGCGGCGAGCACAGTTGCGCGGCGCGCAAAGTCCGCGTCAGCCATGGAGGCA<br>TTGCTATGAGCGACGGCTACAGCGACGGCTACAACCGGCAGCCGACTGTCCGCAAAAAGCGGCGCGTGA<br>CCGCCGCCGAAGGCGCTCGAATCACCGGACTATCCGAACGCCACGTCGTCCGGCTCGTGGCGCAGGAAC<br>GCAGCGAGTGGCTCGCCGAGCAGGCTGCACGCCGCGAACGCATCCGCGCCTATCACGACGACGAGGGCC<br>ACTCTTGGCCGCAAACGGCCAAACATTTCGGGCTGCATCTGGACACCGTTAAGCGACTCGGCTATCGGG<br>CGAGGAAAGAGCGTGCGGCAGAACAGGAAGCGGCTCAAAAGGCCCACAACGAAGCCGACAATCCACCGC<br>TGTTCTAACGCAATTGGGGAGCGGGTGTCGCGGGGGTTCCGTGGGGGGTTCCGTTGCAACGGGTCGGAC<br>AGGTAAAAGTCCTGGTAGACGCTAGCTTCACGCTGCCGCAAGCACTCAGGGCGCAAGGGCTGCTAAAGG<br>AAGCGGAACACGTAGAAAGCCAGTCCGCAGAAACGGTGCTGACCCCGGATGAATGTCAGACGCAAGCGC<br>AAAGAGAAAGCAGGTAGCTTGCAGTGGGCTTACATGGCGATAGCTAGACTG**ACAG**ATTAGCAGGTGTGC<br>ATGATCTACGTGCGTCACATGCAGTAC |
| pB264 |
| CACTAGCTCAGATTCAGTAGACCGCTGTTGCACCTGCATTA**TACA**ACCGAAACATCTGACTTGGCTTCA<br>CGATTAACTTTGAACACACCGAGGGAAATGCCCAGGTTTCGTTCGAGAGCATGCGACGGTCACCAGCAT<br>GGACACACACGGGGTGCGGGAAGGTGGGGACTGGGAGCAAATGTGGCTACCGTTCTGGCCTTTGGCAAC<br>AAACGACTTCCTTGAGGGGGTCTATCGAATGCGCCGCCCTGCCGCACTTGAGCGGCGCTACATCGAGGC<br>CAACCCGCAGACACTGAGCAATCTGCTCGTCGTCGACGTCGACCACCCCGACTCCGCCCTGCGCGCACT<br>GTCAGCGGCAGGCAATCACCCTATGCCGAACGCCGTGATCGAAAACAGCAGCAACGGGCATGCGCACCT<br>GCACTGGTGGCTGCGCGAGCCGTTCACCCGCACCGAGTACGCCCGACGGAAGCCCCTCGCGTACGCCGC<br>CGCCGTTACCGAAGGGCTACGCCGGGCCGTCGAAGGGGACATCGGCTACTCAGGCCTGATGACCAAGAA<br>CCCCACACACTCAGGCTGGGACACGCACTGGATCCACACCGAACCGCGCAGCCTCGCCGAGCTCGAGGC<br>CGAACTCGGCACACACATGCCCTCCCCGCGCTGGCAGCACACCAAGGCCCACCGTGACGCCCCCATAGG<br>ACTCGGCCGCAACTGCGCAATCTTCCACGCCGCTCGCACCTGGGCCACCCGCCCCGGCCTCATGCGCAA<br>CTACCTGCCCACTCATGACAGCGCCGGCCTCGAACTCGCCCTCCACCGCGAAGTCACCGCCCTCAACGC<br>CAGCTACACCGAGGAACTCCCACCCTCTGAAGCCCGCGCCATCGCCGCCAGCATTCACCGATGGATCAC<br>CACGCGATCCCGCATCTGGAAAGACGGTATCGCCGTCTACGAAGCCACACTCTCGACCATCCAATCCGC<br>CCGTGGCCGCAAAGGCGGCGTCGCCAGCGGCCAGGCCCGGCGTGCACGCAGCACCAAACTGGATCAGGT<br>GCGACTGGCAATTGAGGAGGCCACACATGACTGAGCGTCTGCCTCGCAACGGGATAACAGTGCGGGAGC<br>TCGCCGAACGCACAGGCGTATCGACGGCCAGCATCATCCGGTGGACATCAGAGCCGCGAGAGGCCTACC<br>TGTCCCGGGCACAGCAACGGCGCGAGAAGATACGGGAGCTACGCGGTACTGGCATGTCCATGAGAGCTA<br>TCGCCAATGAGCTGGGATGCTCGGTCGGCACTGTCCATCACGCCCTGGCAGGAATCGAGCGATGAGTAT |

CAGCTGCGATGACGTCCACTACCAGCCCGTCCCCCTTGGGACAGCCCTGCCGTGCGGGTGCGTTGCAC**A**
**CAG**ATTAGCAGGTGTGCATGATCTACGTGCGTCACATGCAGTAC

pN30

CACTAGCTCAGATTCAGTAGACCGCTGTTGCACCTGCATTA**TACA**CAGCTCCCGGGAGTACCGCCGTAC
TCACCCGCCTGTGCTCACCATCCACCGACGCAAAGCCCAACCCGAGCACACCTCTTGCACCAAGGTGCC
GACCGTGGCTTTCCGCTCGCAGGGTTCCAGAAGAAATCGAACGATCCAGCGCGGCAAGGTTCAAAAAGC
AGGGGTTGGTGGGGAGGAGGTTTTGGGGGGTGTCGCCGGGATACCTGATATGGCTTTGTTTTGCGTAGT
CGAATAATTTTCCATATAGCCTCGGCGCGTCGGACTCGAATAGTTGATGTGGGCGGGCACAGTTGCCCC
ATGAAATCCGCAACGGGGGGCGTGCTGAGCGATCGGCAATGGGCGGATGCGGTGTTGCTTCCGCACCGG
CCGTTCGCGACGAACAACCTCCAACGAGGTCAGTACCGGATGAGCCGCGACGACGCATTGGCAATGCGG
TACGTCGAGCATTCACCGCACGCGTTGCTCGGATCCATCGTCATCGACTGCGATCACGTCGATGCCGCG
ATGCGCGCATTCGAGCAACCATCCGACCATCCGGCGCCGAACTGGGTCGCACAATCGCCGTCCGGTCGC
GCACATATCGGATGGTGGCTCGGCCCCAACCACGTGTGCCGCACCGATAGCGCACGACTGACGCCACTG
CGCTACGCACACCGCATCGAAACCGGCCTCAAGATCAGCGTCGGCGGCGATTTCGCGTATGGCGGGCAA
CTGACCAAAAACCCGATTCACCCCGATTGGGAAACGATCTACGGTCCGGCAACCCCGTATACATTGCGG
CAGCTGGCAACCATCCACACACCCCGGCAGATGCCGCGTCGGCCCGATCGGGCAGTGGGTCTGGGTCGC
AACGTCACCATGTTCGACGCCACCCGGCGATGGGCATACCCGCAGTGGTGGCAACACCGAAACGGAACC
GGCCGCGACTGGGACCATCTCGTCCTGCAGCACTGCCACGCCGTCAACACCGAGTTCACGACACCACTG
CCGTTCACCGAAGTACGCGCCACCGCGCAATCCATCTCCAAATGGATCTGGCGCAATTTCACCGAAGAA
CAGTACCGAGCCCGACAAGCGCATCTCGGTCAAAAAGGCGGCAAGGCAACGACACTCGCCAAACAAGAA
GCCGTCCGAAACAATGCAAGAAAGTACGACGAACATACGATGCGAGAGGCGATTATCTGATGGGCGGAG
CCAAAAATCCGGTGCGCCGAAAGATGACGGCAGCAGCAGCAGCCGAAAAATTCGGTGCCTCCACTCGCA
CAATCCAACGCTTGTTTGCTGAGCCGCGTGACGATTACCTCGGCCGTGCGAAAGCTCGCCGTGACAAAG
CTGTCGAGCTGCGGAAGCAGGGGTTGAAGTACCGGGAAATCGCCGAAGCGATGGAACTCTCGACCGGGA
TCGTCGGCCGATTACTGCACGACGCCCGCAGGCACGGCGAGATTTCAGCGGAAGATCTGTCGGCGTAAC
CAAGTCAGCGGGTTGTCGGGTTCCGGCCGGCGCTCGGCACTCGGACCGGCCGGCGGATGGTGTTCTGCC
TCTGGCGCAGCGTCAGCTACCGCCGAAGGCCTGTCATCGACCGGCTTCGACTGAAGTATGAGCAACGTC
ACAGCCTGTGATTGGATGATCCGCTCACGCTCGACCGCTACCTGTTCAGCTGCCGCCCGCTGGGCATGA
GCAACGGCCAACTCTCGTTCAAGATCCGCGATCCGCTGGACATGGTCAAAAT**ACAG**ATTAGCAGGTGTG
CATGATCTACGTGCGTCACATGCAGTAC

pNC500

CACTAGCTCAGATTCAGTAGACCGCTGTTGCACCTGCATTA**TACA**CTTAGCTAATCAACTAGTGGTTGG
TACAGGTGGTTGGGACAACGATCCCCTCGACTTGGTTCGTAACTCCGGCCCCTGGCGACCCTGTTGACG
CCGACCTCGCCGAGTATGAACGCGGCTGGAACGACGGCTGGACCGAGTACTTCGCGGCCCGCATCTGAC
CCCTGGGGCTTACTCAGAGCCGTACCCGGCTTACCTTCCGCTGTCGACTACCCAAACTGCACCACCCGA
ACGGGGGGACTGGTGGGAGTCCCGTTCACCGTAGTAACCCTCAAGGAGTGAGTGAGGGTTGTGTGGGGA
TTCCGGGATACCTGATAAGGCTTTTGGTTCGTCGTACCCCAATATTTTTTCGAGTCTGCGGCGTGTCGC
ACACCAATATTTGGTTCCTTCTGCGGAATCGTGTCCGGCATGACTTCGCCGGGATCGGTCCTCACGCCT
GAGCAATGGGGGCGTGATGTGCTGCTGCCGCACCGTCCGCACGCGACGAACCACTTGGACAGGGGTCAG
TACCGCTACAGCCGCGACGACGCCCTGCTGATGCGCTACATCGAGCACTCGCCGCACGCGCTGCTCGGG
TCAATCGTCGTCGATTGTGACCACCCTGACGCCGCCCTGCGTGCGTTCGAGCAGCCCTCGGACCATCCG
ATGCCGAACTGGGTGGCCCAGTCGCCCTCCGGTCGGGCACACCTGGGCTGGTGGCTCGCCGCCCCGGTC
TGCCGAACCGACTCCGCTCGGTTGCAGCCGCTACGGTTCGCGCACCGCGTCGAGCGGGGCCTCTGCATC
TCTGTGAGCGGCGATTTCGCCTACGGCGGCCAACTCACCAAGAACCCGATCCATCCCGACTGGGAAACG
ATCTACGGACCGTCCACGCCATACGAGCTGCGCGACCTGGTCACCATCCACACGCCCCGGCAGATGCCG
CGCCGTCCTGACCGGGCCGCCGGCCTCGGCCGCAACGTCACCATGTTCGACACTGCCCGCCTCTGGGCC
TATGGGCAGTGGTGGGAACACCGGCACGGCGGCGTCGACGCCTGGCTCCAGCTCGTCCTGCAGCGCTGC
CATGCCATCAATACCGAATTCGCCGATCCGCTCCCCTTCGTGGAGGTCCGCGCCACGGCGCATAGCGTG
GGCAAGTGGATCTGGCGCAATTTCGATGAAGTGGCGTATCGCGCCCGCCAGGCCGAACTCGGCCGGAAG
GGCGGCAAGATCGGCGGCAAGATCGGCGGCCGCGTTGTCTCCCCTGCCAAAATCGAGGCTAATCGTCAG
CGCCGCATCAAGATTGATCGCGCATTAGCTTTAGAGGTGTTACGATGACCGAGCACATCCCCGTCCGCA

AGCGGAAGGGCGCGGAAACCCGTGCGCGCCGGACCATGACCGCCAAGGAGGGCGCGGCCCGACTCGGTG
TCTCGCCCCGTACGATTCAGCGGATCATCGCCGAACCTAGAGAGGAATTCGAGGCCAGGGCACGAGAAC
GGCGAGAACGCGCCGCTGAGCTGCGGGCCTCGGGCCTGAAATACAAGGAGATCGCAGAGGAAATGGGCA
TCTCTACCGGTGCCGTCGGCGCGCTGCTGCACGAAGCACGCAAGTACTCGGTCAGCTCCTAACCGAGAA
GCCGTCGTCGTATCCGGGCCAGGGCACCCGTAACCGCCCGACTATCGGCTGCCGGGGCAGGAATGGGCG
GCGGATCTTCCGCAGCGTCTGTGGGCTCGGGGGCCGACACTGGCCGTGCCTCGAGCATCCGGAGCATCT
GCTGCTGCACATCGATGATCCGATTGCGTTCGCTGGCCAAGGCTTCGGCCACGGCCAGGCGCTGGCGGA
GCTCGGCGTTCTCGGCGGCCAGATCACGGACCCGGTCAGCTGACGTGGGCACCGGGTCAGGAGCGGCCA
TTCTTCCTGGTCTAAGCCCTGCCGCCGATAGGTCACCGAGGGGGATCATCCACCTTCCGTCGGCGGTCT
GCGCCGCATTTGGGAACGCCCCGGCCCGCAGTCGGCGTTTGATGGTGCTCTCGCTGGTCTGCGTCGCGT
CCACGGCCTCTTTGATGGTCAACTTGGGTAGGTCATGGTTCCCGGTCACCGCGTCATCGTGACCCGATA
ATCCAGTTCGCTCGCGCATTTCACCAGCGTGTTGGCAGGTTCACGCCACCGGGTCAACTTCACCGTGAA
CCGGGGCACACCGGTTCCGGGTCACGGCAGTAGTCGCCGAAGAATCCAACGAAAGGTTCCGGCCTTCAT
CCCCCTGCTCCGTGCCAGTTTCGACCGGGAGTCCAAGCTCGGGGAATCCCTATTGATAAAGTCGACGTT
AACTAGTAAATCGCCAGGTTAGGAGCCTTGATGAGCCCAGGAAAGCGTCGTCACCCACGCGGTGCGCTG
AAGGAGTCGATCCGCGAGTACTTGGCTGGTCTGAACGGTGATCCGGCGAGCATTGCCGAGATCACTGCG
GCGATAGAGCCGAAAATCGGGACTCCCGTGCCTTCGTCGGTCAGGACCACGCTGCAGGATGATCGCTAC
TTCGTTCGTGTCTCGCGTGGTGTCTTCCGCATCAAGGGCGACAGCGAGTGAGCGAGGTCATTGACGGAG
TCGCAACGCTGTCTGATCTCGCGTGCCGGCTTACATGGCGATAGCTAGACTGGGCGGTTTTATGGACAG
CAAGCGAACCGGAATTGCCAGCTGGGGCGCCCTCTGGTAAGGTTGGGAAGCCCTGCAAAGTAAACTGGA
TGGCTTTCTTGCCGCCAAGGATCTGATGGCGCAGGGGATCAAGATCTGATCAAGAG**ACAG**ATTAGCAGG
TGTGCATGATCTACGTGCGTCACATGCAGTAC

pNC903

CACTAGCTCAGATTCAGTAGACCGCTGTTGCACCTGCATTA**TACA**CTGGCTGTGGTCCAGCAGCAGCTG
GCACCCAAAGCACCACACACCTCATGCACTAAAGCTGCGACCACGAAGAACGGGATGGTCCGGGTGCGA
CGGAAGTGTGTTTTCCCAGAAGAGTCGCCGAAACATCTGACTTGGTTGGCGTGTCCTACCTAAAAAAAT
TGATCTTGCGTGTGAGGGTGTCACGCATGGATATGCGCGGGGGATCTCTCAGTGGGGACTGGGAGCAGT
TGTGGCTGCCTCTATGGCCGCTCGCAACGGACGATTTGTTGCTCGGGGTCTACAGGATGCCTCGCCAGG
ACGCGCTCGATCGGCGCTATCTGGAGGCCAATCCACAGGCTCTGAGCAATCTCCTCGTCGTCGATGTCG
ATCATCCGGACGCGGCACTGCGGGCCCTGTCTGCTGCCGGCAACCACCCCTTGCCGAACGCGATCGTGG
AGAACCCACGCAATGGACACGCACATGCCGTGTGGGCCTTAGCCGAACCCTTCACGCGCACCGAGTACG
CCAGGCGTAAGCCCCTCGCTTACGCCGCTGCGGTAACCGAGGGGCTGCGTCGAGCTGTCGATGGCGATG
CTGCCTATTCGGGGTTGATGACGAAGAACCCGACTCACTCGGCCTGGGACACACACTGGATCCACGCCG
AGACTCGATCGCTGGCAGACCTCGAACACGACCTCGGCAAGCACATGCCGCCACCCCGGTGGCGGCAGA
GCAAACGTCGTCGCGAAAACCCAGTCGGACTCGGACGCAATTGCATGCTCTTCGAAACGGCTCGTACTT
GGGCATACCGCGAATTGCGTTGCCATTGGGGAGATCCCGAAGGATTAGGGAAAGCGATTCATGTCGAGG
CTGCAGACCTTAACGCCTCCTTCTCCGAACCTTTGCCGGTGAGCGAGGTACGAGCCATCGCAGCAAGCA
TTCACCGCTGGATCGTCACCAAGTCCCGCATGTGGGCCGACGGCCCTGCGGTTTACGAAGCCACCTTCA
TCGCTATCCAATCCGCTCGCGGACGCAAGATGACGGAGAAGAAGCGCGAGGCCAATCGTCGCCGTGCAA
CGAAGTACGACCACGACCTCGTGAGGAAGGAGGCGACCGATGGGAGCTGAAACGCCGGCCCGGCGAAAC
CGCACAGCTCGCGAAGTGGCGGAACGAATCGGTGCGTCCCCGCGCACGGTGCGGCGCATCATCGCGGAG
CCTCGAGCTTCGTACGAGGCTCGCGCAGCTGAACGTCGAAAGCGAGTGCTCGAACTCCGTGCGAACGGG
ATGAAGCTGCGTGAGATCGCGGCGGAGGTAGGTATGTCGGTCGGAGGAGTAGGGACGATCCTGCATCAC
GCCCGTAAGGCCGAGCAGTCCAAGGCAGAAGGAGCTATGGCATGAACGACGCCATATCGGCCCGCATCA
CTGCAATGCAGACCCAACTGACTGCTGTACACACCGAGCTACGTGCTCTAGCGGAGCTGGTGGACATGC
TTGATGCCGACACCCTCGACGCCGAGACTGAAGACTCGGTACGCGAAGTGATCGACTCCCTGGCAGACG
CTGGGCGAGCTTTAGCCGGCGCCGACGAGCCGCTCCAGGCCGCAGTTCATCACGCCCGACGACTGCCTT
AGTCAGCCTCTGTCCGATC**ACAG**ATTAGCAGGTGTGCATGATCTACGTGCGTCACATGCAGTAC

pOTS

CACTAGCTCAGATTCAGTAGACCGCTGTTGCACCTGCATTA**TACA**CGGACGTCATCGACCAGGAACTCG
AGGCGCTCTCGGACGAGACGGGCCGCTCCGTCTCCAAACTGATCCGCGAGGCCGTGACCTCGTTCATCG

CAGACGCGAAGAAGGAAGCGAGCTAACAAGCCCCAGAAATGCCAAACCCCCCGGCTGCCACCGAGGGGT
CCGACAATTGAAGATCAGATACACGCTCGAGCAGGCGACCCACCAAGCGGCATCCAGATTGTCCATCGC
CGATATTACAGGGTCGCCGGTAGCTCGACTAGTCATACGTGCCTGCCAGCATCGCCGTGTCTCCGGTCT
TCTTCTCACAACGAGAGGAATTACCCACAATGGTGACGACCCACAGCTCCAATGAGCCGATCCTGACCT
GGCTATCCGCTGTACGCAGCAAGGCCAGCGACCTCCCGCACAGCGCACGGTCCGTCGCGCTCGCAATCC
TCACGCACGTCCACCCTGAAACGGGAATGACACCGCCGCTCAGCTACGCGGCCCTGCAGCAGGAGGCCG
GTTGCTCTCGCGGCACCCTGGTCGCCGCCCTGCGCCGCCTCGACGACGCCGGGATGATCCGTCGATTCG
GCGCCCACCAGCAACTGAACGTCTACTCGTTGGCTGTGGATAACTCGCTGGCCAGCGCTGTCGAACCCG
GTACGGAATTCGCACCGACCCCGGTACAGCCGCTGACCGCTGCGGACCACGTCGGCAGCGATTCCGCAC
CGACCCCGGTCCAAATTTTGCACCACGAGAAAGAGCTTTTCTCTTCCTCTTCTCCTTCTCTCTCCTCGG
TTGACCCCGATGCCCTGGGGACCGCTGCACCGATCCCGCTGCCGCGGGATTGGGCACCGAATCCCCAGC
ATCTGAGCCGTGTCGCCGCACGAGGCCTCGATCTCGCGGCAGTGGTCGAAGCGTTCCGCCAGTACGCGG
ACGGGAAGACTCGCACGGGCGTGGACAAGTGGGACAAGGCGTTCGGCTGGTGGCTCAAGCCCGGCGAGG
GCTGGGACAAGCACGTCGCCACACGGGCCGCAGCCGCCACTGCGCCATCGGAGCAGGCCTCTGCCAGCG
AGGAAACGCCCGTCAAGTGCACGAACCACCGTGGGATCAGCACCGTAATGCGCACACTGCAAGCCCAGG
GCATCTCTGGCGCACGTGTTAGGGATCGGGTGGTCGAACTCCTCGACGAGGGCACCCCGGTCTACGAGA
TCGCGGGCATGATCGAGAACGAACTCTTCGCGCTACCTGTGGGGGCCTGACACCCGCGACGAAGGAGGG
GGGCTCGCGTGAGGTACTCGGCGATCGACGCCGCCAGAGTGTGGGTGGGGGAGCAGCTGCGTTACCCGC
AGCGGATCC**ACAG**ATTAGCAGGTGTGCATGATCTACGTGCGTCACATGCAGTAC

pPRH

CACTAGCTCAGATTCAGTAGACCGCTGTTGCACCTGCATTA**TACA**CTCAGCATTCAAGCCGGTTTCCGG
CGTGGCGAACAGCCAAAAGGGCCGATTCCACTATGAGCCTGTTGCGATATGGCCGTCGGGGGAGCTGGA
GAAGGGCGCGGAGCCCCAGCGGAGCGCCCTGGTCAAACCGTGCGGGGACCCTTGGGGACCACCAATATC
TGATAAGGGTGTGTTCCTACCTGAGGGATGAGGGGTAGGCCGATACCGTCATCCTGTGATGATGACGGC
GGAGCGGTGGGCTGAGCACTACCGGCAGAAGTGGCCCATGGCCACTGACCATGACAAGGGGCGGTTCCC
TGTCCGCCAGTCCCGCGCCGACGCTCTCAAGCGTCGGTACATCCAGGCCAACCCCGCAGCCTTGACCAC
GCAGATCGTCATTGACCTCGACCATGAGGACTCCCTCGGGCTGGCGTTGGAGCTGAACGGCGTCCCGAC
GCCGAACTATGTGGCGCAGTCGCCCTCCGGCCACGCCCACGTGGCCTACCTGCTGGCGGCTCCCGTCTG
CCGCACGGACAATGCGCGGCTGGAGCCGATGAAGTTCGCGGCCCGCGTCGAGCGGGGTCTGGTCAATGC
GCTACGCGCTGACCTCGGTTACGCGGGCTTCATGACCAAGAACCCCATCCATGACGGCTGGGATACGGT
CTGGACCAATGACCACCTGTGGACGCTGGGAGAGCTCGCCACCCAGCTTTCCGGCTGGCTACCCCGCAG
CCTGCCCCGCCGTGCAGCGGACAACTCCGGCCTGGGCCGCAACGTGGCCCTGTTCAACGACGTTCGGCT
GTGGTCCTACCGGGCGATTCGGAAGCACTGGGAAGCGGGACCGGATGCTTGGGAACAGGCCACCTACGC
CTATGCGCTCGCCGTGAACCACAAGTTCGCGGTGCCCCTGGACGCCTCGGAGGTGGTCCACTTGGCCCG
GTCGGTGTCGCGGTGGACGTGGCGGAACTTTACGCCCGAGGGATTCAGCAAAGTGCAGACACGGCGGAG
CCAGAAGGCTGCCGCCGTCCGCACCGCCGAAAAGATCGTGAAAGCCGAGTTGATTAGGAGCCTCGTCTA
ATGAGCGCACTAGATCCCCGCCGCCGGAAGATCACCGCTCGAGAAGCAGCCGAGCAGGTCGGCTGCACC
CCGCGCCATATCCGGAGCGTGGTTGCCGAGCCTCGACATGAATTCCTGGCGAGAGCGGCTGAGCGCCAG
CGTAAAGCCGCGGACTGGAAGGACGAGGGCCTGACCTACCGGGAGATTGCGGAGCGGCTGGATTGCACC
CCCAAGGCCGCTGAGAACCTGGTTCTGCGTGGCCGCAAGGCCCGCAAGGTGACGGCCTGATGGCCACCT
CCACCGCTCCCACGCTCGGCCTCGCCGAGGCCGCCGCAGCTTGCGGGGTGTCCCTTTCCACCCTTCGCC
GTCGACGGGATGACCTGCGGGCGCTGGGCGCCACGGACGGACCCAAAGGGTGGAGCATCCCCGTGCCGG
CCCTGATCGAACTGGGACTGATGGACC**ACAG**ATTAGCAGGTGTGCATGATCTACGTGCGTCACATGCAG
TAC

pGA1

CACTAGCTCAGATTCAGTAGACCGCTGTTGCACCTGCATTA**TACA**ACGTTTAACAAGGTAGTTAAGCGT
TCATTTACGAAGAAAACACGATAAGCTGCACAAATACCTGAAAAAGTTGAACGCCCCGTGAGCGGGAAC
TCACAGGGCGTCGGCTAACCCCCAGTCATCAGCTGGGAGAAAGCACTCAAGACATGACTCTAGCCGATC
CGCAGGACACAGTCACAGCTAGCGCGTGGAAATTTTCCGCCGATCTGTTCGACACCCACCCCGAACTAG
CGCTGCGCTCACGCGGCTGGACGGCAGAAGATCGCCGCGAACTGCTCGCTCACCTGGGACGCGAAAGCT
TCCAGGGCAGCAAGACAAGAGATTTCGCGAGCGCCTGGATTAAAAACCCGGATACCGGCGAAACCCAAC

CAAAGCTCTACCGGGCTGGCTCAAAAGCGCTGACGCGGTGCCAGTACGTTGCGCTGACGCACGCGCAAC
ATGCCGCGGTGATCGTGCTTGACATCGATGTGCCCAGCCACCAGGCCGGCGGGAAGATTGAGCACGTAA
ACCCGCAGGTCTACGCGATTTTAGAGAAATGGGCACGCCTAGAAAAAGCGCCGGCTTGGATCGGCGTGA
ATCCGCTGAGCGGGAAATGCCAGCTCATCTGGCTCATTGACCCGGTGTATGCCGCAGCAGGTAAAACCA
GCCCAAATATGCGCCTGCTGGCTGCAACGACGGAAGAAATGACTCGTGTTTTCGGCGCTGACCAGGCTT
TTTCGCATAGGCTGAGCCGGTGGCCGCTGCACGTCTCAGACGATCCGACAGCCTATAAATGGCACTGCC
AGCATGATCGTGTGGATCGGCTGGCCGACCTAATGGAGATTGCTCGAACGATGACCGGATCACAGAAGC
CGAAAAAGTACATTGAGCAGGACTTTTCCAGCGGACGCGCCCGCATTGAAGCGGCACAACGCGCCACCG
CAGAAGCCAAGGCGCTAGCGATTTTGGACGCGAGCCTGCCGAGCGCCCTGGACGCGTCCGGCGACCTGA
TCGACGGCGTGCGAGTGCTCTGGACAAATCCAGAGCGAGCGCGCGACGAGACCGCGTTTCGCCACGCGT
TGACCGTGGGATACCAGCTCAAAGCTGCTGGTGAGCGCCTAAAAGATGCCAAGATCATCGACGCGTATG
AAGTGGCGTACAACGTTGCCCAGGCGGTCGGTGCAGACGGCCGGGAGCCGGATCTTCCCGCCATGCGTG
ATCGCCTGACGATGGCGCGTCGTGTGCGCGGCTACGTGGCTAAAGGCCAGCCAGTCGTCCCTGCTCGTC
GGGTGGAAACGCAGAGCAGCCGAGGGCGGAAAGCTCTAGCGACGATGGGGCGACGGGGCGCAGCTACAT
CGAATGCACGCAGATGGGCTGACCCAGAAAGTAAGTATGCGCAGGAGACGCGACAGCGATTAGCGGAAG
CAAACAAACGCCGAGAAATGACAGGCGAGTTGCTCGAACTTCGCGTCAAAACTGCGATCCTGGATGCCC
GTTCTCAATCGGTTGCTGATCCCTCGACTCGTGAGCTTGCAGGCGAACTAGGTGTCAGTGAAAGGCGCA
TCCAACAAGTCAGAAAGGCACTTGGAATGGAAGCTAAACGCGGCCGTCCACGGGCTGAAAACTAATAAA
CGAAACACCGTCAGCAGAAAACGGTTCCCCCCTTTAGGGGTCCCGTCCTTGCTCTGGCTCTCACTTGCC
CTCACCCTCCGCTATCCACGGGCTGAAAACTAATAAACGAAACACCGTCAGCAGAAAACGGTTCCCCCC
CTTTAGGGTGTCTCGCTCCTAGCTCTGATCCCTCCCCGGTTCCTCCCCGGCCTGATTTTTAAGGGGGGC
TCACGCTGTCGGCAGAGAACGGTTCCCCGCCTTCTGCTCTGGCTCTTCCTCGACTCCCTCCCCCTCAAA
AATCTCCTCGAG**ACAG**ATTAGCAGGTGTGCATGATCTACGTGCGTCACATGCAGTAC

pNG2

CACTAGCTCAGATTCAGTAGACCGCTGTTGCACCTGCATTA**TACA**ATGGTAAATCTGCGCAGACAGCCC
TGTGCAGCTGAAACGCGGTTACGTATAGCTTGCCATATGTCTAGCCATACGTAACCGCAGGTAAAAGGC
ATATTTTTCGCGTGTCATGGCTAGTAAATAACACCGGTGTCATTTAGAGTCAGGGAAAGACAATGAAAA
ACGAAGAAAGCCACCGGGCGGCAACCCGATGACTTTCGCTTATCACCCAGCACACACCTGGGAGAAATC
ACGGTCATGAGTTTACAGACTCATGCGCAGAATGCGCACACTAAAACACCTACCCGCGTCGAGCGCGAC
CGTGGTGGACTGGACAACACCCCAGCATCTGCCAGTGACCGCGACCTTTTACGCGATCATCTAGGCCGC
GATGTACTCCACGGTTCAGTCACACGAGACTTTAAAAAGGCCTATCGACGCAACGCTGACGGCACGAAC
TCGCCGCGTATGTATCGCTTCGAGACTGATGCTTTAGGACGGTGCGAGTACGCCATGCTCACCACCAAG
CAGTACGCCGCCGTCCTGGTCGTAGACGTTGACCAAGTAGGTACCGCAGGCGGTGACCCCGCAGACTTA
AACCCGTACGTCCGCGACGTGGTGCGCTCACTGATTACTCATAGCGTCGGGCCAGCCTGGGTGGGTATT
AACCCAACTAACGGCAAAGCCCAGTTCATATGGCTTATTGACCCTGTCTACGCTGACCGTAACGGTAAA
TCTGCGCAGATGAAGCTTCTTGCAGCAACCACGCGTGTGCTGGGTGAGCTTTTAGACCATGACCCGCAC
TTTTCCCACCGCTTTAGCCGCAACCCGTTCTACACAGGCAAAGCCCCTACCGCTTATCGTTGGTATAGG
CAGCACAACCGGGTGATGCGCCTTGGAGACTTGATAAAGCAGGTAAGGGATATGGCAGGACACGACCAG
TTCAACCCCACCCCACGCCAGCAATTCAGCTCTGGCCGCGAACTTATCAACGCGGTCAAGACCCGCCGT
GAAGAAGCCCAAGCATTCAAAGCACTCGCCCAGGACGTAGACGCGGAAATCGCCGGTGGTCTGGACCAG
TATGACCCGGAACTTATCGACGGTGTGCGTGTGCTCTGGATTGTCCAAGGAACCGCAGCACGCGACGAA
ACAGCCTTTAGACATGCGCTTAAGACTGGCCACCGCTTGCGCCAGCAAGGCCAACGCCTGACAGACGCA
GCAATCATCGACGCCTATGAGCACGCCTACAACGTCGCACACACCCACGGCGGTGCAGGCCGCGACAAC
GAGATGCCACCCATGCGCGACCGCCAAACCATGGCAAGGCGCGTGCGCGGGTATGTCGCCCAATCCAAG
AGCGAAACCTACAGCGGCTCTAACGCACCAGGTAAAGCCACCAGCAGCGAGCGGAAAGCCTTGGCCACG
ATGGGACGCAGAGGCGGACAAAAAGCCGCACAACGCTGGAAAACAGACCCCGAGGGCAAATATGCGCAA
GCACAAAGGTCGAAGCTTGAAAAGACGCACCGTAAGAAAAAGGCTCAAGGACGATCTACGAAGTCCCGT
ATTAGCCAAATGGTGAACGATCAGTATTTCCAGACAGGGACAGTTCCCACGTGGGCTGAAATAGGGGCA
GAGGTAGGAGTCTCTCGCGCCACGGTTGCTAGGCATGTCGCGGAGCTAAAGAAGAGCGGTGACTATCCG
GACGTTTAAGGGTTCTCATACCGTAAGCAATATACGGTTCCCCTGCCGTTAGGCAGTTAGATAAAACCT
CACTTGAAGAAAACCTTGAGGGGCAGGGCAGCTTATATGCTTCAAAGCATGACTTCCTCTGTTCTCCTA

| |
|---|
| GACCTCGCAACCCTCCGCCATAACCTCACCGAATTC**ACAG**ATTAGCAGGTGTGCATGATCTACGTGCGT<br>CACATGCAGTAC |
| pRET1100 |
| CACTAGCTCAGATTCAGTAGACCGCTGTTGCACCTGCATTA**TACA**GGATCCGTCATGACTGGACCACAG<br>GAGAGAAAGCGCAAGGCGGCGAAGCCGTCGCGGGAGCCTCAGTTGAACTGCTGTGAAGCGGACGTGCCG<br>AAACGAGCAAAACAGCCCCCGGTTCCCTCTACGTTCGACCTGCTCACGGTGAAGGAGACTGCGGGGCTG<br>CTGAGAGTCAGTCAGGCAACTCTTTACCGGCTGCTTCGGAGTGGGGAAGGACCCACATACACACGGATC<br>GGTGGACAGATACGCGTTCACCGCGAGTCGCTGCGTCGGTTCATCGAACCGCGTGGATAACGTCACAGA<br>GACAGCGAAAACGCCTCCCCTGGGTCAATCCGGTTACCGCCGGACTGGGGGAGGCGCTTCGACACCTAC<br>ATCCGTCGCCCCTCGAAAGGCTCAGATGCACTTCCACGATAACGCAGAGGTCGGACAAGAGGGAAGAAC<br>TGCCGTTCTCTCGCCGTTGCGCGGCGTAGCCGCCAAGCGGGACGTGTCTGACGATGCAGCGAAGCGGAG<br>TCGGCAGGCGCGGCACGCGCCTGGGCTTGTTACATCTGCCACAACTGTCCGTGAATCTCTGCCAGCTCC<br>TGAAACCGCTGGTCAGGGCCTTGCGGAATCCGTGACCGCTGATGATTTTTGGTCCCATTCGTTCCCCCG<br>CGCTGACGATGTACGCGGCGCAGCTGCTTCCTTCCAGTCGGTGGCTAACTGGGATGGGCGTGAGGGTCC<br>GAGGCCGCGTTTCGTTGTCGCGCCTGGCGTTGTCCGCTTGGAGGTTTGTGATCTCGCACGCCGCGAACG<br>AACGGCTGAACGTGCGTATCTGGCTGCTCGGGCTCGGGTGGATATGGCGGCTGCCAGGCATAACTCGCC<br>GTACGACTTCGACGTGGACGATGAAGAGTTGGCGGAACTGGCTTCTCTGCAAGGCCTCGAGGACGACGA<br>CATTGGGGGCTGGTCTGCGGAGAGGGAAATAGTGGGCTGGTCTGCTCGTTCTCGGTCACGGATGATCTT<br>GCGAATGGCAGAACTCGACTGGGCTCCCATGATGGATTTGCCGGGCATTCCTGCGATGGTGACCCTCAC<br>CTATCCGGGGGACTGGCTTACGGTTGCCCCCACCGGCGCTGAGGTCAAAAAACATCTCCAGACGTTCTT<br>CAAACGGTTCCAACGGGCCTGGGGCATTGCCTGGATGGGTGCGTGGAAAATGGAGTTCCAAAGCCGAGG<br>CGCTCCGCATTTTCACCTGTACATGGTCCCTCCTCATGGGAAGGCAGGAGACTCGCGGAAGCTGCGGCA<br>TGATGCTGAGCTCTTGAAATGGGAGATAGCACGTGCAGAGGGTGAAGACCCAGGTCGCAGGCCGTATTT<br>CCGGGAAGCTCCAAGCGATGGATTGAAGTTTCGTCCGTGGCTTTCTGCGGTGTGGGCCGACGTCGTAGA<br>TCATCCGGACCCCAAGGAAAAAGAAAAGCACGTCAGTGCCGGCACTGGAGTGGACTACGCGGAGGGCAC<br>GCGAGGGTCAGATCCGAAAAGGCTTGCGGTGTACTTCTCCAAGCATGGAACCTTTGCCGACAAGGAATA<br>TCAGCACGTAGTTCCTGCTCAATGGCAGAAAACGGGTGCGGGACCTGGCAGGTTCTGGGGCTACCGCGG<br>TTTGTCGCCGGCCACGGCTGCCACCGAGATTTCCTGGGATGAGTACCTGCTTTTATCTCGCACGTTGCG<br>ACGATTGTCAGCGCGAACGAAGATCTGGGACCCGGCTTTACGAGGCGGTAGCGGCGGCCACAGATGGAC<br>TAAGGCGATGATGCGACGCACGGTTACCCGGCACCGCTTGGACCTCGTGACCGGTGAGATTCTGGGCAC<br>GAAGACGCGGAAGGTTCGGGCGCCAGTGAAGAGGTTTGTCCGGACTTCGGGATACCTGTGTGTCAATGA<br>CGGGCCCGCACTGGCTCGAACCCTCAGCCGTCTTCGTACAAGCTGCCTGAGCTAGACGCGCGGAACGCC<br>TTTCGGCTTTGTCTTTTGCTGGATGGCGGGTTTTGGGCGGCTTCTGGTGATGCGCTGCTGCGCTCCGTG<br>GGGAGAGAGTCCCAACGACTGACCTATCTCTACCCAGGTGCAATTCATCTCCCGCGCTCTGTCGGCTAG<br>GTAAACGAGGTGCTCCCGCGCGAGCTTTTCCATGTGGTCGGCCAATGTCAGCTCGGTCAGGACAACCTG<br>CTGTTGTTGCGATAGTTGTGTCCGCACGGGTCGATTGTCTTCTGTTGCGGCATAACGGTTTTCGTCGTT<br>CGCGGAGAGTGCGGCTAAATGAATTGCATCCTCGATTGAGCGGAGCATTTCGACGCGGAACCTGGCGA**A**<br>**CAG**ATTAGCAGGTGTGCATGATCTACGTGCGTCACATGCAGTAC |
| pSR1 |
| CACTAGCTCAGATTCAGTAGACCGCTGTTGCACCTGCATTA**TACA**GTACTGCGGCGTCGCTGATCGCCC<br>TGGCGACGTTGTGCGGGTGGCTTGTCCCTGAGGGCGCTGCGACAGATAGCTAAAAATCTGGGTCAGGAT<br>CGCCGTAGAGCGCGCGTCGTCGATTGGAGGCTTCCCCTTTGGTTGACGGTCTTCAATCGCTCTACGGCG<br>ATCCTGACGCTTTTTTGTTGCGTACCGTCGATCGTTTTATTTCTGTCGATCCCGAAAAAGTTTTTGCCT<br>TTTGTAAAAAACTTCTCGGTCGCCCCGCAAATTTTCGATTCCAGATTTTTTAAAAACCAAGCCAGAAAT<br>ACGACACACCGTTTGCAGATAATCTGTCTTTCGGAAAAATCAAGTGCGATACAAAATTTTTAGCACCCC<br>TGAGCTGCGCAAAGTCCCGCTTCGTGAAAATTTTCGTGCCGCGTGATTTTCCGCCAAAAACTTTAACGA<br>ACGTTCGTTATAATGGTGTCATGACCTTCACGACGAAGTACTAAAATTGGCCCGAATCATCAGCTATGG<br>ATCTCTCTGATGTCGCGCTGGAGTCCGACGCGCTCGATGCTGCCGTCGATTTAAAAACGGTGATCGGAT<br>TTTTCCGAGCTCTCGATACGACGGACGCGCCAGCATCACGAGACTGGGCCAGTGCCGCGAGCGACCTAG<br>AAACTCTCGTGGCGGATCTTGAGGAGCTGGCTGACGAGCTGCGTGCTCGGCCAGCGCCAGGAGGACGCA<br>CAGTAGTGGAGGATGCAATCAGTTGCGCCTACTGCGGTGGCCTGATTCCTCCCCGGCCTGACCCGCGAG |

| |
|---|
| GACGGCGCGCAAAATATTGCTCAGATGCGTGTCGTGCCGCAGCCAGCCGCGAGCGCGCCAACAAACGCC<br>ACGCCGAGGAGCTGGAGGCGGCTAGGTCGCAAATGGCGCTGGAAGTGCGTCCCCCGAGCGAAATTTTGG<br>CAATGGTCGTCACAGAGCTGGAAGCGGCAGCGAGAATTATCGCGATCGTGGCGGTGCCCGCAGGCATGA<br>CAAACATCGTAAATGCCGCGTTTCGTGTGCCGTGGCCGCCCAGGACGTGTCAGCGCCGCCACCACCTGC<br>ACCGAATCGGCAGCAGCGTCGCGCGTCGAAAAAGCGCACAGGCGGCAAGAAGCGATAAGCTGCACGAAT<br>ACCTGAAAAATGTTGAACGCCCCGTGAGCGGTAACTCACAGGGCGTCGGCTAACCCCCAGTCCAAACCT<br>GGGAGAAAGCGCTCAAAAATGACTCTAGCGGATTCACGAGACATTGACACACCGGCCTGGAAATTTTCC<br>GCTGATCTGTTCGACACCCATCCCGAGCTCGCGCTGCGATCACGTGGCTGGACGAGCGAAGACCGCCGC<br>GAGTTCCTCGCTCACCTGGGCAGAGAAAATTTCCAGGGCAGCAAGACCCGCGACTTCGCCAGCGCTTGG<br>ATCAAAGACCCGGACACGGAGAAACACAGCCGAAGTTATACCGAGTTGGTTCAAAATCGCTTGCCCGGT<br>GCCAGTATGTTGCTCTGACGCACGCGCAGCACGCAGCCGTGCTTGTCCTGGACATTGATGTGCCGAGCC<br>ACCAGGCCGGCGGGAAAATCGAGCACGTAAACCCCGAGGTCTACGCGATTTTGGAGCGCTGGGCACGCC<br>TGGAAAAAGCGCCAGCTTGGATCGGCGTGAATCCACTGAGCGGGAAATGCCAGCTCATCTGGCTCATTG<br>ATCCGGTGTATGCCGCAGCAGGCATGAGCAGCCCGAATATGCGCCTGCTGGCTGCAACGACCGAGGAAA<br>TGACCCGCGTTTTCGGCGCTGACCAGGCTTTTTCACATAGGCTGAGCCGTGGCCACTGCACTCTCCGAC<br>GATCCCAGCCGTACCGCTGGCATGCCCAGCACAATCGCGTGGATCGCCTAGCTGATCTTATGGAGGTTG<br>CTCGCATGATCTCAGGCACAGAAAAACCTAAAAAACGCTATGAGCAGGAGTTTTCTAGCGGACGGGCAC<br>GTATCGAAGCGGCAAGAAAAGCCACTGCGGAAGCAAAAGCACTTGCCACGCTTGAAGCAAGCCTGCCGA<br>GCGCCGCTGAAGCGTCTGGAGAGCTGATCGACGGCGTCCGTGTCCTCTGGACTGCTCCAGGGCGTGCCG<br>CCCGTGATGAGACGGCTTTTCGCCACGCTTTGACTGTGGGATACCAGTTAAAAGCGGCTGGTGAGCGCC<br>TAAAAGACACCAAGATCATCGACGCCTACGAGCGTGCCTACACCGTCGCTCAGGCGGTCGGAGCAGACG<br>GCCGTGAGCCTGATCTGCCGCCGATGCGTGACCGCCAGACGATGGCGCGACGTGTGCGCGGCTACGTCG<br>CTAAAGGCCAGCCAGTCGTCCCTGCTCGTCAGACAGAGACGCAGAGCAGCCGAGGGCGAAAAGCTCTGG<br>CCACTATGGGAAGACGTGGCGGTAAAAAGGCCGCAGAACGCTGGAAAGACCCAAACAGTGAGTACGCCC<br>GAGCACAGCGAGAAAAACTAGCTAAGTCCAGTCAACGACAAGCTAGGAAAGCTAAAGGAAATCGCTTGA<br>CCATTGCAGGTTGGTTTATGACTGTTGAGGGAGAGACTGGCTCGTGGCCGACAATCAATGAAGCTATGT<br>CTGAATTTAGCGTGTCACGTCAGACCGTGAATAGAGCACTTAAGTCTGCGGGCATTGAACTTCCACGAG<br>GACGCCGTAAAGCTTCCCAGTAAATGTGCCATCTCGTAGGCAGAAAACGGTTCCCCCGTAGGGTCTCTC<br>TCTTGGCCTCCTTTCTAGGTCGGGCTGATTGCTCTTGAAGCTCTCTAGGGGGGCTCACACCATAGGCAG<br>ATAACGTTCCCCACCGGCTCGCCTCGTAAGCGCACAAGGACTGCTCCCAAAGAGCTTCAAAGCCACTGC<br>CGCGACTGCCTTCGCGAAGCCTTGCCCCGCGGAAATTTCCTCCACCGAGTTCGTGCACACCCCTATGCC<br>AAGCTTCTTTCACCCTAAATTCGAGAGATTGGATTCTTACCGTGGAAATTCTTCGCAAAAATCGTCCCC<br>TGATCGCCCTTGCGACGTTGGCGTCGGTGCCGCTGGTTGCGCTTGGCTTGACCGACTTGATCT**ACAG**AT<br>TAGCAGGTGTGCATGATCTACGTGCGTCACATGCAGTAC |
| pXT107 |
| CACTAGCTCAGATTCAGTAGACCGCTGTTGCACCTGCATTA**TACA**GTTGGATTTGGCGGCAGGATCGGG<br>AAACCGTACCGCCTCACCATCGGGCAGGCACCCCGTGAAATCGCCCCGGCCACACCGGTCGCGCTCCTC<br>GAAAAGGCCCAGGACGAGAACTAGTCTCACACCGATGTAATTCGGGCTCCCCCGAAACAAAGTCGAGGG<br>CGGACCACTCCAACTGGTCCGCCCTCTGGCTTTCACCCGGCAGAACCTCTCACAGCCACTCGCCGGGCG<br>GGGGTGCTCCCACTGAAGGAGACCGATCGATGGTATCAACCGAACCCGTCATCGAGACGACGGACAACC<br>CGAGCTGTGTCGTGTGCGGCTCGGAGCTCCCCCGGCAGAAGAAGGGCCGACCCCGTCGGTATTGCAACC<br>CAGCATGTGCCCGTCGAGCACGCCAGGCGTCTCCCTCCAACCAGGCATCCGAAGGTCCCTCCCATACGC<br>CCGCACCCGGTCGTCGCGGAGCAGAGGCAAGGGGGCAGGGCGTCGGAATAGAACAGGCACCAAGCGATG<br>CACTGGATCCCCCGGCAGACGCCGTGGAAGACTTTCGTACTCGGTACTACGCAAACTTTGCGTCCACCC<br>CAGGTCAGGACCCGATTTCAACACGGCGGAAACTCCGATACCGCCTCCGGCACCTGCTCCGGGAAGTCA<br>CGACCGTCAACCGATGCAAGGGCTGCGGATGGGAACCCATCGCCAACGGCATCGCCATCAAGATGGGCA<br>CCAAGGACGGCCGGAACACGGCAGGCTTCGGAGGACTGGAGACCTGCGGCCGGATCTGGCTCTGCCCGG<br>TCTGCTCGGCCAAGATCCGCGTTCGCCGCGGCGACCAGATCGCAGAAGGAGTTGGCCGGCATATCGACG<br>GCGGCGGATCAGCCTGGTTCCTCACCATCACCCTGCCGCACGAGAAGGGCGACGCCCTCAAGGACTCGT<br>TCGACGCGCTCACCCAGGCCTGGCGCTACGTCAAAACCGGCCGCGCCTACCAGGACGAAAAAAAGTGGT<br>TCGGGATCCTCGGCGAGATCAAGGCCGTGGAGGTCACCCACGGTGACAACGGCTGGCATCCACACGCCC |

ACATCCTGATCTTGACCAGCCGAAACCTGGAATTCGACGAGTTCTGCTCCTGGGTCGCTCGACTCGACG
CTCGATGGGCCAAGGGACTGGCCAAGGTCGGATGGTCCACCGGCCGCCCCGGAATCCGCCTCGTCCTGG
TCCCCGTCGAGAAGGGCAAGGGCATCGCTGCCTACGTCGCCAAGGTCCAGGAGAAGGGCCTGGGCAACG
AGATCGCCCGCGCCGACATGAAGGACGCCCGGCCCGGCAACCGGACACCGTTCGGAATTCTCGCCGACT
TCGGGAACCACGCCCTGGCCGACGACGTCGAACTCTGGTGGGAGTACGAACAGGCCACTGCGGGAAGGT
CAGCAATCCGATGGAGTCCGGGCCTCTGGGCCAAGCTCATGCCGGATGCCGACGACCTGAGCGACGAGG
AGATCGCAGCCGAGGACGAAGGCAGCGCCGCAATCGCCTACCTGGCGCCCTGGACCTGGCGTCGTATCC
GAACCATCCCCGGGGCCGAAATCGCCATCCTGGAAGCCGTAGAGCGGTCCGGATGGGAGGGCCTACTGC
GGGTGCTCATCCGTTACCGCGTGAGCGCTGCGGGCGTCTACACCCCGGAGGAATGGGCCACCCCGAACG
AGGTCGAACTCGAACCGCTGTAGAACCGCGACCCGGGTGCAAGAGGTGTACACCGAAAAGGCCGGCCGA
ATCGCGCTGGGTCCGGGCACACCTCTTGCACCATCGCTCGGACCAGCCTCCTCCCTTCCCCGATATAGG
CCATCTCCAAAGCCATGGAGAGGCAGTCAAGGGCGCTCTTTCCCTTGACTTCCTCGGAAGGGCTGGACA
ATCGAGGCCAGGGGAAGACGGGCAACCCATCAAGATGATGCCGACGTATCGAGATGATGCGAACGC**ACA**
**G**ATTAGCAGGTGTGCATGATCTACGTGCGTCACATGCAGTAC

pYS1

CACTAGCTCAGATTCAGTAGACCGCTGTTGCACCTGCATTA**TACA**ACATCTCGGCACATTGCCCGACTC
CCCTGCGGCGTTGCTCGCACCGTCGGACGTCGACGAGAACTAGCCTCACCCCCGCATAGCACCCCGAAA
CGCACTGAGGGCGAACCAATGCACTGGTTCGCCCTCGGGGCTTATCCGGCCTTCTTGCCTAGCCAGCAC
GGGCCGGATGGGTGCTCCCACTGAAGGAGACCGATCGATGGTAGCAACTGCGCACGATAACGGGCCTGC
GGACCGCCGAAGCTGTGTCGCGTGCGGCACGGAACTCCGGTCGGCCAAATCCGGACGACCTCGCCGGTA
CTGCAACGAGGCGTGCCGCAAGGCCTTCTCTCGTACAAAATCCCAGGTCACCCCCTCTTTTGCCCGGAG
GCGGGGAGGCCTTGGAACACCCCGGCGAGAGTTTCCGGACTTGGTACTACGCAAACTTTGCGTCACCCG
CAGGTCAGGACCCCATTTCGGCCCGGCGGAAACTCCGATACCGCCTGCGACACCTGCTCCGAGAGGTCT
CGACAGTCGACCGGGTGAAGAGCTGCGGATGGGAACCAATCGCCAACGGAATCCAGATCAGGCTGAGCG
CCACCGACGGCCGGATGGTCGCGGGCTTCGGAGGCCTGGAGACCTGCGGTCGAATCTGGCTCTGCCCGG
TGTGCAGCGCCAAAGTCCGCGTGCGACGCGGTGATCAGATCGCGGAGGCTTGCGCGCGGCACCTCGACA
ACGAGGGCGCGGCCTGCTGGTTCATCACGGTCACGCTGCCGCACGAGAAAGGCGACGCGCTCAAGGACT
CGTTCACCGCGCTGAATGAGGCGTGGCGGTACGTGAAGACCGGCAAGGCCTACCAGGCTGACAAGAAGC
GGTTCGGGATCCTCGGCGACATCCGGGCCGTCGAGGTTACCCACGGCGAGAACGGCTGGCACCCGCACG
CCCACATCCTGGTGTTCACCACGAAACAGCTCGAATTCGACGAGATGTGTGCGTGGGCGGCCCGGCTCG
ATGCCCGCTGGGCTCGGGGACTGGCCAAAGTCGGCTGGCCGCAGGGCAAACCCGGTGTTCGGTTCACGA
TGGTCCCGGTCACCAAGGGCAAGGGCGTAGCGGCCTACGTCGCCAAGGTGCAGGAAAAGGGACTCGGCA
ACGAGATCGCCCGAGCGGACATGAAGGACGCGACCAGCGGGAACCGCACCCCGTTCGGAATCCTCGCCG
ACTTCGGAAACGACGCCCTGGCCGACGATCTCGAACTCTGGTGGGAATACGAAGCCGCAACCGCCGGCC
GATCCGCGATCCGCTGGAGCCCCGGCCTGTGGGCCAAGCTCATGCCCGATGAGGACGAGCTGACCGACG
AGGAGATCGCGGCCGAGGACGAGAAGGGCTCCACAATCGCTTACATGGCCCCGTGGACGTGGCGGCGGC
TCCGGACTATCCAGGGAGCCGAAATCGCCGTCCTGGAGGCCGTAGAGGCCCACGGATGGGGCGGCCTAC
TCCGGGTGCTCATCGACCTCCGGATCGGTACTCAGGGAGTCTTCACGCCCGACGAATGGGCCACCCCGG
GAACTGTGGATCTCGTCGGCTGACTCCGGATTCACTGATGGTGCAAGAGGTGTAGCGGAGATTGGACCG
TTACCGGGGAGCATGAGGTGTGACCCTCCGCTGCTCCCCCTTCCCCGCTCGGAACTGCCGCCCGGGTAC
ACACCTCTTCCACCATCGGTGACCCCCGTACTCCCTTCCCCGATACAGGCCAGATCCAAAGCACTGGAG
AGGGTGTCAAGGGCGAAGAGCGAAGCGTCCCTTGACTCCATCGGAAGGGCTGGACAATCGAG**ACAG**ATT
AGCAGGTGTGCATGATCTACGTGCGTCACATGCAGTAC

pJAZ38

CACTAGCTCAGATTCAGTAGACCGCTGTTGCACCTGCATTA**TACA**GGATCCTCTAGAGTCGACCTGCAG
GCATGCAAGCTTCACGATTTTTGCAACAGAGCCCTGCGCTCCTTGTACTTCAGGAAACCGTCCACGGCC
TGCTTCGTCACACCGAGTTGGTTTGCGATCCACGTCGTCCCGTGTCCACTGTCGACCAGCCGCCTCGCC
GCCTGCCGGCGCCGCTCTCGTGCGTCCGCGATGTCCTCAGTGAGGCGTTCCAAGTCTGCGTGCGCGCGC
ACCAGCTCTGCGAGCGCGTCGCTTTCATCCACGGTTGCCACCTCACGCTTAAGCGATAACAAAGTCAAG
ACTGTCTTGACAGCCGCTCTCGGCGACCCTACTGTCAAGAGAGTCTAGACGACCCGGTTTCGTTAGAAA
CTTCCGTAACGTTTGAACCCAAAAAGCTGGGCAGTCACCAATGAGCCTTGTCGTCTTTGTAAGAGCTCG

CTCGCGCACACTTTCGCTAATCCGCCGGGCGGCCGGGCATATCCAATCCACCCCGGATTCCTTCTCCTG
GAATGTCGTCGACATTCACGGGGCACCCGTAGCCACTGGCTTGACCAAGGGAGCTGGGCAACGAGTCAC
GCAACTCCGGGTTCGCCGCCGCTCAGTGGTACCAACGAACCGACTGGTACCTGGGTTACGGTCGAGGCA
GAGCTCTCACTTGTGCGGAACGGCTCATCATCGCCGACATCGTCGAATCCTTCAGTAGAGCCGCCGAAC
AGTTCCGTTCTGCAAGCGCACTCCGCCCGGTGCGCTTTCTCGACCAGAGTGCCGACGACGATCGGATTT
GGGACGCAGCGCTTCTCGCCGACGGCCAATACCTGCTCCATGGCGACTACTTCCACACCTACCGTGCCA
AGGATCTTGATTTTCTTACCACAGATCGATCGCGGCGACCAGCGATCTCGCTGCATTCCTGCGCGATCT
TCTTTCAGACACGGCCACGTGCGGTTCAGCCTGAACACGAGCATCTCAGCCTGACGGTAGGGCGATTTG
GGGCGATTGCCGGAAATCTGCCGCGCACACCGGCGTGGCTTCGTGATTTCCGGCAAAGCTACCGGCAGG
ATCGCGCGTATGACCGGATCAACTCGAGCAGGCCGCCCGTCCCTGGGCGAGCGAGCCGTCATGACCGCA
ACACTCCCCGTCACCGTGCTCAATGCAGCAATCGCGACCGCGGCTGACTACGGCATCGACCGGTCCGCC
GTGCTGACCGACATCGTCTGTTTCCACTACGGCCGGCCCGATCTGATGCGTCACCTGCCGCAGCAGCTG
TTGTTTGAATCGCAGCCCGCCACCGCCGATACCGACGACACGATCGGACCGCACGCGAAAGTCCGTCTG
CCGCTACCGATCGCTCAGCTCGTCGAGCGCGACCATCAGCGACTCGGAATGCAGCGCTCCACCTACCTC
GGTGACATCATCTGCCGCCACATGGGCTTCCCGCACCTTGTCCGCGAACTCGACAAGGAGGTTCTGCTG
CTGGCGATGTGACACCGACAGTCCGGCGACTTTTCTTTTGGCTTGTCGTGGTCGACCGCCGGACGGACG
TGACATCTGAATAGCTGTGCAGACCGCCTGACATGACGGAGGCCTCGCCGTAGCGAGGCCCTGTCACAA
GGGTTCGAACCGGGTTGCCGCCCGGTTCAGGTCCGCACCTGTGCCCCTGAAAGGGACGTGTCTTGCTGG
ACTCGTCTCACGGTAGCGCACACCTTGCGCACGTACCAGATACGCCGCGCATCTTGCGTGCAAACCGTG
ACCAAAACGGCGCCGGCGACCGTTGCCGACGACCGTTCACCCCCACACCGCCGACGTACCGCGGCTGTA
GCCCGACCGTGCGCCGGCGCGGTGTCCTGCGCACCACCCGCACCGCTGAGGTGTTGGCGATCCGTCGCG
GCCGGCGACTGGCAGGGCGCGCCATGCCGGCACCGAGCGAGTTCGTCGGCCTGGAGCTCGACGCCGACG
CCTATGCCGGCGTGCCGTGCTGGAGCGGTGGCCCCGCGCACTGGGCGCACGTCACCGTCGCAGTGGCTT
ACGACGTGCACTACGCCATGGTGCGGCCGCGCATGTGCAACGGCGGCATCGCCCGCACCACGTTGATCG
TCATCGCCGCCGCCATGGCGCAGACCGCCGACTGGGATACCGGCCGCAACTGTCGGCCGACCAACGAGC
AGCTCGAAGCGGCCACCGGTTTCGATGAGCGCACCATCCAGGGGGCGCACGAATGCCTGCGCTTGCTGG
GGGTGGCCACCGAGGTCCTCCGTGGCCGCCAACGCACCTACACCGAACGCATGCCGCCGTGGCGAATGG
GCGATCGCCACCGCGGCTGGCCGTCGTTGTGGGCACTGCACGGCAACCCCCACATCGCCCGAGTTGTCC
ACAGCTTGTCACCCCACCTGGAACGGTCTCAAGCAACCACTAAAAACTCACCTTTGAAGAGACTGGTCA
CTACCCAAGGCGGGCGCAAGCGCCCGGCACCCAAGCCTGCGCGCCGCCGAGCACCTGACGAGGCCGGCC
GCCGGCTGGCCACCAGGTGGCGGGCCGACCGGCACGCTCCGCCCTGGGTTCGCACGTATGCCGCCGACA
GCTGGGCGGCGATGTTGGCCGCACCCGCGGCCGCCGGTTGGACGCCGGCCGACCTGACGGCGTTGGTAC
GCGACTGGTTGAGCACCGGTCACTGGGTGCCCGATGTGCCGGCGCGGCCCATCGCTTTGTTGGGCACGA
TGCTGGCCTGGCACACCAGCCACAACAGTCTGGAGGATCGGCCCGCCGCACTCGACGAAGCGCGTGAGG
CTGAAGAACTTGCGGCTGCCCGACGTTGCGTGCGCGACCAGTTCCGAGCACATGACGAATATGCGACCG
ATCGCGCCACTGCCCAAGCTGCCTTAGACGGTCCTGGTCACGCCGCAGCGCGTCAGGCCGTTGCAGAGG
CTGTGCGTCGTGCCGCCTGTAAACGCACCGTCGTCGTCGCGGCCGAAACCGCTCAATTCGCCGCAATGG
TTCAGACTGCCCGTAGCCCCCGCTGATCTACATGAGCCCGCAGCGCTGTTCGTTAGCGAGAGCGGAGCT
ATCTCGATGCCCAGCCAACTCAGAAGGTTCGACTGCACTTGGCTCCCGGTATCCCGTCGAAAAATAGCA
ACCCGCTCGTCGGTTTCACTCGGCTCAATTAGCGGTTGCGATGATGCTTGGTTCTCGACCTCAATCGTG
GGCCCGCCGAAGGCGCAAGGGGCAAGCGATTTCACTTTCCAACACCCCTTGGGTTTGATAAGAATGTTT
TCCCTCGCTTGCCATCTTTTCCCAGACCAGATGCTTTGTTGTTTTGGCCTGTTTTGTCCTCGCCTCCTT
CCCTCGCCTGCCTCCCAGCTCCCAATCCAGTGTCCAGTTCCCCAGCTACTTCCTTGATAACAAGCATTT
AGAGTTCACACTTTGGCACAAGGTGTGCAAGACTGGCGGTAGCACACGTCCGTGTGCTACCCGAAGGCC
CCCGCCCCGGGGCCTTGGGGTTGGGAGCGTGACCGGCTGTGGTGGCTAATGTGACCCCGCTCAAGGCGT
GGGGGATCAAGTACTACGAGGACACTGCTGCCGCTGCCGGCCGGGCCGCGCGCGACACCCGATACGGCA
ACGGCGGGCTCGGTGAGTACTACTCCGAGCGCGACACACGCCAGCCGGTTTGGCTGGTGTCTGGGGATC
GCCAACGGGTCGCAGAACTGACCGGTTTATCACCGGAGCAGTGCGCCGGCGGGGACGCGGACAGCGCGG
CAGTAACGCGCTGGTTCAACGACGGCATCGCTCCGAATGGGGCGCACGGTCGAGCCTTTCGCGCCACCG
ACAATCACGGTTTCGACGTCACTTTCAGCGTTCCGAAAAGTGTCAGTCTGCTGCGCGCATTCGGCAACG
ATGTGACGCAAAAAGCCGTCGAGGACGCGGCCAACCGTGCCGTCGAAGAGGCGATGGCTTACCTGTCCG

ATCACGTCGGATACACCCGCGTTCACAATCCGACCACGGGCGAAAAGACCTCCAAAAGTTGCCCGGTT
TGGTGTCGGTGGTTTACCAACATGAGACCAGCCGGGCAGGGATCC**ACAG**ATTAGCAGGTGTGCATGATC
TACGTGCGTCACATGCAGTAC

pKB1

CACTAGCTCAGATTCAGTAGACCGCTGTTGCACCTGCATTA**TACA**GTCCTTGGTGACGTTGAGTCTGTC
TCCAACGGAAGCATTGAAGACCACGATCCAGTCGGCCGCCAAGGCGCGCGGGATGCATGTTCTCGACTA
CCTGGGCGCCCTCGTGGCGAACGATCTAGGCCAACCCATTCCCGGCGTACCACCTTTGAAGGAGGTTCT
ACCGCTGCGTGATAGCGCATAACTGAAACGACGAACGCCCCCGGCGGGAACCGAGGGCGTTCGTGAAGT
TTGTGAGTGTGAGCCGGGAATCTCGCACTCAATGACCCCGAAGGGACAGTCTGTGTCGGACTGGTCTCA
GGGTAGCGCGTACCCTTTTGTTCGCGCTAGTTCTGATGGACGCGTGTCGGATCACGACTCGGTTACTGA
GTCGCGGTGGTCCGAAGTTTACCGGTTGCTGCCAGCTCCCGGTGGCGAAGCGTTCGCATTGGCGGGCAC
TGCGACGGCGAACAAGGCACGCCGTGAGCGCAGTGGCACTCCGATCGAACTGATCACAGAAGCAGAACG
GGCAGGTGGCGGTAAAGCCATCTGGGCGGTTGTGGCCCCGATTCGCAGCGATTTGCTTGTGATCGACCT
CGACCGCTGCGCTGAGCAGGTGTGGCCACGCCTCCGCGACGTGCTCGGCGACTACTCCGCGACTACCGC
TTACCTGGCTCGCTCCGGTAGCCCGGACTCGCTGCACGTGATCCTTAAGTGCGCCTCCCCTGACTCGGA
TCGCGGCATCCGCGAGTTCCTCGCAGATCTGCGCCGCCACCTCAACCTGTCCGCTCGTCGTATCGACGT
ACTGTCGCCTGGACACCTGTTACGCTTACCGGGGTCGGCTTCTCTTAAACCGGGCGGCCACCATTGTAC
ACCAGTTAATGACGACCTCACGCCAATCACCGCTGTTCGGACCGCAGAAAGGCTGCGCACCGCCTTAGG
TAGCGATGAACCCGGGCCAGTTCAGGCCGCATTCGCCCGTCACGACGATGGTTGGTGCGTACAGCTGGA
CGCGGAGGTATATCGTCCTGGGCTGGAAGTAGAAGTCCACCGTCGGGACGGTCGTGTCAAACGCGTTCG
GATCGGAGCCGTGGTAGCGCATGCTGGCGATCGGGTTCGGGCAGCGATTGCAGATGATCTGACACCACA
CAGCCCAACACCAGTGCCTACCATCGCGCTCTCTGCGGCCGCTCAAATGCACGAGCTGGCACCAGACAC
GGATGGTGCTGAGGAACTGCAGTGGCAGGCCCCACGCGCATGGCGTCGTCGTACGCCAATCAGCGCACA
ACAATGGCGGATCTTAAACGATACAGGCGGGGATGATCGTTCCGCCGCAGCTACCGCGGCAGCCTGGGT
TTTATGGGATGTAGGGCTCCGCTCTTTTGCCGCGGTGCGTTGGTTCTATGCACATTGTCCAGCGTTCGC
GAAGTTTCGGGATCGTGATGGGGATCGGCGTCGCGCCCGGTCGGCTTGTGCTGCACACTGGCAATCGAT
TACAGCGCGCGCTTCTACGCATCGTCCGCCTCTTGATCCTGCCGATCAAGCTTTAGTGGACGACGCTTT
AGCCGAGGTGGCAACGTGGGATGAGCCGGACCTTGTCGCAGCAGCGGTCGCTGTAATCACGCATCGGTT
TACCGATGGCCACGGACTCGTAGATCGTCCTATCGCAAAGCGCTGCTTAGCGACATGGATGTCTGTATG
CGATTCGCGCGCGTATGTCCACTTACAACGGCTTCAAGAACGTGGTTTGCTTGTGCGCACGCGGGATTG
GCAGGACGGACCAGCGAATGAAGCAGCGCTGTATGCGCTACGGGTGCCAGACACTTTTTACCGGGGAGA
TGGTGACCATGACGTCACAGACCACCAACGCCTCCACCCCCTGTGGGGGCAACTAGGCCACCATGCACA
CGCCCTGTGGTCTTCGCTGTCCACCCAAGAAAGTGCGGTCACCACCCCAGAAGTGGCAGCTAAGTCCCC
CCTGCCCTCCGGCGGTACCGGATGGGGCGTTCTGGGCCTGCTCAACACCCTCGGTTCGTTGGGGCTGGT
CACCCGGATCGGCACGGGGAAGGGAACCCGCTGGACAGTCACCCCCGCTCAGTCAGATCAAGGAGCTGT
TGACTCCGCGGCTACCTCTAGCGGTGCGTTCGAACGAGCGAAACAACTCCGGGCGCGCATCATCGGCGA
ACGCAAGGCCTATCACGCCGAAGGGCCAGGTGAGAGGGCTCGCATGCGGAGCGGGCTCAAGGTTCTTCG
GGACCGTCTAACCCACACCGATATTGCAGGACGGGAACAAGGCAGCCTGTTCGTTACCGTGGCAGCCGA
AACGTCTGGTGGCAGCCATTACCGGTGGGCAGGTGCGCGGCCTGGGCCGCCACCCGAAACGGCGACTCC
GCCCGATAGTTGAAAACCCAGTCCCCCTCTGGGCCCTCACCTCGTCGATGCTGCTTCGTGATCGTCTGC
ACCTCCATCGAGAGGCAGCGCTGACGAGCGTTGTAACGCTACAACTGCGAAAAACCTAAACATTAGAGC
TTGGGAACGTTAGGCGATGGAAACGTTAGACGCTGGGAACGTTGTAGGCGTGTAACGTTGTCACGTGGG
AACGTTGTGTAGTTAAATACTTCCAGTGCTACGTAGCTACAGCCCTACAAAGTTGCGGCAACGTAACGA
GATAACGTTTCAACATGGCAACTGCCAAACGTTGCACGGGTCAAAGGTTTCAGATTTACAACGTTGCAC
CGTTGTGCCGCTACAAATCTGAAACGGCAGAACGCTCCGGCGTTGAAGCGCCGGGATAAGTTTCGAGCA
TTGTGCAGTCACAACGTTTCAACGATGGAGGATCGCAATGGCAAAGATCGTCGTGCACCACTCCCGGAA
AGGCGGGGTGGGTAAGAGCACGGGAGCCTACGAACTCGCTCACCTACTCGATGCGGTCCTAGTTGACTT
CGAACATGACGGTGGGGGAGTGACCGGCAAGTGGGGGTATCGGCCGTTGGATCGGCTCCGCATCCCCAT
TCTCGATGCACTCCAAAGCGGCCGTACCCCGCGTCCGTTGAAGGGCCACGGCAAACCACGTCTGGTTCC
AGGCCACCCCGACCTCTACGACCAGATGCCGCAGCATGACGTCGTGGCCGACGCCCTTGAACGATGGGC
GGGGGAGTGGGACACCGAATGGGTCATCGTCGATACCCACCCCGGGATCTCCCCAGCTGCGGCTGGCGC

CCTCAGCGTCGCTCACGCCGTCATCGTGCCAACTCCACTTCGAAACCTGGACCTGACGGCCACGGAGGA
ACTGATCAACGACATGCCGGACTATCCGCTCATCATCTCGCCCACGATGATCCCGCCTGTTCCGCCTGA
GGCGGAACTCAGAAGGCTCGCGAGGGCCGTCGAAGGCACCCCGGTCCAGGTTGGCCCCCCGATTCCACG
AGCAGTTGCGGTGGAACGACGGACAAAGAGGATCGCCATCACTTCAGAGAACCCGCCGGCAAAGGCGCT
GCAGCCTGTCGCCGAGGCATACGAAAAGACCGCGAAGTTCGTCCAGGAGTACGTCCGATGAGTGGACTC
GATGACATGGCCCAGGCCCGACAGAAGCGCGCGAGGGCACGTTCAATGCCACCAC**ACAG**ATTAGCAGGT
GTGCATGATCTACGTGCGTCACATGCAGTAC

pMF1

CACTAGCTCAGATTCAGTAGACCGCTGTTGCACCTGCATTA**TACA**GGGAGCCGGATGGGTAGTTGGTCC
CGTCGGGTAGGGGTGCTCCATTGGGCCAGCGTATTGGGGACCACCGACAAAATCACGCGCCGAACGCCC
CCAACTCCCAAGTCAGACAACGCCTTCAGATGGAAAGTGTTCGCGCAGTGCCTGGCGTTGACTTCTTTG
TCTCGCCGGCAGTCATGCCGGGGTGAACTCGGCCGAAAACCGAGCACCGCTTCGGCGTGTCGGTACCAG
GTACAGGACTTTCTCTGGGGGACGATCGTCGGCATGGCCCAACCGCATAAGGGGCAGCGCGAGCAGATC
AAGGTCCGCGCTGCCGCAGTCGTGTACGCGCGTCTGCGCGAAATGGCCGCCCAACGCGGCACCAGCGTC
AGCCAGCTCTCGGCAGACCTCCTCGCCATTGCAGTCGGTCATCCCGAGGCCGTCCGCGAACTCGACAAG
GAGGTGCTGCCGCTGGCGATGTGAGAGCTCGCGCTGAGGCCCCGCCCTGCTCGACTGCGGGGGCCTTGG
CAGATGTGACATCTCAATAGCTGGTGTGCGGATTCGCCCGCTGACATGACGAAGGCCCCCGAAGGGGCC
TTGTCTGAAGGTATCGACCTGAGCGCCAACTCAGGTCCGGGTCCGCACCCGTTGCCCCTGAAAGGGACG
TGTCTTGGTCGACTCGTCTCACGGTAGCGCATGCCTGCCCGGCGTACCAGATAAGCCACGCAGCAAGTG
CGCGAATCGTGATCTTCTCACCGCCCGACCGAGTCATCGGGCGTGTTCTTCACCGCCACCGCTCGGCGC
GGCGGCCTATGCCGGACACAGCCCGCACGTCGAGCGCGCCCGGGTGGCACAGGGTCGGCGTGACTGGGT
GCTGCGCGCCGGTGCGAAACATCGCGCGCCGATGTGGACGAGTCGCGTGTCATGGTTGGACGGTCTGCG
GCGGTGGGCGCAGTCGCCGGCGCTGGGCGAGTTGTGTGCGGAGCAGCGGGTGTCGATGACCGCGGCCAC
GTTGTTGGCGATCGCGGCAGTAATGGCTGAGCACGCAGACCATGCTACCGGACGCCATGTGGCAGTAAC
GCGCGCGACCATCGCGGACACGGTCGGATGCGATGTGCGGACAGTTACAGCCGCCTGGCGCGTCCTTCG
CGTGTCCCGTTGGGCAGTAGAGGCTCAACGTGGTCATGGCTCGCCGACAACGCCATCCGTTGGTCGGCG
TCCGAGCGTATATCATCTGGTACCTCGTCGGGCGCCACAACCAGCGGGGGATCAGCCAGTACATGACTT
TCATCTGCCGCCACCTGGGGGAGATGGATTCTTGACACCGGTAGGATCTAAATCGCCGAGCGTACGTGC
ACGTGCACACGCAACATACTCGCGGGAACAGCAGCCAGATCCTCCTCGTCGCTGGCGCACGACCCCTCG
CCCGTTAGCAATCCAATTATTGGCTGCGCAGCTGGTCGCTCGTACCCACGGACTTGGCCGGGGCCATAT
CGGCGCAATTTGTGATACACTCACCGCAGCAGGTATCGAACCAGCAGTATGGTCGGCACGGGCAATCAC
AGATGCCCTGGATGCTGATATGCGGACGCGCGGCTGTAGCTGGCCTGATCACATCACCAACCCGGCTGC
CTTTCTGTCGAGCCGCTTGCGCCGTCTCTCGTGGACGACACCTCCGGCCCCTCCAAAGGGTGGGGGCTA
CGCAGCAACCCGGATTGACCAAACACCAGCTCGTGCAACGCTGACCGATGCGGCCCAAGCGCGGATCGC
AGCGGCGCGTGAACACATCCGCCAAGTGCTTACCAACCGCGCTCGACGCCCCCGCCCTGAGCGCACTCC
GAGCGCGTGTTCAGACGCACCGCAACGATCTATCGTCGAAGGCATGGCACCGTCCACTCCCAACGAGGC
GATTCCGCCGTCAACGCCGGTGCGCCGCTGGCGGCACGAGGAGTGCACCGTGCACGACAAACTGCCCGC
AGGGCGTATCTACTGAGTTACTGATCTACTGATAGAATGCGTTCATGACCGATGAAGGACCGCAGAGCG
GTAGCCAGATGAGCGCGGCTGAACACTGGCGGGCGTGGCTAGAACGCTACGGCGACGATTACGCCACGG
AGG**ACAG**ATTAGCAGGTGTGCATGATCTACGTGCGTCACATGCAGTAC

pMyong2

CACTAGCTCAGATTCAGTAGACCGCTGTTGCACCTGCATTA**TACA**AGGCGGGCAACACGACATCTCAAT
AGTTGGCACGGAGCTCCTAAACGACGACAGCCCCGCTAATGCGGGGCCGCAACGTCGATTTGGCTCTAC
TTCGTCTCTGGAGTTTCGTCTCTGGAGTTTCAGGTCCGAGCGGCTAACTCGGTCCTAGGCGTCTCCCGT
GCCGCGCCCCCGAAGGGACAGTCTTGCTGGACTGGTCTCACGGTAGCGCATACCCGAATCAGATTCGTG
GTGACGCGCAAAACCGTGATCTCACATTTACAGATTCCCGTGCATCCGTGTGCACGGATTCACGCCCAT
CTGGGCATTCACGTGGCCGTTTTGGTGCCTCCGTCTGGGCTCTCGTCGCTCCCGGCGGCGCGTCGTGGT
GACAACCGCGTTCGACGCGGCAGAGATTGCCGCCGCGGTCCCGGAATCTGGCTCACCGTGCAGCGGTAC
AGCGTCAGCTCGACTGCGCAGAATCCGCGCCAAGGAAGCCCGCAGGGCGTCGATCGAACGCTGTGGGGC
GGTGCCGGCCGCGGTGCCGATGTGGTCGTCGCGGGAGTTGTGGACTGCCGATCTGCGGGTGCTTTTGTC
GGGTCCGGAATTCAGCACGCGCCGGGTTATTTCGGCAGCGACGGTGCTCGCTGTCGCGGTGGCAATGGC

CGAATTCGCCGACCATGCCACGGGCCGCAATGTGGCGGTGACAAACGAGGTTCTGGCCGAGCGCGCGAG
GTGTTCTAAGCGTTCGGTGACCGCGGCGCGCGGGGTGTTGAAAGCGTTGGGTGTCGCGGTGGAAGCGGT
ACGTGGGCATGGCTCTGCGACCACACACACGGTCGGTAATCGACCGAGCATTTGGCACCTGGTAAGCCG
GCGTCAGCCAACAATCGATAATCCACCGACAGCTCCGCAAAACGGGCGCGGGGAACCTGCCGACACCGT
ACCGGACCGGGGGCAATCGGCACCAGTGGCTGTGGAGACTTGCGACCTACCACCATCCCGTAGGGATAG
GTGGGTAACTCCTGTTGAGAATTACTCACCAAGCACGCGCGAGCGCGCGAGCGCGGAAAATTCTTCCCC
AAAACAAACACAACCGGCCCGGTCCCGTCGTCGCTACCGGGCTACGCCGCGGCCTCTCGATGTACAGCG
CTTAGCAGCCGGTCTGGTGACGCCTGCCGTTGGACACGGCCCTGATAATGACGGACGTCGTACAGCGCT
CATTGCAGGTCTCGAGCAAGGACACATCGGGGCCATCTGCGACGCCATTACAACCGCAGGTATTGATGC
AACGGCGTGGACCCCGAAGACCTTAACCGCCGCCTTAAACGCGGACGCGCGTGTTACCGGGTGGTCCTG
GCCAGATCGGATTGAACGCCCAGGAGCTTTTCTTGCATCGCGTCTGCGGCGTCTTCCGGCGCGTCCTGA
TACCTCCGGGCCAGTGGATAACGGGCTTGATCAAGCTCGTCGGACACCAGTGGAACCATCGGCCGCCCG
GGTGGCTCCAGTGCAGACAGCAGCTGGCCGGGCCTATGCCCGTGCTTTGTTTGCGGAGCAACGTCGTCA
TCGGGTTACAGCTGCTAATGCCCAATCCGCAGCGGTCCCTGTACGTCAATCCGCACCAGAAACGGCTGT
CTGCGCTACCTGCGGATGCTCGGATGCACCACGGCGTCGGTTCTTGCCGACACGCCGTGCCCACATCTG
TGACGCCTGCTTCCAGGGCTGCGGTGGTGGTCAGGCACGTACCGGGCGTGTGGGGACAGTTGGTAGCAG
CTCGGCAGTACCACAGTGCCAATGATCGGGCAGGGCTTGCACGGGGATGCGGACCCATCCGCCGGCCGC
GGCCGGCGATGGGTCCAGTGTCGCGTTAGGGCCGTCGTCCAGCCGCGCCAGCTCGCGGTCGGATGCGAT
CAGCTTGGCACCGTAGATGTCGTCGTGGCGGCGGTCCCACGTGTTGGTGATGCGGTCGCGGTACCAGCC
TGTCCAGTCTTCGCCGGTGCCGCCGGCGGTGAGCCCGGCCCGGTACCACGCTTCGAGGCGATCACGGCC
CCGTGCGTGGTTGATGGTCTGGGCCGCAGCCCAGTCCGCGTCACGGAGGGCTCTGTCGGCTTCGTCCAT
GAACCCGCGCCGGGTCATGTCGATGATGGCTGGCGACGGGGCGTCGTAGGGGCCTGGCATGGGCGGAGG
CAGTCGGAACTGTTGCGGTTTGCTTGCGTTCAGTTCCTCGTCCTGCTGGTGGGTGCGGTCGTGCAGGGC
AAGGCTGTCGCGGTACCAGGCGTCGAGCGCGGTGCGCCAGTCCCCGCCAGTGGCGTAGGGCTCCCACGG
TTGCGGTGGCAGGGTGGTGAACTCGTGCAGGACCAGGCCGTCGATGTCGTCGCGCAGTTCTTCACCGGC
GATGGCGGCGATGCGGGCGCAGTAGCCGGGCAGATGGCTACGCCAGTCGGCGGGCAGTTCGCCGCTGCG
CGCCCACCGCAGCACATCGGCCCACAACCC**ACAG**ATTAGCAGGTGTGCATGATCTACGTGCGTCACATG
CAGTAC

pSOX

CACTAGCTCAGATTCAGTAGACCGCTGTTGCACCTGCATTA**TACA**CTTAGCTAATCAACTAGTGGTTGG
TACAGGTGGTTGGGGTACCACTCCAGGGAGGCGCAAGAATGAGCAGGCAAGCAGGCGATCTGACCCCAG
GAGTTGATGAAGTGCCACGGAGTGGAGTGCGCGGCTTTCGGGCCGACAGGCTCCGTGAGTTGCGTGTCA
AAGCGGGGCTGACGCCAGACGATTTGTCTGTACGACTAGGGTCCAGCCGTCAATCCGTTTCCCATTGGG
AAACGGGGCGTTCCACGCCAGCTCCCCCGGTGCTCAAGCAGATTGCACAGGAACTTGACGTGTCGATCA
GTGTGTTGGTCCCGATTCCGGACAACAGATTGCGTATGGGCGACCTTCGGGTGCGTGCCGGCCTAACTC
AAATCCAAGCTGCCGAACGCCTCGGAATCTCGCCCACCTCCCTTGCCGAGATCGAAAAGGGCATGAAAC
CGGTGAATGACCAGCGTGTCTCCGCGATTGCGGAGCTTTACAACGTCGAAACGGCGCAGGTTGTCGAAG
TTTGGGAACGTGGCCGTGAGACACGCGAGACACGCGCAAGATCCAAGTAATAAATCACGCAACAAAATC
ACGTGCCCGCAGCATTAATTCGGTGATCTAACATTCAGTGCGGCGTGTCACGCGAAGGAAACCGCCAAC
TTGGCGCAAGATGGCGGTGAACCTGGCGCACGTTCAGGCGCAGGATCGAAAGGAGAGGCATATGCCTAA
CCCAAGTAAAGGCAATCGCACTCGCGTGTCGACACGCCTTCCTGTGGACGTGGCCCAGGCGCTCGAAGA
GCTTCGGGCTAAGACGGGTGTCAGTTCGGAGAGCCAACTCGTAGCCGATGTCTTGGCACTGATAACTGG
TCACTCTGAACTCGTGGTCGAGCTTGACCAGGGAATCCTGTTCGATCTTGGACTCGATCAGAAACTGCA
CCGTGAGGAGGTTCTGCCGCTGACTGCGTAGCGAAACCCCTTCTCCGGCAGGAGAATACGTGACAATTT
CACACGCTTTGCAGTAGCACGAACCCCCTAGAAATCGAAAATCCCCCGGCGGGCAGGCCGAGGGTTTCG
AAGTGGGTGAGAGGTTGAGCTGGTAACTCGCGCTCTCGGTGGTACTGCGCTTCCCCGAAGGACAGTCT
TGGAGGACCAAGGGTAGGGTACATCGCGCCCTGTTGCAGATTCAATAACTGATTTTTCGCGTGTCTTGG
TGATGGAACACGCGCCCACATCCGTACAACGCCCGTACTGCGGGAGCAGCCCCACGCGTCGCCGGTACG
GCACAGCCCGCGCACCCCGCACAGTCCGCGGGCACGTCCGCGGGCATCAGTGCCAATCGCTCACGCCGA
TGTTCATCAAGGCACTTGGCGGGCCGAAAGAAGCGTGCCGGCGGTACTGCCAGATGCGGCTCGACCTCG
GCGACGGTGCATACAACGGCATCCCCATCTGGCAGGGCGCTCAGCACTGGGTGGAAATCGCTGTCCGCG

AGGCATATACGGCCGAATACAAGAACATTCGCCCCGCCCTGGTTGAAACGACAGGCGGCGGAATAAGCC
TCAAGACTCTCCTAGCTGTCGCCACAGTCATGGCGTCAGTCGCTGAGTTCGACACTGGCCGTGAATCAC
GTTTGTCTCTCGATAAGACCATCGAACGTACGGGCAAGGGTGAGAGAACCGTGCAGCGTGCCCGTCAGG
CCCTGAAACTGCTTCGAGTGGCCACCGAGGTCTTCCGTGGACGGCTTCGTAGGAAGAAGGGCGAACGAC
AGGGTTCGTACCGGGTGGGCGACAAGGGCCGGGGATGGGCGTCAGTCTGGGCACTCCACCTACGTAAGC
CTGTGGATAAAACCCGTGTCTACCTGGACGGATCCATAAAGATGGCACCCCATCCCCGTAGGGGTCATC
TTTTGTCTCTTCCCTCTCGTAGAGAGGTTATTAATACAAAACGATCTGTGAATAAACGAGCCGCTCCGC
GGCGCAAGGAATCCAAGGCCGAGGTCGAGAAGGTTGAAGCCATCCGAAAAGGCCTGTTGCTTGCATCCA
AGTGGCTCAGCAATCCGCGAACGCCGGTCTGGGCGCGTCGGCACACCCCCCGAGGGTGGGCATCGGCAC
TGACCGAACCAGCCGCGCACGGTTGGACCGCAGCGGATCTGAACGACACGATCGACGACTGGGCGAACG
CCCAGAACATGGTTCCGACCCCGAAGCACCCCATCGCATTCATTCGTTGGCTCATGAAGCAGCAGGATC
TGGCCTTCGCACCGCATGTTCTCGCTCAGATTGCCGCTGACCAGGAGAAAGCTGAACGTGAACGCCAGT
CCGCCGCGTTAGAGATGGAGCGAGAACGCTACGCATCGGCCGCGCCCGAGGACTCCCCTGGCCGCCAAG
CAGCGCGTTTGGTTGCTCGTCGAGCAGCCGACACGGCTCGCTGCCGCAAGGTGGATACCTCAGCTCGTG
AGAACGCCGCGCAACCTGTGTGGATCACGCATTTACGCGACCTCGGACCGCAGTGAGCAGCCGCGATCA
GCTTTCGGTGAGCGGAGAAATCTGTACGATCGCCAAACGCTCGGTCAAAGGTCCTGGGTACGGGCTGAT
CCGCAGGGTCGAAGGACGCCAATCATCATTTTCAGTCCGAAGACGGGCCACGACGTGGACCGGATCGGT
CGAACCTAGATTGAACTTGTCCTGGGCCGAAGACAACAGATGTCGATCGTCAGCGTGGATGACCTGGTT
CGCGCCAGGAATGTCGCGCCACTTCACCCACGGCGCTGGTTCAGGCAACCACAACGCCAACACAGCGGT
GGCTTACATGGCGATAGCTAGACTGGGCGGTTTTATGGACAGCAAGCGAACCGGAATTGCCAGCTGGGG
CGCCCTCTGGTAAGGTTGGGAAGCCCTGCAAAGTAAACTGGATGGCTTTCTTGCCGCCAAGGATCTGAT
GGCGCAGGGGATCAAGATCTGATCAAGAG**ACAG**ATTAGCAGGTGTGCATGATCTACGTGCGTCACATGC
AGTAC

pVT2

CACTAGCTCAGATTCAGTAGACCGCTGTTGCACCTGCATTA**TACA**CCCAATAACCCTTCGGTTAACGGC
CGCAGCGGAATTAACGGGGCGGCGTACATACATATAGGCATCAATACGCGCGTACATATAGAAATATAT
AACCGCATTACGCCAGTTCAGGGTATGCGCCCACAATCGCCGCCGACACCAGCGCTTGCACACTGGTGT
CCTCGTCAGCCGCCAATCGCTTAAGCGTTCGGTGCATGCCGCGGGGAAGATCGAGCGTGGTCCGCGCCC
CGCCGCTTGCGCGTCGGTGTGCCATCGAGGCCGCCGCTAATCCTGGTGCGTTCTGCAGCCCAGCATTGA
TCGCCGCGCGAATCAGTTCCGTCATCGTGGTGTCTGCGTCCAGAGCGCGGCGCTTGAGTCCGGTCCGCA
GCGACGACGGCAGGTGCAGTGTGGTGCGCATCGCTTCATGAACTGCGGCTGCCGCACCGCTGGCGGGTG
CGTCATCGGCACTTTGAAATGTGCTCGCTGCACGCGCCGCTGCGGCTGCACCGGTCGGACGGCGAATAA
CCGACGCACGCACCGCATCCGCTCTCGTTGACTTCGCCGTCATACCGGTACCTCCTCAAGTCCTACCGA
GGCCCACAGCTCGTCGAAAACATCGCCGTAGTCACCCAGGTACCGCGGAATGCCGCGGCCGAATGCCCG
TTCGACCGCGGTCAGATCCCGAACGACGTTGCGCAGTACTGGCGCGTTCGCTTCAACCAGACCACGCGG
CATTTCCACCGCCTCCACCTTGCGCATGTCGACCAGAGTCAAAAGGACCGTCGTCGACCGGTGCGCAGT
GATATCCAGCGTCGGCCACACCCGGTCAATATCCGCCGCCCGAGGGCCGGTCGGAATGATCACTAGGTC
GGCCGCATCAATCGCCGCGTCGATCGCAGCCGCCGTTCCCGGCGGCGTATCCACCAGAACCAGCTCACC
TGGCTCGCTGGACAGCTCTCGCAATCCCGCCGCGGTTACCGGCGACACGTCAAACGGCAGCGGTGTACC
TATGTGCGCCGCACGATCCGCCCACGAGCTCGCCGATCCTTGTGGATCAGCGTCCACCACCCGCGCCGG
CACACCCCGACGCACGGCCGCTGCTGCCAAAAACATGCAGGTCGTCGTCTTCGCGACGCCGCCCTTCGT
GTGCACAAGTGACCAAATCACGATGCAAAACCATACATCCATATGTGCATCTACACGCGCACGTGGACG
CGGCGCGGCGCGCGTGACATCTAGCCGTTTTTCTGTCAAAATGGCGGTATGGCGCGAAAGAAGACCACC
GTCTACATCGACGAGGCGCTGCTGCGCGCCGCGAAGGTCGCCGCCGCACGCTCAGGCAAGCGCGAATAT
GAAGTCTTCGAGGACGCGCTGCGGCGGCACCTGGGATTCGCCGAGACCCTCGAGCGGATCTGGGCCGGC
ATCGGCCCTGAAGGGGCTCCCAGCGAGGAAGAAGCTGCGCAGCTCGCCGCCGAAGAGTTGGCCGCGGTA
CGGGCGCAGCGCACGCCCCGCCAGGCGGGCTGAGTCCTGGACCAACCACTGCGGGTCGTCATCGACCCC
AACGTGTGGATCTCGGCTGTAATCAACCCGTACGGCACCCCCGCCCGGGTCGCGCAGGCCGTAGCCGAC
GGCGCGATCACGGCCGTGGCGACCCAGCATCTGCTCGACGAACTCGCCGCCGTGCTGATCCGGCCGAAA
TTCCGGCGCTGGATCAGCGTCGCTGACGCCATCGCCTTCGTCGAGAGCCTCGGCGGCCAGGCCGATCTG
CACGACGATCCCGGACCACCCGAGACGAAAGTGCGCGACCCAAACGACGACTACCTGGTCGCACTCGCT

GACGCGGCCGACGCCGTGATCGTCACCGGCGATAACGACCTCCTGACCGCCGGACTTGAACCACCAGCG
ATTACACCCGCTCAGCTTCTCGCGCGTCTCTAAACTTCATCAATATGTGCGTCCATAAGTACGTACATA
TGGCTACATCTGCTTGCATTACCGCACGTCAGGTGCTCACCGGCCCTTCCTCGGCCGCCATTGTGGTGA
ATGTGGGGATGGAGGCTGTCAAGGCCATGCCGCTGCGGCGGTCGACCAAGAGTCGAGCCTTGACAGCCA
CCATCCCCACGCACAGGAGCGGCCAGTATGAGGAAGGGCCAACCAAGAAAGGGGCCGTCACATGTCGGA
GTGTCGTCATACTGTGGGTCCCGTGAATGCTGGCGAGCACTGGCGCGCATGGCTTGATCGGTACGGCGA
TGACTATGCGACCGACGACGAACGCCGTGCGGCCTACCGCGATTTCAAGGCCAATCTCGCCCGGTTGAC
CGAAATGTTCTCAGCTGGCGACGACGATCACGGCAATCCGTAACGGCGTTCAGCAATCCCGTACAGTGC
CGGCCCCTGCCCGCCGGCAACCGTCTACGTCCACGCCGCAATTCGCCGCCGAGCTTCCGCGCTCGATCA
CGAAAACATCACAGCAGCGCGGCGTGTCGGAGCTGAGTATCTGATACCGGCGGCGACGATAAGGGCTAT
GGCTCAACCACACAAAGGCCCGCGCCGCCTAGTACAGACGCGGATTCCCGAACAGGTTTATGCCGAACT
GGTCAAACGAGCACGTGAAGCCGGAACGTCAACTTCCCAGTTCATCGCCGACTCAATGGCCTTATCCGT
CGGTCGGGACGATCTCGTCTGGGAGCTGGGCCGGGAACGGGAGGCGTTGCCACTGGCGATATGACCACA
AGGGCCCCGGAACGCCGCCCCGTGCATAGCGGTTGTCGTAGCCGGGCAGACGTGACAGCTCAATAAACG
GCGAAACCCCCGGCTGCCGGGCCGAGGGTTTCCGCTTCCCCGAAGGGAACTGCTTTGCTAGGCAGCTCT
CACGGTAGCGCATACCTGAACCAGGTTCCAGACGGCGCGCGCCGTGACATTTCTGAAATTGCTTCGCCG
CCGTATCGCGGGTGTACCTCGTTGCGGCCGCGCCGTGGTGTGGTGTGTACGCGGCCGTGGGCGATGCGG
TTGGCGGCTGCCGAGCATGAGCACCGCCGGCAACAAGCAGCCACGGCGGTCGCGGCGATTGTGCTCGAG
CTCGGCGAGGCCCCGTATGCGGGGGTGCCGTGTTGGACCGGGCGGCTCGAGCGCTGGGTTAGGTGGACG
GTTCCGGTGGCCTACGACTGCCGCTACGACACCGAGATACGCCCACTGATGCCAGGGAATCCGATTTCC
CGGCAAGCCTTGTTGACAGTTGCCGAAGCACGGGCGCGCTACGCCGACCACGCGACAGGTCGCAACTGC
CGTCCATCTAATGAGCGGCTTGCCGCGGATACGGGACTTAGCGTGCGGACCATTCAACGGGCGGACACC
GTGCTCCGTCTCCTCGGAGTGGCTACCGAAGTCCTGCGTGGGCGGCAGCGTACACGTGTCGAGCGCCTG
GCGTCTTGGCGTGTAGGAGATCGTGGGCGCGGGTGGGCCAGCGTATGGGCTTTGCACGATTATCCTCAG
CTCACCCGCCTTGTGTACCGGTTATCCCCGCATCCACGTAGCGGGCCAGTCCGGGATCAACCGAGCGGA
GAAAAAGTAATTACAACGGATCCTGGCGGGCCGGCAGGCCGTCGGCAGTCTGGCGCGACCCGCCGTCGG
ACCCCGGACGCTGGTGGTTCTGCGTTAGCTCGGACATGGCGGGCTGATACACACGCGCCGCCTTGGGCG
CGGCGGCATACCGCTGGCGCATGGGCCGCTCTTCTTGCGGGACCAGCCGCCTACGGCTGGTCGCCACGC
GATCTGAACGCCTTGATCACCGATTGGGCAGCTGTGACCGGACGCCGTATTCCAGATCATCCTTATAAG
CCAATCGGTCTCCTGGGGGCAATTTTGGCCTGGCATGGACGGGAACGTCTCGCCGAACGCCCGGCTGCC
TTAGACGAGGCTCGTGAGGCGGCCGAACTCGCTACACATCATGCCCACCTTGCAGCTCAGCGCGCTGCG
CACCGCGAACACGAACGCGCTCGGGCTGCAGGGCGGGCAGCGTTGGGTGGACCTGGCCATGCGGCAGCA
CGTGCTGTGGCGGCGGAAGCGGCCCGTCGTGGAGCGCATAAGCGCACCCACGTTGTGGCCGAGGCTGCT
GCGCAACGTGCAGCCGCTGTCGCGGCCGCACGCACACATCGTCGTGGACCGGGCCCACTTGGATGATGG
TTGTCGAGCCCGTCGCCGCCGAGGCCCGGCCAGTCGCGCTGATCGCAGCACATCACGTCGAGGCGCGGC
AGCCGCGGCAGCGGTGTCCCGTACACCAGCCAGTCGAGCACCCGTTCCCCACCGGGCTGGGCGCGGCGC
ACCAGCAGCTCGGTCACCTTCGCCTCGACGCTCATCGGTTCCCTCCGCTTCGTGCGGGCTGGCGTGATG
TTCCCATCATCAACCCGCGAAGCGGTGCCCAACCTGGGTCACGGTGACGTATCTGGGTGCGCTGGATG**A**
**CAG**ATTAGCAGGTGTGCATGATCTACGTGCGTCACATGCAGTAC

pDA71

CACTAGCTCAGATTCAGTAGACCGCTGTTGCACCTGCATTA**TACA**AGGCCTGACGGGAGTAGGAATCGA
CGGGGCCGTCGTGCCACGGCTTGCGGCCCTTGCGCGGTCGGCGGTCGGGGTGCCGCTCGTCGAGCTGGG
CGGCGAGCTCGGCCTTGAGGCGGGCGCGATGCTCGGCCGAGGACCCGCTCGGGCCGCGAGTGAGGCCGC
GAGGGTCGGCGGCGAGCTCGGCGACCGTGGGCTTGGACGGCCGGCGGGATCGAATGATCTGCGTGATAT
GCCCGGGCATGATGCGCTCGGTGCTGGCCGAGTAGTGCACCGCGATGGCCATGAGCGCGGCCTGCGGAG
TCCAGCCGGCGAGGCGGGCCGCGAGCAGCCATGACTCGCGCATAGTCTCGTCGATGTTGCGGTTGTCGA
ACGACTGCGCGGACTTGAGCAGCTCGTCGACGAGCTCGGTGGGGTTGTTCACGGGGTGTTCCTTTCGCG
GGGTGGGTGCCTGGCGGGGCGGTTACTGGCCGATGGCGACGAGGGGGCGGCCCTCGGCCTCGGCGGCGT
CCATGGCGTCGAGCAGCGAGCCGATGAGGCTGCGCTGGGGCTTGGGGGCGGCGTCGAGCAGCGAACCGT
CGTAGGCGAACCCGTGCTCGTCGACGACGGCGGCGGCCTGGGCTGCCTTGGCCTCGGCCGCGGCGCGCT
GCTCGGGCGACGGGGTGATGCCGACGAGGGCGAGCAGCGACAGGTCGGGGCAGGCCTTGCGTGACTCGG

TCTGCTGGATCGACTTGATTTCGTGCGGCGTCAACGGCAGGCCGTTGGGCCAGCAGGGGCGAGGACCCT
TGCCGTTGCGGGTGTCGCGCTCGGCGAACTCGACGGCCTGTCGAACCCAGTTCCGGTAACCGGCGTCCC
AGTCGGCGAGCTTGCGGGCATTGGCCCGGGCGTGGCCCTCGAACTTGGTGAACTCGCGGCGCACGTCGA
TGCGGTCGACGAGGCCGGCCTTGGTGAGCTCGGCGACGAGGCCGTCGACGGTGGCCTTGGTGGGCTGCC
ACCCGTCGGGCATCTCGTGCTTGGTGCGGCGGGTGCGCTTGGGCTTGGGCTTGGGCTCGTCGCCGGCCT
CGGCCCCGCCCTCGATGAGGGTCAGCTCGGCCCGGGGAGCGTCGCCCCCCCCGTGGGGGGTAGGGGGGT
TCTTTATCTCTTTATCTTTCTCTATCTCTAACTCTGGTTGGGTTTTGCTTGCGTTTTGCTTAAGCACTT
GCTTAGCACTTGCTTGAGCATTTGCTACCCCCTGCGAGGCGTTCGCGGCCCTCGCCTGCCCACCCTTTT
TGCCCGCTGAGCTGCGCGCTTCCTTGAGTGCCTCGACCTCGGCCTTGCTCTGCTGGTGGTCAAAATAGG
CCGGAAAAAAAACGCAATCCGTCGACAGGATTTCGCTGGACTTCGTCGAAAAAAATTCGGCGAGCGTCG
AATAATCTGCGCCGCCGCCCTTGGGGATGACGTGGCAGATGCCCGTTGCCTCGACGGCCTTGCGGGAGC
CTGCGCCCGCCATCTTGCGCCACTGAGCCATGGTGATGATGCCGTCGGTGAGGTTGCGCCGGCAGTACA
TGAGCGACTTGGTGATGAGCCATTTCTGGGCGTCGGTGAGGGCGGCGAACTTGGGGTGTTCGTCCATGT
CCACCGCGAGGGTGATGTAAATGCGGTTGTCTTTGGGCATGTCGTGGTCCTTTCGGTTAGGCCGCGTAG
CCCTGGCGGGGCGCGGGGGCTACTTCGTCGTCGGGGCGTAGCGCGGGTGGGATGTTGCGGTCCTCGCGT
ATCCGGATGACGGTGTAGAGCGTCCAGCCGGTATGCGAGGCGATCTGTTTGTCGGTCCACCCCCGGCCG
ACGAGGGCACGCATGAGCGCACCCCGGTCGGCCGGCGACAGGACCTCGGCGGGCAGCTCGCCCGCCAGC
GCGGCCCGGTAGGCCGGCGGATTGTCGTGTGGTGCCGACGTAGCGTTCGCCGCGCAGGGCCTCGGGGGC
CTCGACCGTGGGGGCGCTCACCCGAACACCACGTGAGCGGTGGCGGACAGTGCCCACAGGCCCGCAACC
GAGGTCACGAGTGCGGTCGCGGTTGCGATTCCCATAGCCACCGATGACAGGGTGTCGCGGAAAGGCGAG
TCGTCGACATCATCGAACAAGGGGCCGGTTTGGCCGCGGTCGATGATAGAGTTGTGGTGCATCTTGAGG
TTCCTTTCGGGTGCTGGGCTCGGCTACATGCATTCCCCTGGCGGGGGGTTGTGGCCGAGCCCAAGGGTT
TTTTCGGGGGTCTTAGGCGGCGACCCGTTCGTCGTCCTCGGCTTGAGTGCTGCCCGTGGGCTGCGCTTG
TAGCCAGTCGTAGAGCTCGTGAATGTCGATGCGCGGCGTGCGTTCGCCGATATACCGGACTGGCAGCGT
GCCGCCCTTGTAGAGCTTTCGGACGTGGTTCGGGGTTAGGCCGAGCAGCTCGGCCGCCTCGGTAACAGT
GACGAGTCGCCACGGAATGTTCGCCTTTGCGCCGCCGAGGTCCGCCGGCGCGGCCGTTGTGTTCTCTGC
CATGTTCTCGCTCTCGTGTCTCGGGGGGGTGCTCGGTGGTGCTCCGCAACTATAGGCGCAACCGGGCCA
CAACCGCAACCTCGTTGCGGGGCTTGCTCGTACTCCTGGGCAATCCTGGGAAATCTTGAACGATGAGAA
CGTAGCTCGAGCCCGGCCGTCCGTCCGGCAACGACACGCCGGGGCGTGTCAGGAAACTTGCGCCGGATC
**ACAG**ATTAGCAGGTGTGCATGATCTACGTGCGTCACATGCAGTAC

**Table A3:** DNA sequences of all Actinomycete resistance markers used in this work. The protein coding sequence is highlighted in yellow. The AarI restriction enzyme recognition sequences are highlighted in green. The cut site is underlined and in bold.

| Origin |
| --- |
| Apramycin resistance marker |
| CACTAGCTCAGATTCAGTAGACCGCTGTTGCACCTGCATTA**ACAG**CCGGAATTCTGATCATCATTCGCC<br>GTGGCAGCGTGGGTCGAATGAGAATACGAATGGATTGGTCAGGGATTTTTTCCCGAAGGGCACTAATTT<br>TGCTAAAGTAAGTGACGAAGAAGTTCAGCGGGCACGGATCCTAATTAATTAAGAAGGAGATATACATAT<br>GCAATACGAATGGCGAAAAGCCGAGCTCATCGGTCAGCTTCTCAACCTTGGGGTTACCCCCGGCGGTGT<br>GCTGCTGGTCCACAGCTCCTTCCGTAGCGTCCGGCCCCTCGAAGATGGGCCACTTGGACTGATCGAGGC<br>CCTGCGTGCTGCGCTGGGTCCGGGAGGGACGCTCGTCATGCCCTCGTGGTCAGGTCTGGACGACGAGCC<br>GTTCGATCCTGCCACGTCGCCCGTTACACCGGACCTTGGAGTTGTCTCTGACACATTCTGGCGCCTGCC<br>AAATGTAAAGCGCAGCGCCCATCCATTTGCCTTTGCGGCAGCGGGGCCACAGGCAGAGCAGATCATCTC<br>TGATCCATTGCCCCTGCCACCTCACTCGCCTGCAAGCCCGGTCGCCCGTGTCCATGAACTCGATGGGCA<br>GGTACTTCTCCTCGGCGTGGGACACGATGCCAACACGACGCTGCATCTTGCCGAGTTGATGGCAAAGGT<br>TCCCTATGGGGTGCCGAGACACTGCACCATTCTTCAGGATGGCAAGTTGGTACGCGTCGATTATCTCGA<br>GAATGACCACTGCTGTGAGCGCTTTGCCTTGGCGGACAGGTGGCTCAAGGAGAAGAGCCTTCAGAAGGA<br>AGGTCCAGTCGGTCATGCCTTTGCTCGGTTGATTCGCTCCCGCGACATTGTGGCGACAGCCCTGGGTCA<br>ACTGGGCCGAGATCCGTTGATCTTCCTGCATCCGCCAGAGGCGGGATGCGAAGAATGCGATGCCGCTCG<br>CCAGTCGATTGGCTGAGCGGGACTCTGGGGTTCGAAATGACCGACCAAGCGACGCCCAACCTGCCATCA<br>CGAGATTTCGATTCCACCGCCGCCTTCTATGAAAGGTTGGGCTTCGGAATCGTTTTCCGGGACGCCGGC<br>TGGATGATCCTCCAGCGCGGGGATCTCATGCTGGAGTT**GAGT**ATTAGCAGGTGTGCATGATCTACGTGC<br>GTCACATGCAGTAC |
| Spectinomycin resistance marker |
| CACTAGCTCAGATTCAGTAGACCGCTGTTGCACCTGCATTA**ACAG**CCGGAATTCTGATCATCATTCGCC<br>GTGGCAGCGTGGGTCGAATGAGAATACGAATGGATTGGTCAGGGATTTTTTCCCGAAGGGCACTAATTT<br>TGCTAAAGTAAGTGACGAAGAAGTTCAGCGGGCACGGATCCTAATTAATTAAGAAGGAGATATACATAT<br>GAGGGAAGCGGTGATCGCCGAAGTATCGACTCAACTATCAGAGGTAGTTGGCGTCATCGAGCGCCATCT<br>CGAACCGACGTTGCTGGCCGTACATTTGTACGGCTCCGCAGTGGATGGCGGCCTGAAGCCACACAGTGA<br>TATTGATTTGCTGGTTACGGTGACCGTAAGGCTTGATGAAACAACGCGGCGAGCTTTGATCAACGACCT<br>TTTGGAAACTTCGGCTTCCCCTGGAGAGAGCGAGATTCTCCGCGCTGTAGAAGTCACCATTGTTGTGCA<br>CGACGACATCATTCCGTGGCGTTATCCAGCTAAGCGCGAACTGCAATTTGGAGAATGGCAGCGCAATGA<br>CATTCTTGCAGGTATCTTCGAGCCAGCCACGATCGACATTGATCTGGCTATCTTGCTGACAAAAGCAAG<br>AGAACATAGCGTTGCCTTGGTAGGTCCAGCGGCGGAGGAACTCTTTGATCCGGTTCCTGAACAGGATCT<br>ATTTGAGGCGCTAAATGAAACCTTAACGCTATGGAACTCGCCGCCCGACTGGGCTGGCGATGAGCGAAA<br>TGTAGTGCTTACGTTGTCCCGCATTTGGTACAGCGCAGTAACCGGCAAAATCGCGCCGAAGGATGTCGC<br>TGCCGACTGGGCAATGGAGCGCCTGCCGGCCCAGTATCAGCCCGTCATACTTGAAGCTAGACAGGCTTA<br>TCTTGGACAAGAAGAAGATCGCTTGGCCTCGCGCGCAGATCAGTTGGAAGAATTTGTCCACTACGTGAA<br>AGGCGAGATCACCAAGGTAGTCGGCAAATGAGCGGGACTCTGGGGTTCGAAATGACCGACCAAGCGACG<br>CCCAACCTGCCATCACGAGATTTCGATTCCACCGCCGCCTTCTATGAAAGGTTGGGCTTCGGAATCGTT<br>TTCCGGGACGCCGGCTGGATGATCCTCCAGCGCGGGGATCTCATGCTGGAGTT**GAGT**ATTAGCAGGTGT<br>GCATGATCTACGTGCGTCACATGCAGTAC |
| Kanamycin resistance marker |
| CACTAGCTCAGATTCAGTAGACCGCTGTTGCACCTGCATTA**ACAG**GGCTTACATGGCGATAGCTAGACT<br>GGGCGGTTTTATGGACAGCAAGCGAACCGGAATTGCCAGCTGGGGCGCCCTCTGGTAAGGTTGGGAAGC<br>CCTGCAAAGTAAACTGGATGGCTTTCTTGCCGCCAAGGATCTGATGGCGCAGGGGATCAAGATCTGATC<br>AAGAGACAGGATGAGGATCGTTTCGCATGATTGAACAAGATGGATTGCACGCAGGTTCTCCGGCCGCTT<br>GGGTGGAGAGGCTATTCGGCTATGACTGGGCACAACAGACAATCGGCTGCTCTGATGCCGCCGTGTTCC<br>GGCTGTCAGCGCAGGGGCGCCCGGTTCTTTTTGTCAAGACCGACCTGTCCGGTGCCCTGAATGAACTCC<br>AAGACGAGGCAGCGCGGCTATCGTGGCTGGCCACGACGGGCGTTCCTTGCGCAGCTGTGCTCGACGTTG |

TCACTGAAGCGGGAAGGGACTGGCTGCTATTGGGCGAAGTGCCGGGGCAGGATCTCCTGTCATCTCACC
TTGCTCCTGCCGAGAAAGTATCCATCATGGCTGATGCAATGCGGCGGCTGCATACGCTTGATCCGGCTA
CCTGCCCATTCGACCACCAAGCGAAACATCGCATCGAGCGAGCACGTACTCGGATGGAAGCCGGTCTTG
TCGATCAGGATGATCTGGACGAAGAGCATCTGGGGCTCGCGCCAGCCGAACTGTTCGCCAGGCTCAAGG
CGCGGATGCCCGACGGCGAGGATCTCGTCGTGACCCATGGCGATGCCTGCTTGCCGAATATCATGGTGG
AAAATGGCCGCTTTTCTGGATTCATCGACTGTGGCCGGCTGGGTGTGGCGGACCGCTATCAGGACATAG
CGTTGGCTACCCGTGATATTGCTGAAGAGCTTGGCGGCGAATGGGCTGACCGCTTCCTCGTGCTTTACG
GTATCGCCGCTCCCGATTCGCAGCGCATCGCCTTCTATCGCCTTCTTGACGAGTTCTTCTGAGCGGGAC
TCTGGGGTTCGAAATGACCGACCAAGCGACGCCCAACCTGCCATCACGAGATTTCGATTCCACCGCCGC
CTTCTATGAAAGGTTGGGCTTCGGAATCGTTTTCCGGGACGCCGGCTGGATGATCCTCCAGCGCGGGGA
TCTCATGCTGGAGTT**GAGT**ATTAGCAGGTGTGCATGATCTACGTGCGTCACATGCAGTAC

Zeocin resistance marker

CACTAGCTCAGATTCAGTAGACCGCTGTTGCACCTGCATTA**ACAG**GGTCGAATGAGAATACGAATGGAT
TGGTCAGGGATTTTTTCCCGAAGGGCACTAATTTTGCTAAAGTAAGTGACGAAGAAGTTCAGCGGGCAC
AGGATCAAGAGACAGGATGAGGATCGTTTCGCATGTCCAAACTGACGTCGGCCGTCCCCGTCCTCACCG
CCCGCGACGTGGCGGGGGCGGTGGAGTTCTGGACCGACCGGCTGGGGTTCAGCCGCGACTTCGTGGAAG
ACGACTTCGCCGGGGTCGTCCGGGACGACGTGACGCTGTTCATCTCCGCCGTCCAGGACCAGGTGGTGC
CCGACAACACGCTCGCCTGGGTGTGGGTCCGGGGGCTCGACGAGCTGTACGCGGAGTGGTCGGAGGTGG
TGTCCACGAACTTCCGTGACGCTAGCGGTCCCGCGATGACCGAAATCGGCGAGCAGCCCTGGGGTCGGG
AGTTCGCTCTGCGGGACCCCGCTGGGAACTGCGTCCACTTCGTCGCCGAGGAGCAGGACTAACACGTCC
GACGGCGGCCCACGGGTCCCAGGCCTCGGAGATCCGTCCCCCTTTTCCTTTGTCGATATCCTGTCAGAC
CAAGTTTACTCATATATACTTTAGATTGATT**GAGT**ATTAGCAGGTGTGCATGATCTACGTGCGTCACAT
GCAGTAC

Nourseothricin resistance marker

CACTAGCTCAGATTCAGTAGACCGCTGTTGCACCTGCATTA**ACAG**GGTCGAATGAGAATACGAATGGAT
TGGTCAGGGATTTTTTCCCGAAGGGCACTAATTTTGCTAAAGTAAGTGACGAAGAAGTTCAGCGGGCAC
AGGATCAAGAGACAGGATGAGGATCGTTTCGCATGACGACCCTCGATGACACGGCCTACCGGTACCGGA
CCTCGGTGCCGGGCGATGCCGAAGCGATCGAAGCGCTCGATGGGTCGTTCACGACCGACACGGTGTTCC
GGGTGACGGCGACGGGGGACGGCTTCACGCTCCGGGAAGTCCCCGTGGACCCGCCCCTCACGAAAGTCT
TCCCGGACGACGAAAGCGATGATGAGTCCGATGCGGGGGAAGACGGCGACCCGGATAGCCGGACGTTCG
TGGCGTATGGCGATGACGGGGACCTCGCCGGGTTCGTCGTCGTCAGCTATTCCGGCTGGAACCGGCGCC
TGACGGTGGAAGATATCGAGGTCGCGCCGGAACATCGGGGCCACGGGGTCGGCCGGGCGCTGATGGGCC
TCGCGACCGAATTCGCGCGGGAGCGGGGCGCCGGGCATCTCTGGCTGGAGGTCACCAACGTCAATGCGC
CCGCGATCCACGCGTATCGCCGGATGGGCTTCACCCTGTGCGGGCTGGACACGGCGCTGTATGACGGCA
CCGCGAGCGATGGGGAGCAGGCGCTCTATATGAGCATGCCCTGCCCGTGACACGTCCGACGGCGGCCCA
CGGGTCCCAGGCCTCGGAGATCCGTCCCCCTTTTCCTTTGTCGATATCCTGTCAGACCAAGTTTACTCA
TATATACTTTAGATTGATT**GAGT**ATTAGCAGGTGTGCATGATCTACGTGCGTCACATGCAGTAC

**Table A4:** List of all Actinomycete strains used in this work.

| Species | Strain designation | Repository code | Transformants |
|---|---|---|---|
| *Dietzia dagingensis* | 263 | DSM 44748 | No |
| *Dietzia maris* | 32D | DSM 44747 | No |
| *Dietzia natronolimnaea* | 15LN1 | DSM 44860 | No |
| *Dietzia psychralcaliphila* | ILA-1 | DSM 44820 | No |
| *Gordonia alkanivorans* | 2102-488, HKI 0136, JCM 10677, IFO 16433, NBRC 16433 | DSM 44369 | No |
| *Gordonia paraffinivorans* | AS4.1730, HD321 | DSM 44604 | Yes |
| *Gordonia polyisoprenivorans* | B293 | DSM 44439 | Yes |
| *Nocardia coubleae* | OFN N12, CIP 108996 | DSM 44960 | Yes |
| *Nocardia harenae* | WS-26 | DSM 45095 | No |
| *Nocardia jiangsuensis* | KLBMP S0027 | DSM 101725 | Yes |
| *Nocardia neocaledoniensis* | SBH$_R$OA6 | DSM 44717 | Yes |
| *Rhodococcus biphenylivorans* | TG9 | DSM 102211 | Yes |
| *Rhodococcus imtechensis* | RKJ300(T) | DSM 45091 | Yes |
| *Rhodococcus phenolicus* | G2P | DSM 44812 | No |
| *Rhodococcus rhodochrous* | RPK1 | DSM 103064 | Yes |
| *Rhodococcus ruber* | C208 | DSM 45332 | Yes |
| *Rhodococcus erythropolis* | N9T-4 | NBRC 110906 | Yes |

**Table A5:** Antibiotic sensitivity of all actinomycete strains tested.

| Species | Kanamycin 100 µg/ml | Spectinomycin 100 µg/ml | Apramycin 50 µg/ml | Nourseothricin 100 µg/ml | Zeocin 100 µg/ml |
|---|---|---|---|---|---|
| *D. dagingensis* | - | - | - | - | - |
| *D. maris* | - | - | - | - | - |
| *D. natronolimnaea* | - | - | - | - | - |
| *D. psychralcaliphila* | - | - | - | - | - |
| *G. alkanivorans* | - | +/- | - | - | - |
| *G. paraffinivorans* | - | - | - | - | - |
| *G. polyisoprenivorans* | - | - | - | - | - |
| *N. coubleae* | + | + | - | - | - |
| *N. harenae* | - | - | - | - | - |
| *N. jiangsuensis* | - | - | - | - | - |
| *N. neocaledoniensis* | + | + | - | - | - |
| *R. biphenylivorans* | - | - | - | - | - |
| *R. imtechensis* | - | - | - | - | - |
| *R. phenolicus* | - | - | - | - | - |
| *R. rhodochrous* | - | - | - | - | - |
| *R. ruber* | - | - | +/- | - | - |
| *R. erythropolis* | + | - | - | - | +/- |

**Table A6**

| | |
|---|---|
| **AACG**GCCTATCGCTGTGAACAGGTGAGATTACGGAGAACGGGGCNNNNNNCCGTCCCTGTCGTGTGGNN**CAAT** | |
| PamiM WT -35 and -10 sequence: TTGTGGCCGTCCCTGTCGTGTGGTACAAT | PamiM 10 -35 and -10 sequence: TTGTGGCCGTCCCTGTCGTGTGGGACAAT |
| PamiM 70 -35 and -10 sequence: TCGACGCCGTCCCTGTCGTGTGGTACAAT | PamiM 2.5 -35 and -10 sequence: TTGGCACCGTCCCTGTCGTGTGGAACAAT |
| PamiM 25 -35 and -10 sequence: TGTCTTCCGTCCCTGTCGTGTGGTACAAT | PamiM 0 -35 and -10 sequence: ACTGGGCCGTCCCTGTCGTGTGGTTCAAT |

The complete annotated sequence of the Promoter P*amiM* T2A and the variable sequences that differentiate each promoter variant. Restriction overhangs are underlined and in bold. The -35 region is highlighted in yellow. The -10 region is highlighted in green.

**Table A7**

| | |
|---|---|
| **GGAG**AGCTGTCACCGGATGTGCTTTCCGGTCTGATGAGTCCGTGAGGACGAAACAGCCTCTACAAATAATTTTGTTTAAGGGCCCAAGTTCACTTA**ACTAAGGAA**TCAACAATGACCATGATTACGGATTCACTGGCCGTCGTTTTAGC**NNNNNNNNN**TTAAA**TATG** | |
| SD2 RhBCD 100: AGAAGGAGA | SD2 RhBCD 20: GCATTGCGG |
| SD2 RhBCD 80: AAAGGAGGT | SD2 RhBCD 10: TTTCCTCCA |
| SD2 RhBCD50: GGAGGCAGC | SD2 RhBCD 1: TCACGTCCC |

The complete annotated sequence of the previously reported BCD layout for *Rhodococcus*. RiboJ is highlighted in grey, the leader peptide 5´ UTR is highlighted in yellow with the SD1 sequence underlined and in bold, the leader peptide is highlighted in red with the SD2 region underlined and in bold, and the start codon is highlighted in cyan. The 4 nt cloning overhangs at the 5' and 3' ends are underlined and in bold. The six different 9 nt SD2 sequences corresponding to the different BCD translation initiation strengths are indicated. Note, RhBCD 80 corresponds to a canonical SD sequence and RhBCD 10 corresponds to a perfect mismatch.

**Table A8**

| | |
|---|---|
| **GGAG**AGCTGTCACCGGATGTGCTTTCCGGTCTGATGAGTCCGTGAGGACGAAACAGCCTCTACAAATAATTTTGTTTAAGGGCCCAAGTTCACTTAA**AAAGGAGAT**CAACAATGAAAGCAATTTTCGTACTGAAACATCTTAATCATGC**NNNNNNNNN**TTAAA**TATG** | |
| SD2 40k: AGAAGGAGA | SD2 0.75K: GGATTCTAG |
| SD2 25K: GGTAAGGAG | SD2 0.1K: TCAGGCCCC |
| SD2 8.5K: GGCGCGGTG | SD2 0.2K: TCACGTCCC |

The complete annotated sequence of the previously reported BCD layout for *E. coli*. RiboJ is highlighted in grey, the leader peptide 5´ UTR is highlighted in yellow with the SD1 sequence underlined and in bold, the leader peptide is highlighted in red with the SD2 region underlined and in bold, and the start codon is highlighted in cyan. The 4 nt cloning overhangs at the 5' and 3' ends are underlined and in bold. The 6 different 9 nt SD2 sequences corresponding to the different BCD translation initiation strengths are indicated.

**Table A9:** DNA sequences of all fluorescent protein and counter-selectable marker coding sequences. Restriction enzyme overhangs are underlined and in bold.

| Coding sequences |
|---|
| *codA* |
| **TATG**ATGGTGTCGAATAACGCTTTACAAACAATTATTAACGCCCGGTTACCAGGCGAAGAGGGGCTGTG<br>GCAGATTCATCTGCAGGACGGAAAAATCAGCGCCATTGATGCGCAATCCGGCGTGATGCCCATAACTGA<br>AAACAGCCTGGATGCCGAACAAGGTTTAGTTATACCGCCGTTTGTGGAGCCACATATTCACCTGGACAC<br>CACGCAAACCGCCGGACAACCGAACTGGAATCAGTCCGGCACGCTGTTTGAAGGCATTGAACGCTGGGC<br>CGAGCGCAAAGCGTTATTAACCCATGACGATGTGAAACAACGCGCATGGCAAACGCTGAAATGGCAGAT<br>TGCCAACGGCATTCAGCATGTGCGTACCCATGTCGATGTTTCGGATGCAACGCTAACTGCGCTGAAAGC<br>AATGCTGGAAGTGAAGCAGGAAGTCGCGCCGTGGATTGATCTGCAAATCGTCGCCTTCCCTCAGGAAGG<br>GATTTTGTCGTATCCCAACGGTGAAGCGTTGCTGGAAGAGGCGTTACGCTTAGGGGCAGATGTAGTGGG<br>GGCGATTCCGCATTTTGAATTTACCCGTGAATACGGCGTGGAGTCGCTGCATAAAACCTTCGCCCTGGC<br>GCAAAAATACGACCGTCTGATCGACGTTCACTGTGATGAGATCGATGACGAGCAGTCGCGCTTTGTCGA<br>AACCGTTGCTGCCCTGGCGCACCATGAAGGCATGGGCGCGCGAGTCACCGCCAGCCACACCACGGCAAT<br>GCACTCCTATAACGGGGCGTATACCTCACGCCTGTTCCGCTTGCTGAAAATGTCCGGTATTAACTTTGT<br>CGCCAACCCGCTGGTCAATATTCATCTGCAAGGACGTTTCGATACGTATCCAAAACGTCGCGGCATCAC<br>GCGCGTTAAAGAGATGCTGGAGTCCGGCATTAACGTCTGCTTTGGTCACGATGATGTCTTCGATCCGTG<br>GTATCCGCTGGGAACGGCGAATATGCTGCAAGTGCTGCATATGGGGCTGCATGTTTGCCAGTTGATGGG<br>CTACGGGCAGATTAACGATGGCCTGAATTTAATCACCCACCACAGCGCAAGGACGTTGAATTTGCAGGA<br>TTACGGCATTGCCGCCGGAAACAGCGCCAACCTGATTATCCTGCCGGCTGAAAATGGGTTTGATGCGCT<br>GCGCCGTCAGGTTCCGGTACGTTATTCGGTACGTGGCGGCAAGGTGATTGCCAGCACACAACCGGCACA<br>AACCACCGTATATCTGGAGCAGCCAGAAGCCATCGATTACAAACGCCTAGGGGGTGGCGGCGGTTC**GGC**<br>**T** |
| *upp* |
| **GCAG**GCGGTGGTGGCTCCTGCATGAAGATCGTGGAAGTCAAACACCCACTCGTCAAACACAAGCTGGGA<br>CTGATGCGTGAGCAAGATATCAGCACCAAGCGCTTTCGCGAACTCGCTTCCGAAGTGGGTAGCCTGCTG<br>ACTTACGAAGCGACCGCCGACCTCGAAACGGAAAAAGTAACTATCGAAGGCTGGAACGGCCCGGTAGAA<br>ATCGACCAGATCAAAGGTAAGAAAATTACCGTTGTGCCAATTCTGCGTGCGGGTCTTGGTATGATGGAC<br>GGTGTGCTGGAAAACGTTCCGAGCGCGCGCATCAGCGTTGTCGGTATGTACCGTAATGAAGAAACGCTG<br>GAGCCGGTACCGTACTTCCAGAAACTGGTTTCTAACATCGATGAGCGTATGGCGCTGATCGTTGACCCA<br>ATGCTGGCAACCGGTGGTTCCGTTATCGCGACCATCGACCTGCTGAAAAAAGCGGGCTGCAGCAGCATC<br>AAAGTTCTGGTGCTGGTAGCTGCGCCAGAAGGTATCGCTGCGCTGGAAAAAGCGCACCCGGACGTCGAA<br>CTGTATACCGCATCGATTGATCAGGGACTGAACGAGCACGGATACATTATTCCGGGCCTCGGCGATGCC<br>GGTGACAAAATCTTTGGTACGAAATGAGCGGGACTCTGGGGTTCGAAATGACCGACCAAGCGACGCCCA<br>ACCTGCCATCACGAGATTTCGATTCCACCGCCGCCTTCTATGAAAGGTTGGGCTTCGGAATCGTTTTCC<br>GGGACGCCGGCTGGATGATCCTCCAGCGCGGGGATCTCATGCTGGAGTT**GCTG** |
| *apr* |
| **GGCT**ATGCAATACGAATGGCGAAAAGCCGAGCTCATCGGTCAGCTTCTCAACCTTGGGGTTACCCCCGG<br>CGGTGTGCTGCTGGTCCACAGCTCCTTCCGTAGCGTCCGGCCCCTCGAAGATGGGCCACTTGGACTGAT<br>CGAGGCCCTGCGTGCTGCGCTGGGTCCGGGAGGGACGCTCGTCATGCCCTCGTGGTCAGGTCTGGACGA<br>CGAGCCGTTCGATCCTGCCACGTCGCCCGTTACACCGGACCTTGGAGTTGTCTCTGACACATTCTGGCG<br>CCTGCCAAATGTAAAGCGCAGCGCCCATCCATTTGCCTTTGCGGCAGCGGGGCCACAGGCAGAGCAGAT<br>CATCTCTGATCCATTGCCCCTGCCACCTCACTCGCCTGCAAGCCCGGTCGCCCGTGTCCATGAACTCGA<br>TGGGCAGGTACTTCTCCTCGGCGTGGGACACGATGCCAACACGACGCTGCATCTTGCCGAGTTGATGGC<br>AAAGGTTCCCTATGGGGTGCCGAGACACTGCACCATTCTTCAGGATGGCAAGTTGGTACGCGTCGATTA<br>TCTCGAGAATGACCACTGCTGTGAGCGCTTTGCCTTGGCGGACAGGTGGCTCAAGGAGAAGAGCCTTCA<br>GAAGGAAGGTCCAGTCGGTCATGCCTTTGCTCGGTTGATTCGCTCCCGCGACATTGTGGCGACAGCCCT<br>GGGTCAACTGGGCCGAGATCCGTTGATCTTCCTGCATCCGCCAGAGGCGGGATGCGAAGAATGCGATGC<br>CGCTCGCCAGTCGATTGGC**GCAG** |

| *spec* |
|---|
| **GGCT**ATGAGGGAAGCGGTGATCGCCGAAGTATCGACTCAACTATCAGAGGTAGTTGGCGTCATCGAGCG<br>CCATCTCGAACCGACGTTGCTGGCCGTACATTTGTACGGCTCCGCAGTGGATGGCGGCCTGAAGCCACA<br>CAGTGATATTGATTTGCTGGTTACGGTGACCGTAAGGCTTGATGAAACAACGCGGCGAGCTTTGATCAA<br>CGACCTTTTGGAAACTTCGGCTTCCCCTGGAGAGAGCGAGATTCTCCGCGCTGTAGAAGTCACCATTGT<br>TGTGCACGACGACATCATTCCGTGGCGTTATCCAGCTAAGCGCGAACTGCAATTTGGAGAATGGCAGCG<br>CAATGACATTCTTGCAGGTATCTTCGAGCCAGCCACGATCGACATTGATCTGGCTATCTTGCTGACAAA<br>AGCAAGAGAACATAGCGTTGCCTTGGTAGGTCCAGCGGCGGAGGAACTCTTTGATCCGGTTCCTGAACA<br>GGATCTATTTGAGGCGCTAAATGAAACCTTAACGCTATGGAACTCGCCGCCCGACTGGGCTGGCGATGA<br>GCGAAATGTAGTGCTTACGTTGTCCCGCATTTGGTACAGCGCAGTAACCGGCAAAATCGCGCCGAAGGA<br>TGTCGCTGCCGACTGGGCAATGGAGCGCCTGCCGGCCCAGTATCAGCCCGTCATACTTGAAGCTAGACA<br>GGCTTATCTTGGACAAGAAGAAGATCGCTTGGCCTCGCGCGCAGATCAGTTGGAAGAATTTGTCCACTA<br>CGTGAAAGGCGAGATCACCAAGGTAGTCGGCAAA**GCAG** |
| *zeo* |
| **GGCT**ATCAGCGGAGCTAATGGCGTCATGGCCAAGTTGACCAGTGCCGTTCCGGTGCTCACCGCGCGCGA<br>CGTCGCCGGAGCGGTCGAGTTCTGGACCGACCGGCTCGGGTTCTCCCGGGACTTCGTGGAGGACGACTT<br>CGCCGGTGTGGTCCGGGACGACGTGACCCTGTTCATCAGCGCGGTCCAGGACCAGGTGGTGCCGGACAA<br>CACCCTGGCCTGGGTGTGGGTGCGCGGCCTGGACGAGCTGTACGCCGAGTGGTCGGAGGTCGTGTCCAC<br>GAACTTCCGGGACGCCTCCGGGCCGGCCATGACCGAGATCGGCGAGCAGCCGTGGGGGCGGGAGTTCGC<br>CCTGCGCGACCCGGCCGGCAACTGCGTGCACTTCGTGGCCGAGGAGCAAGATCAGGGACCTGGA**GCAG** |
| E2-Crimson |
| **TATG**GACTCCACCGAGAACGTCATCAAGCCCTTCATGCGCTTCAAGGTCCACATGGAGGGCTCCGTCAA<br>CGGCCACGAGTTCGAGATCGAGGGCGTCGGCGAGGGCAAGCCCTACGAGGGCACCCAGACCGCCAAGCT<br>CCAGGTCACCAAGGGCGGCCCCCTCCCCTTCGCCTGGGACATCCTCTCCCCCCAGTTCTTCTACGGCTC<br>CAAGGCCTACATCAAGCACCCCGCCGACATCCCCGACTACCTCAAGCAGTCCTTCCCCGAGGGCTTCAA<br>GTGGGAGCGCGTCATGAACTTCGAGGACGGCGGCGTCGTCACCGTCACCCAGGACTCCTCCCTCCAGGA<br>CGGCACCCTCATCTACCACGTCAAGTTCATCGGCGTCAACTTCCCCTCCGACGGCCCCGTCATGCAGAA<br>GAAGACCCTCGGCTGGGAGCCCTCCACCGAGCGCAACTACCCCCGCGACGGCGTCCTCAAGGGCGAGAA<br>CCACATGGCCCTCAAGCTCAAGGGCGGCGGCCACTACCTCTGCGAGTTCAAGTCCATCTACATGGCCAA<br>GAAGCCCGTCAAGCTCCCCGGCTACCACTACGTCGACTACAAGCTCGACATCACCTCCCACAACGAGGA<br>CTACACCGTCGTCGAGCAGTACGAGCGCGCCGAGGCCCGCCACCACCTCTTCCAGTA**ATCC** |
| mCherry |
| **TATG**GTGTCCAAGGGCGAGGAGGACAACATGGCCATCATCAAGGAGTTCATGCGCTTCAAGGTCCACAT<br>GGAGGGCTCCGTCAACGGCCACGAGTTCGAGATCGAGGGCGAGGGCGAGGGCCGCCCCTACGAGGGCAC<br>CCAGACCGCCAAGCTCAAGGTCACCAAGGGCGGCCCCCTCCCCTTCGCCTGGGACATCCTCTCCCCCCA<br>GTTCATGTATGGCTCCAAGGCCTACGTCAAGCACCCCGCCGACATCCCCGACTACCTCAAGCTCTCCTT<br>CCCCGAGGGCTTCAAGTGGGAGCGCGTCATGAACTTCGAGGACGGCGGCGTCGTCACCGTCACCCAGGA<br>CTCCTCCCTCCAGGACGGCGAGTTCATCTACAAGGTCAAGCTCCGCGGCACCAACTTCCCCTCCGACGG<br>CCCCGTCATGCAGAAGAAGACCATGGGCTGGGAGGCCTCCTCCGAGCGCATGTATCCCGAGGACGGCGC<br>CCTCAAGGGCGAGATCAAGCAGCGCCTCAAGCTCAAGGACGGCGGCCACTACGACGCCGAGGTCAAGAC<br>CACCTACAAGGCCAAGAAGCCCGTCCAGCTCCCCGGCGCCTACAACGTCAACATCAAGCTCGACATCAC<br>CTCCCACAACGAGGACTACACCATCGTCGAGCAGTATGAGCGCGCCGAGGGCCGCCACTCCACCGGCGG<br>CATGGACGAGCTCTACAAGTA**ATCC** |
| mScarlet-I |
| **TATG**GTGTCCAAGGGCGAGGCCGTCATCAAGGAGTTCATGCGCTTCAAGGTCCACATGGAGGGCTCCAT<br>GAACGGCCACGAGTTCGAGATCGAGGGCGAGGGCGAGGGCCGCCCCTACGAGGGCACCCAGACCGCCAA<br>GCTCAAGGTCACCAAGGGCGGCCCCCTCCCCTTCTCCTGGGACATCCTCTCCCCCCAGTTCATGTATGG<br>CTCCCGCGCCTTCATCAAGCACCCCGCCGACATCCCCGACTACTACAAGCAGTCCTTCCCCGAGGGCTT<br>CAAGTGGGAGCGCGTCATGAACTTCGAGGACGGCGGCGCCGTCACCGTCACCCAGGACACCTCCCTCGA<br>GGACGGCACCCTCATCTACAAGGTCAAGCTCCGCGGCACCAACTTCCCCCCCGACGGCCCCGTCATGCA<br>GAAGAAGACCATGGGCTGGGAGGCCTCCACCGAGCGCCTCTACCCCGAGGACGGCGTCCTCAAGGGCGA |

| |
|---|
| CATCAAGATGGCCCTCCGCCTCAAGGACGGCGGCCGCTACCTCGCCGACTTCAAGACCACCTACAAGGC<br>CAAGAAGCCCGTCCAGATGCCCGGCGCCTACAACGTCGACCGCAAGCTCGACATCACCTCCCACAACGA<br>GGACTACACCGTCGTCGAGCAGTATGAGCGCTCCGAGGGCCGCCACTCCACCGGCGGCATGGACGAGCT<br>CTACAAGTA**ATCC** |
| BFP-Bluebonnet |
| **TATG**TCCGAGGAGCTTATCAAGGAAAATATGCATATGAAACTTTACATGGAAGGATGGGTTGACAATCA<br>CCACTTTAAGTGTACAGCAGAAGGTGAAGGCAAGCCGTATGAGGGTACTCAGACGATGCGTATCAAGGT<br>GGTAGAGGGCGGTCCGTTACCCTTTGCTTTTGATATTTTGGCAACTTCCTTCTTGTATGGCTCAAAAAC<br>ATTTATTGACCATACACAGGGCATTCCTGATTTTTTCAAGCAATCTTTTCCAGAAGGCTTTACTTGGGA<br>GCGTGTCACTACTTATGAGGATGGTGGAGTCCTGACTGCTACGCAGGACACAAGCCTGCAAGACGGCGT<br>CCTTATTTATAATGTAAAGATTCGCGGCACCGATTTTCCAAGCAATGGACCTGTTATGCAAAAAAAAAC<br>CCTTGGTTGGGAAGCGTTTACTGAGACTTTATACCCCGCCGGCGGTGGCTTGGAAGGACGCAATGACAT<br>GGCACTTAAATTAGTAGGTGGATCGCATTTGATCGCGACAGCAAAGACTACCTACCGTTCTAAGAAGCC<br>TGCGAAGAACCTGAAAATGCCAGGTGTCTATTACGTCGATTACTTGTTGGAACGTATTAAGGAAGCAAA<br>TGATGAAACCTACGTTGAGCAGCATGAAGTGGCAGTGGCTCGTTATTCAGACTTGCCTTCAAAACTTGG<br>TCATAAGTTAAATTA**ATCC** |
